# Screening of HLA-A restricted T cell epitopes of SARS-CoV-2 and induction of CD8^+^ T cell responses in HLA-A transgenic mice

**DOI:** 10.1101/2021.04.01.438020

**Authors:** Xiaoxiao Jin, Yan Ding, Shihui Sun, Xinyi Wang, Zining Zhou, Xiaotao Liu, Miaomiao Li, Xian Chen, Anran Shen, Yandan Wu, Bicheng Liu, Jianqiong Zhang, Jian Li, Yi Yang, Haibo Qiu, Chuanlai Shen, Yuxian He, Guangyu Zhao

## Abstract

While SARS-CoV-2-specific T cells have been characterized to play essential roles in host immune protection in COVID-19 patients, few researches focus on the functional validation of T cell epitopes and development of vaccines inducing specific T cell responses. In this study, 120 CD8^+^ T cell epitopes from E, M, N, S and RdRp proteins were validated. Among them, 110 epitopes have not been reported previously; 110, 15, 6, 14 and 12 epitopes were highly homologous with SARS-CoV, OC43, NL63, HKU1, and 229E, respectively; 4 epitopes from S protein displayed one amino acid distinct from the current variants of SARS-CoV-2. Thirty-one epitopes restricted by HLA-A2 molecule were used to generate peptide cocktail vaccines in combination with Poly(I:C), R848 or polylactic-co-glycolic acid nanoparticles, which elicited robust specific CD8^+^ T cell responses in wild-type and HLA-A2/DR1 transgenic mice. Seven of the 31 epitopes were found to be cross-presented by HLA-A2 and H-2K/D^b^ molecules. Unlike previous researches, this study established a modified cell co-culture system of DC-peptide-PBL using healthy donor’s PBMCs to validate the CD8^+^ T cell epitope on-silicon predicted; provided a library of CD8^+^ T cell epitopes restricted by a series of high-frequency HLA-A allotypes which covering broad Asian populations; identified the HLA-A cross-restrictions of these CD8^+^ T cell epitopes using competitive binding experiments with HMy2.CIR cell lines expressing indicated HLA-A molecules; and initially confirmed the *in vivo* feasibility of 9 or 10-mer peptide cocktail vaccines of SARS-CoV2. These data will facilitate the development of vaccines inducing antiviral CD8^+^ T cell responses.

## Introduction

The highly contagious COVID-19 has spread worldwide at an unprecedentedly quick speed since its first identification at December 2019, leading to an ongoing global pandemic^1^. As 22 of April, 2021, there have been more than 143 million confirmed cases and over 3.0 million deaths. Although relentless efforts have been paid in effective vaccine race, there are still many risks for a long-term immune protection^2^, since host immunity to Severe Acute Respiratory Syndrome Coronavirus 2 (SARS-CoV-2) has not been fully understood. Currently, most vaccines focus on the induction of neutralizing antibodies against spike (S) protein of SARS-CoV-2 ^3, 4^, which can block the virus from entering and infecting human cells, helping the immune system to clear the virus and to prevent future infections^5^. However, researchers have found that circulating antibodies to SARS-Cov-2 declined rapidly and persisted only around seven months. And, certain patients who are asymptomatic or mildly symptomatic do not have detectable neutralizing antibodies^6^. In addition, growing incidence of COVID-19 re-infections emerges since the first reported re-infection in August, 2020. This suggests that neutralizing antibodies cannot offer long-term protection. Furthermore, SARS-CoV-2 constantly undergoes mutations as it spread from person to person. Therefore, more studies are needed to determine whether the current vaccines will be still effective against the virus quasispecies.

It has been studies that T cells, especially CD8^+^ T cells, also play a critical role in the defense of many viral infections^7^. Increasing evidences indicate that T cell responses are important in the immune response against SARS-CoV-2, and may mediate long-term protection^8–12^. The specific T cells showed a highly activated cytotoxic phenotype in the acute phase that correlated with various clinical markers of disease severity, whereas the convalescent-phase specific T cells were multifunctional and showed a stem cell-like memory phenotype in the mild and asymptomatic patients^12–19^. Moreover, intensive expansion of highly cytotoxic T cells was associated with convalescence in moderate patients^19^. SARS-CoV-2-specifific T cells were detectable in antibody-seronegative exposed family members and convalescent individuals with a history of asymptomatic and mild COVID-19^12^. As a result, T cell immunity would be critical in the pathogenesis and immune protection mechanism of COVID-19, thus providing a potential way to develop long-term effective vaccines and treatments^20^. However, thus far limited information is available about the vaccines inducing T cell immune protections against SARS-CoV-2. Very recently, SARS-CoV-2 T cell epitopes restricted by H-2^d^ and H-2^b^ molecules were identified, and three of which were used to generate the venezuelan equine encephalitis replicon particles (VRP) expressing single T cell epitope^21^. This VRP vaccine induced robust CD4^+^ or CD8^+^ T cell responses, which mediated more rapid viral clearance than neutralizing antibodies and decreased the extent of lung pathological changes in Ad5-ACE2-tranduced and SARS-CoV-2-infected mice^21^, indicating the potential of T cell epitope vaccine. In addition, HLA-DR-restricted peptides cocktail vaccine from Tubingen University of Germany has been enrolled in phase I clinical trial (NCT04546841). Whether the 9- or 10-mer peptides cocktail restricted by HLA class I molecule can induce SARS-CoV-2 sepcific CD8^+^ T cell responses *in vivo* remains unknown.

Identification of T cell epitopes in SARS-CoV-2 proteins can contribute greatly to the development of T cell epitope vaccines and precise evaluation of host cellular immunity. However, most studies have utilized pools of predicted or overlapping peptides spanning the sequences of different SARS-CoV-2 proteins ^22–28^. The functionally validated T cell epitopes are still limited and have thus far come from only a few laboratories^10, 11, 29–32^. These T cell epitopes are presented only by several HLA class I or II molecules and the precise HLA restrictions of each epitope needs to be further defined.

To screen more CD8^+^ T cell epitopes which cover more predominant HLA-A allotypes and more SARS-CoV-2 proteins, here we dedicated to the mapping of HLA-A-restricted epitopes from envelope proteins (E), membrane protein (M), nucleocapsid protein (N), spike glycoprotein (S) and RNA-dependent RNA polymerase (RdRp) of SARS-CoV-2. Four hundred and nine epitopes, which restricted by nine high-frequency HLA-A allotypes, were on-silicon predicted and selected. Then the immunogenicity and HLA-A cross-restrictions of 120 epitopes were validated by DC-peptide-PBL co-culture experiments using healthy donors’ PBMCs and competitive binding experiments with HMy2. CIR cell lines expressing indicated HLA-A molecules. Thirty-one epitope peptides restricted by HLA-A2 molecule were used to generate the 9- or 10-mer peptide cocktail vaccines in combination with Poly(I:C), R848 or polylactic-co-glycolic acid nanoparticles (PLGA-NPs), which induced robust SARS-CoV-2 specific CD8^+^ T cell responses in HLA-A2/DR1 transgenic mice and wild-type mice.

## Results

### 1. HLA-A restricted 409 T cell epitopes were on-silicon predicted and selected from SARS-CoV-2 proteins

SARS-CoV-2 T cell epitopes, which restricted by nine high-frequency HLA-A allotypes (HLA-A*02:01, A*11:01, A*24:02, A*02:06, A*02:07, A*33:03, A*30:01, A*02:03, or A*11:02) with the total gene frequency of around 87% in Chinese population (http://www.allelefrequencies.net), were predicted from four structural protein (E, M, N, S) and one non-structural protein (RdRP) using five epitope predication tools (IEDB-ANN, IEDB-SMM, SYFPEITHI, EPIJEN, NetMHC and ConvMHC). For each HLA-A molecule and each protein, one to twenty 9- or 10-mer peptides with the highest score (highest affinity) were selected as candidate epitopes. A total of 409 peptides restricted by indicated HLA-A molecules were finally selected as candidate epitopes with the number of 45, 63, 71, 130, and 100 from E, M, N, S and RdRp proteins, respectively. Among them, 139 indicated epitopes are common or highly homologous epitopes cross-presented by several HLA-A molecules, thus finally only 270 indicated candidate epitope peptides (9-mer or 10-mer) were synthesized for further identification (Table S1).

### 2. Immunogenicity of 120 candidate epitopes was validated by DC-peptide-PBL co-culture experiments

In order to validate the immunogenicity of candidate epitopes, PBMCs from healthy blood donors were collected and HLA-A alleles were identified. DCs were induced for 7 days and then coincubated with candidate epitope peptides and autologous PBL for 14 days. Cells were harvested and stimulated by corresponding candidate peptides for another 16 hours followed by IFN-γ intracellular staining (ICS) and flow cytometry. In some DC-peptide-PBL co-culture wells, the autologous PBLs were prelabeled with CFSE. After 14-day co-cultures, the cells were harvested followed by flow cytometry to detect the proliferation percentage of CD8^+^ T cell. When the frequency of IFN-γ^+^ T cells in CD3^+^/CD8^+^ T cell population increased by more than 100% compared with the negative control or proliferation percentage of CD8^+^ T cells in CD3^+^/CD8^+^ T cell population increased by more than 20% compared with the negative control, the candidate epitope peptide in the co-culture well was identified as positive peptide with immunogenicity.

To evaluate whether this procedure is sensitive for identification of peptide immunogenicity, several reference peptides were tested using this DC-peptide-PBL co-culture procedure. These HLA-A restricted T cell epitopes derived from hepatocellular carcinoma (HCC)-associated tumor antigens (HCC 1-1, HCC 1-2, HCC 5-3, HCC 5-4, HCC 5-5) or from hepatitis B virus antigens (HBV 111, HBV 118) have been validated as real-world epitopes previously in-house by using HCC patients’ PBMCs or chronic hepatitis patient’s PBMCs (manuscript submitted). They can effectively stimulate the patient’s fresh PBMCs to produce IFN-γ in *ex vivo* 20-hour co-culture as detected by ELISPOT assay. Here, they were also defined as positive peptides in DC-peptide-PBL co-cultures using healthy donor’s PBMCs as detected by intracellular IFN-γ staining and CFSE proliferation analyses, meanwhile the weak positive reference peptides (HCC 1-1, HCC 1-2) were defined as negative peptides here (Figure 1).

**Figure 1:**
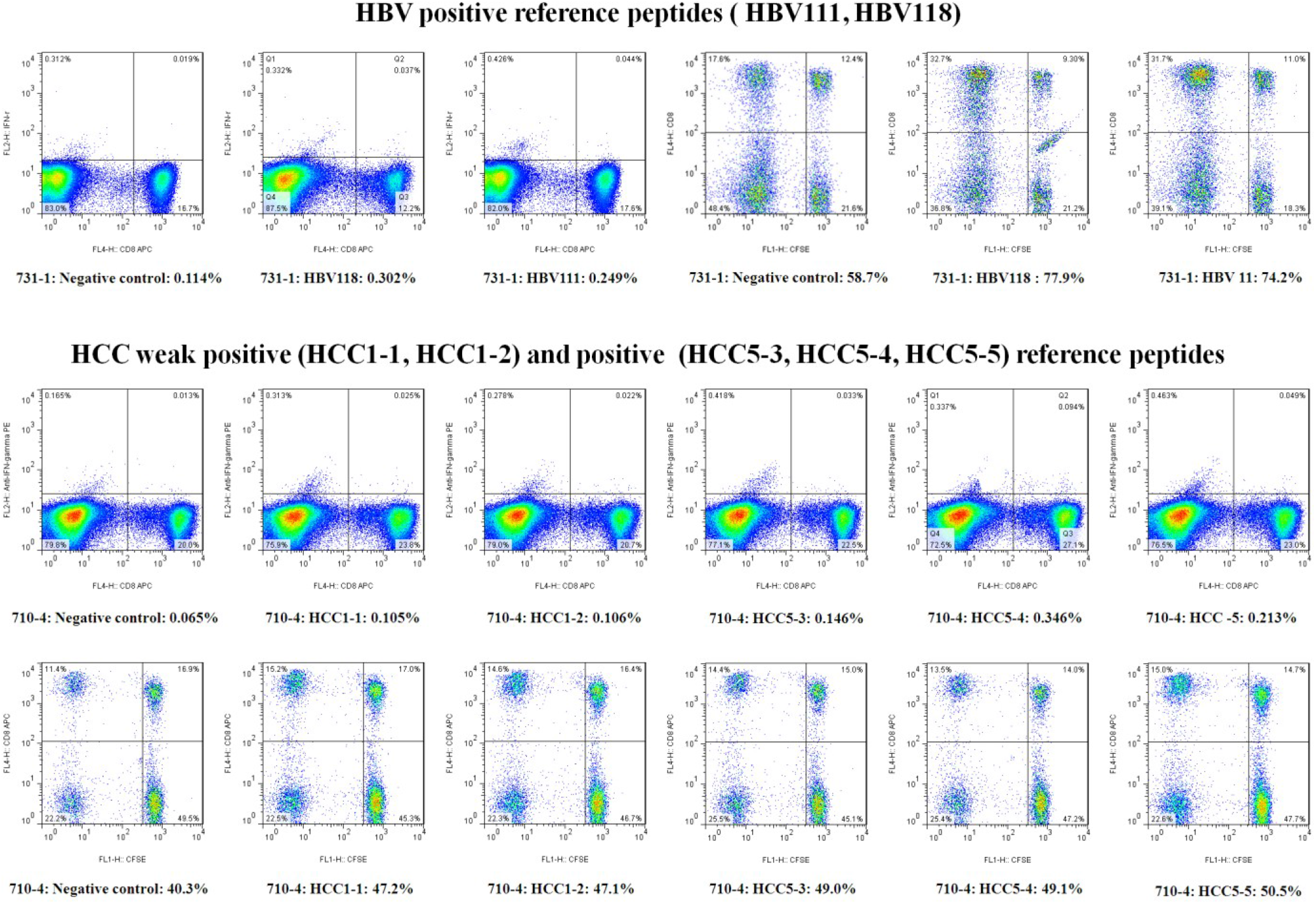
Reference epitope peptides were tested in the DC-peptide-PBL co-culture system. The HLA-A restricted HCC 1-1, HCC 1-2, HCC 5-3, HCC 5-4, HCC 5-5, HBV 111 and HBV 118 peptides, which have been validated as real-world epitopes previously in-house by using HCC patients’ or chronic hepatitis patient’s PBMCs and ELISPOT assay, were co-cultured with DC and PBLs from healthy donor’s PBMCs for 14 days. As detected by both IFN-γ ICS and CFSE proliferation analyses, the weak positive reference peptides (HCC 1-1 and HCC 1-2, SFUs/2×10^5^ PBMCs <10) were defined as negative peptides while other positive reference peptides (SFUs/2×10^5^ PBMCs >15) were identified as immunogenic epitope peptides. The frequency of IFN-γ^+^ T cells in CD3^+^/CD8^+^ T cell population increased by more than 100% and proliferation percentage of CD8^+^ T cells in CD3^+^/CD8^+^ T cell population increased by more than 20%, as compared with the no peptide negative control.

After the PBMC samples from 156 healthy donors were tested, a total of 120 candidate epitope peptides of SARS-CoV-2 have been defined as antigenic T cell epitopes by using the DC-peptide-PBL procedure, which indicating that they can elicit naive peptide-specific CD8^+^ T cells to activate and produce IFN-γ or proliferate after 14 days co-stimulation. Some candidate epitope peptides have been validated in several healthy donor’ PBMCs. 110 of 120 positive epitopes have not been reported previously. The detailed data of all positive peptides were summarized in Table 1. Of them, the number of epitopes derived from E (75aa), M (222aa), N (419aa), S (1273aa) and RdRp (932aa) proteins is 18, 27, 12, 36, and 27, respectively, with a relatively bias distribution. The density of CD8^+^ T cell epitopes per 10 aa is 2.40, 1.216, 0.286, 0.283, and 0.290, respectively. Their sequence homology between SARS-CoV-2 and other HCoVs were aligned and exhibited in Table S2. Of the 120 validated SARS-CoV-2 CD8^+^ T cell epitopes, 110, 15, 6, 14 and 12 epitopes were highly homologous (0-2 amino acids deviation) with SARS-CoV, OC43, NL63, HKU1,and 229E, respectively. The common epitopes with common-cold HCoVs mainly locate in RdRp protein (39/47). In addition, 4 epitopes displayed one amino acid distinct from the current mutant variants, such as D50 and D82 for B.1.1.7, D82 for B.1.351 and P.1, D78 for B.1.617, and D53 for Denmark Variant (Table S2).

**Table 1:** 120 SARS-CoV-2 T cell epitopes validated by DC-peptide-PBL co-stimulation experiments.

| Epitope | On-silicon<br>HLA-A<br>restriction | Blood donor 1 |  |  | Blood donor 2 |  |  | Blood donor 3 |  |  |
| --- | --- | --- | --- | --- | --- | --- | --- | --- | --- | --- |
|  |  | A allele | Method | Enhance% | A allele | Method | Enhance% | A allele | Method | Enhance% |
| A1 | 0201, 0207, 0206, 2402 | 0201/6843 | CFSE | 24.4% | 3101/0207 | IFN-γ | 130.0% | 1101/0201 | IFN-γ | 120.7% |
| A3 | 0201, 0203, 1101, 1102 | 0201/2402 | IFN-γ | 314.2% | | | | | | |
| A4 | 0201, 0207, 0203 | 0201/3303 | IFN-γ | 116.2% | 6801/3303 | IFN-γ | 436.6% | 2402/0207 | IFN-γ | 137.3% |
| A5 | 0201, 0203 | 0201/3303 | IFN-γ | 589.2% | 0201/3001 | IFN-γ | 650.2% | 6801/3303 | IFN-γ | 728.8% |
| A6 | 0207, 0206 | 0206/0207 | IFN-γ | 198.5% | 1101/0206 | IFN-γ | 263.0% | | | |
| A7 | 0207 | 0206/0207 | IFN-γ | 253.8% | | | | | | |
| A9 | 0206 | 0206/1101 | IFN-γ | 163.2% | 0206/0203 | IFN-γ | 131.7% | 1101/0206 | IFN-γ | 666.0% |
| A10 | 0203 | 0203/3101 | IFN-γ | 109.1% | | | | | | |
| A12 | 1101, 1102, 3303 | 1101/3001 | CFSE | 22.1% | 0201/1101 | IFN-γ | 158.5% | 3303/1101 | IFN-γ | 263.0% |
| A16 | 1101, 3303 | 0201/1101 | CFSE | 40.8% |  |  |  |  |  |  |
| A18 | 1102, 3303, 3001 | 1101/3101 | IFN-γ | 119.1% | | | | | | |
| A19 | 1102 | 1101/3101 | IFN-γ | 102.9% | 1101/2402 | CFSE | 22.3% | | | |
| A20 | 2402 | 2402/2601 | IFN-γ | 543.9% | 2402/2402 | IFN-γ | 146.1% | | | |
| A21 | 2402 | 2402/2601 | IFN-γ | 1154.4% | 2402/2402 | IFN-γ | 153.2% | | | |
| A22 | 2402 | 2402/2601 | IFN-γ | 378.9% | | | | | | |
| A23 | 2402 | 2402/2601 | IFN-γ | 303.5% | 2402/2402 | IFN-γ | 135.5% | | | |
| A25 | 3001 | 1101/3001 | CFSE | 129.2% |  |  |  |  |  |  |
| A26 | 3001 | 1101/3001 | CFSE | 39.5% | 1101/3001 | IFN-γ | 291.0% | | | |
| B1 | 0201, 0206, 0203 | 0201/3303 | IFN-γ | 110.8% | 0201/3001 | IFN-γ | 222.9% | 6801/3303 | IFN-γ | 398.0% |
| B2 | 0201, 0206, 0203, 0207 | 0201/0101 | IFN-γ | 128.1% | | | | | | |
| B3 | 0201, 0207, 0206 | 0201/3001 | IFN-γ | 407.3% | | | | | | |
| B4 | 0201, 0206 | 0201/3001 | IFN-γ | 216.6% | | | | | | |
| B6 | 0201, 0207, 0203 | 0203/0207 | IFN-γ | 242.1% | 0201/1101 | CFSE | 35.7% | 2402/0207 | IFN-γ | 164.8% |
| B10 | 0207 | 0207/0206 | IFN-γ | 888.7% | 3101/0207 | IFN-γ | 120.0% | 2402/0207 | IFN-γ | 123.8% |
| B11 | 0206 | 0206/1101 | IFN-γ | 154.5% | 0206/3201 | CFSE | 25.8% | 1101/0206 | IFN-γ | 307.0% |
| B12 | 0206 | 1101/0206 | CFSE | 22.5% | 1101/0206 | IFN-γ | 104.0% | | | |
| B15 | 0203 | 0203/3101 | IFN-γ | 416.5% | | | | | | |
| B16 | 0203 | 0203/3001 | CFSE | 24.7% |  |  |  |  |  |  |
| B17 | 0203 | 0203/3101 | IFN-γ | 122.7% | | | | | | |
| B18 | 1101, 1102 | 1101/3001 | CFSE | 79.0% |  |  |  |  |  |  |
| B20 | 1101, 1102, 3303 | 0201/1101 | CFSE | 33.4% |  |  |  |  |  |  |
| B21 | 1101, 1102 | 0201/1101 | CFSE | 55.4% |  |  |  |  |  |  |
| B23 | 1101, 1102, 3303 | 3303/1101 | IFN-γ | 431.5% | | | | | | |
| B26 | 2402 | 2402/2402 | IFN-γ | 214.5% | 2402/2601 | IFN-γ | 277.2% | | | |
| B28 | 2402 | 2402/3303 | IFN-γ | 214.5% | | | | | | |
| B29 | 2402 | 2402/3303 | IFN-γ | 500.0% | 2402/2402 | IFN-γ | 341.8% | 6801/3303 | IFN-γ | 142.5% |
| B30 | 2402 | 2402/3303 | IFN-γ | 362.7% | | | | | | |
| B31 | 2402 | 2402/2402 | IFN-γ | 129.0% | 2402/2402 | IFN-γ | 139.0% | | | |
| B34 | 3303 | 3303/3001 | IFN-γ | 102.1% | | | | | | |
| B35 | 3303 | 3303/1101 | IFN-γ | 148.2% | | | | | | |
| B36 | 3001 | 1101/3001 | CFSE | 54.9% | 1101/3001 | IFN-γ | 374.0% | | | |
| B37 | 3001 | 1101/3001 | CFSE | 34.9% | 3001/2402 | IFN-γ | 110.5% | 1101/3001 | IFN-γ | 191.0% |
| B38 | 3001 | 2402/3001 | IFN-γ | 186.8% | | | | | | |
| B40 | 3001 | 1101/3001 | IFN-γ | 140.0% | | | | | | |
| B41 | 3001 | 1101/3001 | IFN-γ | 425.0% | | | | | | |
| C1 | 0201, 0206, 0203 | 0201/1101 | CFSE | 30.0% |  |  |  |  |  |  |
| C3 | 0201, 0206, 0203 | 0201/2402 | IFN-γ | 270.9% | | | | | | |
| C10 | 0206 | 0206/1101 | CFSE | 25.9% |  |  |  |  |  |  |
| C12 | 0206 | 0206/3201 | CFSE | 20.8% |  |  |  |  |  |  |
| C16 | 0206 | 0206/1101 | IFN-γ | 187.7% | | | | | | |
| C17 | 0203 | 0203/3101 | IFN-γ | 157.4% | | | | | | |
| C27 | 1101 | 1101/1101 | CFSE | 40.1% |  |  |  |  |  |  |
| C35 | 2402 | 2402/2601 | IFN-γ | 144.0% | | | | | | |
| C45 | 3303 | 1101/3303 | IFN-γ | 181.1% | | | | | | |
| C46 | 3303 | 1101/3303 | IFN-γ | 100.0% | | | | | | |
| C47 | 3303 | 1101/0101 | IFN-γ | 223.8% | | | | | | |
| C49 | 1102 | 0201/1101 | IFN-γ | 1663.3% | | | | | | |
| D2 | 0201, 0206, 0203 | 0201/1101 | IFN-γ | 122.3% | | | | | | |
| D5 | 0207, 0206, 1102 | 0206/1102 | IFN-γ | 479.2% | | | | | | |
| D6 | 0201, 0207 | 0201/2402 | CFSE | 105.5% | 0201/0101 | IFN-γ | 106.9% | | | |
| D12 | 0201, 0207, 0206 | 0201/0201 | CFSE | 23.4% |  |  |  |  |  |  |
| D13 | 0201, 0203 | 0201/2402 | IFN-γ | 1001.3% | | | | | | |
| D17 | 0207, 0203 | 0206/0207 | IFN-γ | 112.3% | | | | | | |
| D26 | 0206 | 0201/0207 | CFSE | 29.9% |  |  |  |  |  |  |
| D30 | 0203 | 0203/0206 | CFSE | 34.6% | 0203/0207 | IFN-γ | 145.3% | | | |
| D31 | 0203 | 0203/0206 | CFSE | 33.5% |  |  |  |  |  |  |
| D32 | 0203 | 0203/0206 | CFSE | 33.3% |  |  |  |  |  |  |
| D33 | 0203, 0206 | 0203/0206 | CFSE | 28.0% |  |  |  |  |  |  |
| D34 | 1101/1102 | 1101/0101 | IFN-γ | 117.2% | | | | | | |
| D38 | 1101, 1102 | 1101/0101 | IFN-γ | 105.9% | | | | | | |
| D40 | 1101, 1102, 3001 | 1101/3303 | IFN-γ | 100.0% | | | | | | |
| D41 | 1101 | 1101/0101 | IFN-γ | 268.0% | | | | | | |
| D42 | 1101, 0206, 0201 | 1101/0101 | IFN-γ | 128.4% | 1101/1101 | CFSE | 47.6% | | | |
| D46 | 1102 | 0201/1101 | IFN-γ | 458.6% | | | | | | |
| D47 | 1102 | 0201/1101 | IFN-γ | 179.9% | | | | | | |
| D48 | 1102, 3303 | 0201/1101 | IFN-γ | 166.9% | 3303/1101 | IFN-γ | 117.8% | | | |
| D50 | 1102 | 0201/1101 | IFN-γ | 242.0% | | | | | | |
| D52 | 2402 | 2402/3001 | IFN-γ | 241.7% | | | | | | |
| D53 | 2402, 3303 | 2402/3001 | IFN-γ | 460.8% | 3303/1101 | IFN-γ | 275.5% | | | |
| D55 | 2402 | 2402/0207 | IFN-γ | 110.2% | | | | | | |
| D56 | 2402 | 2402/2601 | IFN-γ | 266.9% | | | | | | |
| D62 | 2402 | 2402/3001 | IFN-γ | 182.4% | | | | | | |
| D64 | 2402 | 0201/2402 | IFN-γ | 144.9% | | | | | | |
| D65 | 3001 | 0101/3001 | CFSE | 20.8% |  |  |  |  |  |  |
| D71 | 3001 | 0301/3001 | IFN-γ | 132.9% | | | | | | |
| D72 | 3001, 1102, 0201, 0207 | 3303/1101 | IFN-γ | 157.7% | | | | | | |
| D76 | 3303 | 1101/0101 | IFN-γ | 346.7% | | | | | | |
| D77 | 3303, 1102 | 3303/1101 | IFN-γ | 109.5% | 3303/0203 | IFN-γ | 102.9% | | | |
| D78 | 3303, 0203, 0206 | 3303/1101 | IFN-γ | 101.4% | | | | | | |
| D79 | 3303, 1101, 0207, 2402 | 3303/1101 | IFN-γ | 175.8% | | | | | | |
| D80 | 3303, 1102, 0201, 0207 | 3303/1101 | IFN-γ | 148.2% | | | | | | |
| D81 | 3303, 1102, 0203 | 3303/1101 | IFN-γ | 149.6% | | | | | | |
| D82 | 3303, 1102 | 3303/1101 | IFN-γ | 101.7% | | | | | | |
| R4 | 0201 | 0201/1101 | IFN-γ | 107.8% | 0201/3201 | CFSE | 47.3% | | | |
| R5 | 0201, 0203 | 0201/3201 | CFSE | 107.8% | 0201/0101 | IFN-γ | 117.5% | | | |
| R6 | 0201, 0207, 0206, 0203 | 0201/3201 | CFSE | 49.3% |  |  |  |  |  |  |
| R8 | 0201 | 0201/0101 | IFN-γ | 113.2% | | | | | | |
| R9 | 0201, 0206, 0203 | 0201/0101 | IFN-γ | 107.0% | | | | | | |
| R10 | 0201 | 0201/2402 | IFN-γ | 104.3% | | | | | | |
| R11 | 0201 | 0201/3001 | IFN-γ | 203.6% | 0201/1101 | IFN-γ | 104.4% | | | |
| R12 | 0201 | 0201/3001 | IFN-γ | 147.0% | 0201/0101 | IFN-γ | 188.6% | | | |
| R13 | 0201 | 0201/3001 | IFN-γ | 147.0% | 0201/1101 | IFN-γ | 126.9% | | | |
| R14 | 0201, 0203 | 0201/0101 | IFN-γ | 133.3% | | | | | | |
| R15 | 0201 | 0201/2402 | IFN-γ | 218.4% | | | | | | |
| R17 | 0207 | 0207/0206 | IFN-γ | 103.0% | | | | | | |
| R23 | 0206 | 2402/0207 | IFN-γ | 109.4% | | | | | | |
| R24 | 0203 | 0203/0206 | CFSE | 60.8% |  |  |  |  |  |  |
| R30 | 1101 | 1101/3001 | CFSE | 110.2% | 0201/1101 | CFSE | 38.7% |  |  |  |
| R32 | 1101/1102 | 0201/1101 | CFSE | 29.3% |  |  |  |  |  |  |
| R34 | 1101, 3001 | 0201/1101 | CFSE | 20.0% | 1101/3101 | IFN-γ | 122.1% | | | |
| R35 | 1101, 1102 | 1101/3101 | CFSE | 24.9% |  |  |  |  |  |  |
| R38 | 1101, 3303 | 0201/1101 | CFSE | 50.2% |  |  |  |  |  |  |
| R39 | 1101, 1102, 3001 | 0201/1101 | CFSE | 54.3% |  |  |  |  |  |  |
| R40 | 1101, 3001 | 0201/1101 | CFSE | 39.7% | 0201/1101 | CFSE | 19.5% |  |  |  |
| R41 | 1101, 3303 | 0201/1101 | CFSE | 53.3% |  |  |  |  |  |  |
| R42 | 1101 | 0201/1101 | CFSE | 21.0% |  |  |  |  |  |  |
| R43 | 1101, 3303 | 1101/3001 | CFSE | 41.9% | 0201/1101 | CFSE | 34.5% |  |  |  |
| R44 | 1101, 1102 | 0201/1101 | CFSE | 37.6% | 0201/1101 | CFSE | 24.8% |  |  |  |
| R47 | 2402 | 2402/3303 | IFN-γ | 339.8% | | | | | | |
| R48 | 2402 | 2402/3303 | IFN-γ | 309.6% | | | | | | |

Note 1: **CFSE**: After DC-peptide-CFSE-prelabeled PBLs co-cultures, the proliferation percentage of CD8<sup>+</sup> T cells in CD3<sup>+</sup>/CD8<sup>+</sup> population was analyzed according to the reduction of CFSE-staining brightness. **IFN- $\gamma$** : After DC-peptide-PBLs co-cultures, the frequency of IFN-γ <sup>+</sup>/CD8<sup>+</sup> T cells in CD3<sup>+</sup>/CD8<sup>+</sup> population was analyzed by flow cytometry. **Enhance%**: The increased percentage when the frequency of IFN-γ <sup>+</sup>/CD8<sup>+</sup> T cells or proliferation percentage of CD8<sup>+</sup> T cells in the DC-peptide-PBL co-culture wells was compared with that in the DC-PBL co-culture well without peptide. Note 2: The previously reported epitopes include C1, C35, C49, D34, D46, D47, D48, D52, D53 and R32.

Figure S1 showed the phenotypes of mature DC as verified by flow cytometry. Figure 2 presented the IFN-γ ICS flow plots and CD8^+^ T cell proliferation flow plots of partial positive peptides. All flow plots of 120 positive peptides were displayed in Figure S2 and S3.

**Figure 2:**
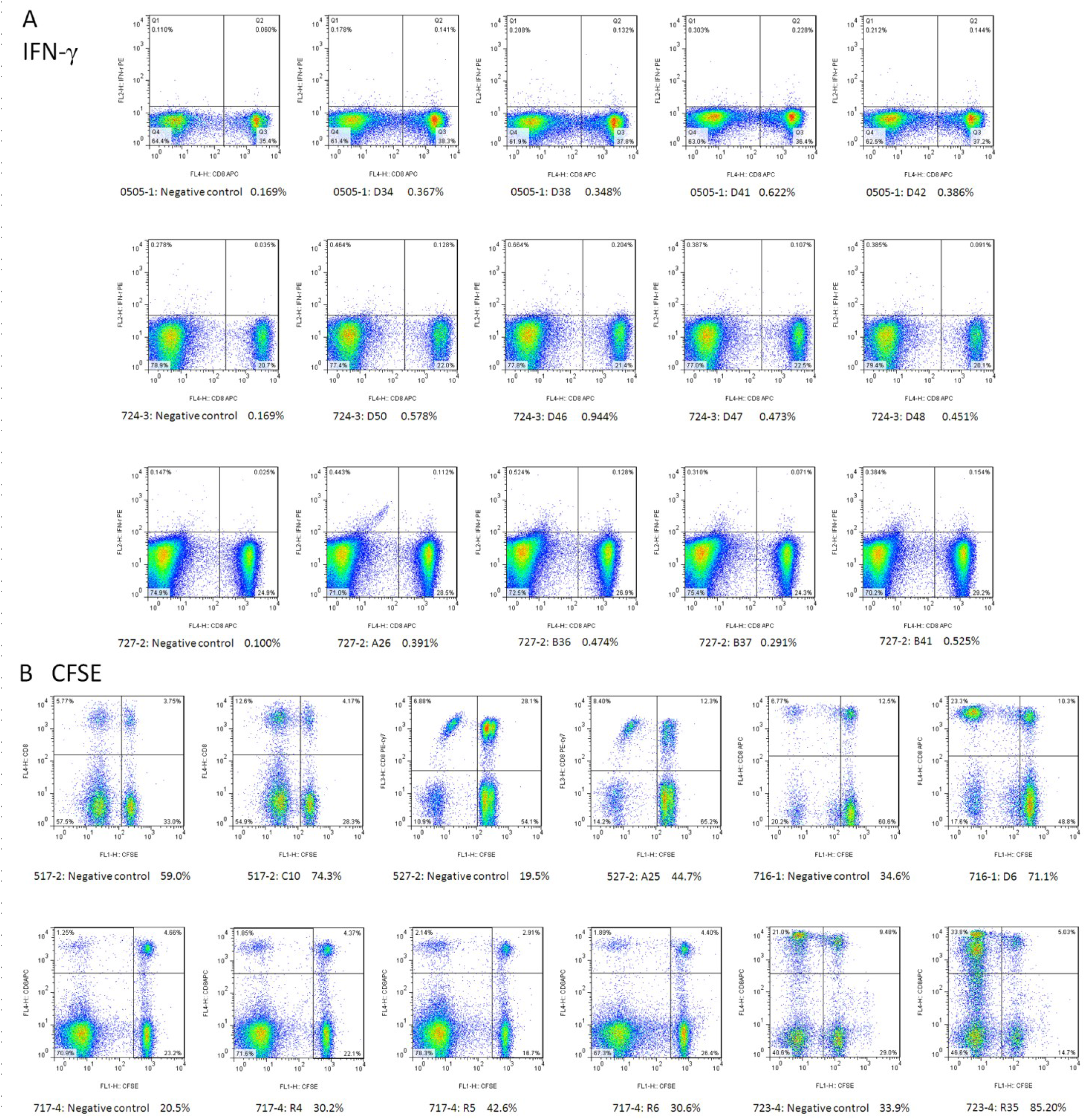
Immunogenicity of candidate epitopes was validated by DC-peptide-PBL co-culture experiments. DCs were induced for 7 days from healthy donor’s PBMCs, then coincubated with candidate epitope peptides and autologous PBLs for 14 days. Cells were harvested and stimulated by corresponding candidate peptides for another 16 hours followed by IFN-γ ICS. In some co-culture wells, the DC and peptides were co-cultured with CFSE-prelabeled PBLs for 14 days, and cells were then harvested to detect the proliferation percentage of CD8^+^ T cell. (A) Representative flow plots of IFN-γ ICS. The data in horizontal coordinates mean the frequency of IFN-γ^+^ T cells in CD3^+^/CD8^+^ T cell population. (B) Representative flow plots of CFSE staining. The data in horizontal coordinates mean the proliferation percentage of CD8^+^ T cells in CD3^+^/CD8^+^ T cell population.

### 3. Binding affinity and cross-binding of positive epitope peptides with HLA-A allotypes

HMy2.CIR is a human B lymphocyte strain with HLA class I antigen deficiency, which does not express HLA-A and B molecules and only expresses trace HLA-Cw4. To assess the affinity of the positive peptides with the corresponding HLA-A molecules, the transfected HMy2.CIR cell lines expressing indicated HLA-A molecules (HLA-A2402, A0203, A0201, A0206, A1101, A3303, A0101, or A3001) were generated firstly, sorted by flow cytometry, and identified by sequencing. The purity of these transfected CIR cell lines after sorting was 80% to 94% (Figure S4).

Then, the unlabeled positive peptides of SRAS-CoV-2 competed with fluorescent-labeled reference peptides for binding with the corresponding HLA-A molecules onto indicated transfected cell lines for 24 hours. As the FACS data showed, most positive peptides could result in a left shift of the fluorescent peak of reference peptide (Figure S5). According to the IC50, peptides were classified into three categories. Binding affinity was high when IC50 was less than 5 μM, intermediate when between 5 μM and 15μM, and low or no binding when more than 15 μM. Table 2 exhibited the binding affinity of each tested peptides with associated HLA-A molecules. These data also revealed the cross-binding of indicated peptide with different HLA-A allotypes (Table 3). Surprisingly, all epitopes derived from E protein displayed low affinity with corresponding HLA-A allotypes in this HLA-A competitive binding assay (Table 3), but these data are inconsistent with the results from DC-peptide-PBL costimulation and vaccine immunization in HLA-A2/DR1 transgenic mice.

**Table 2:**
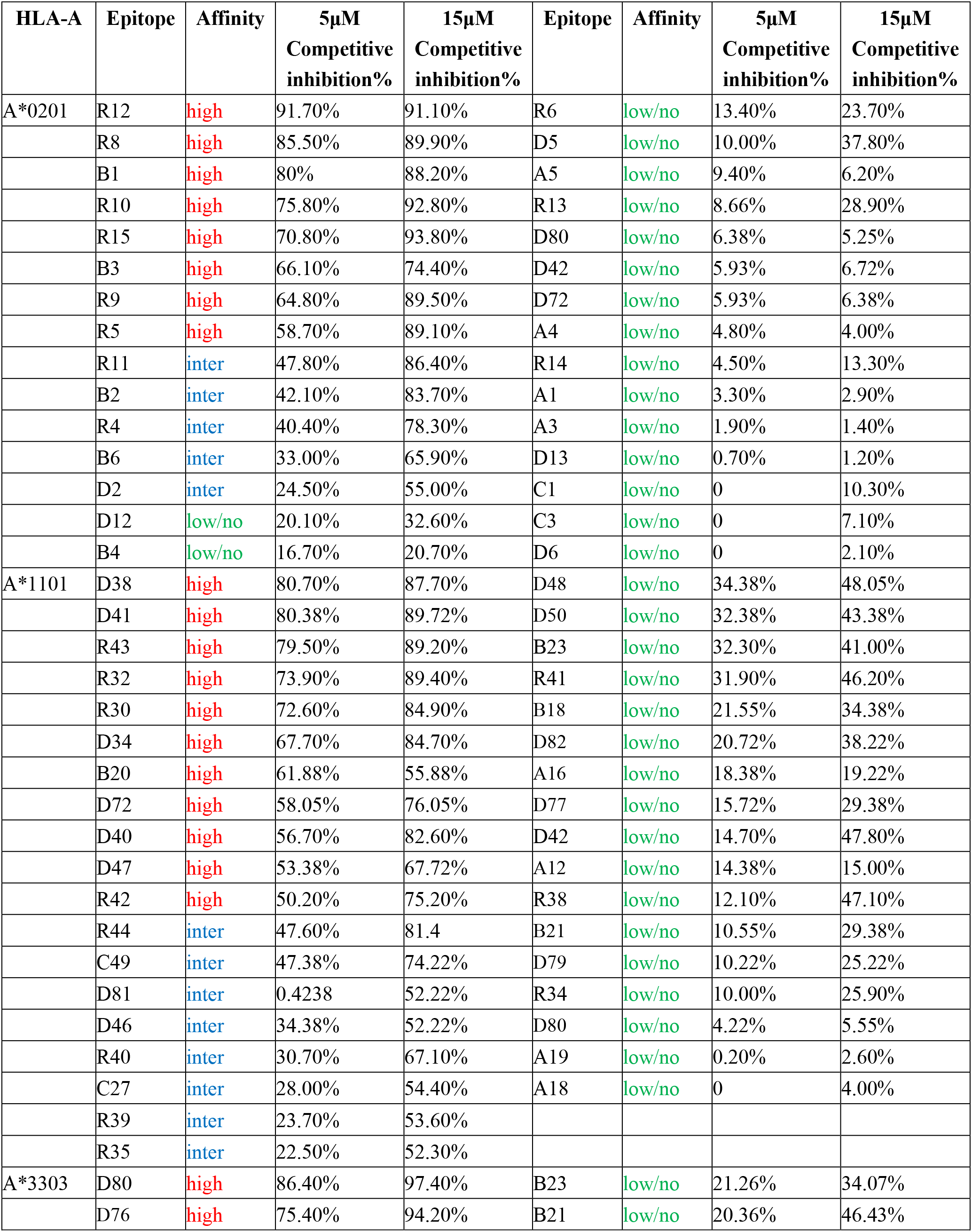
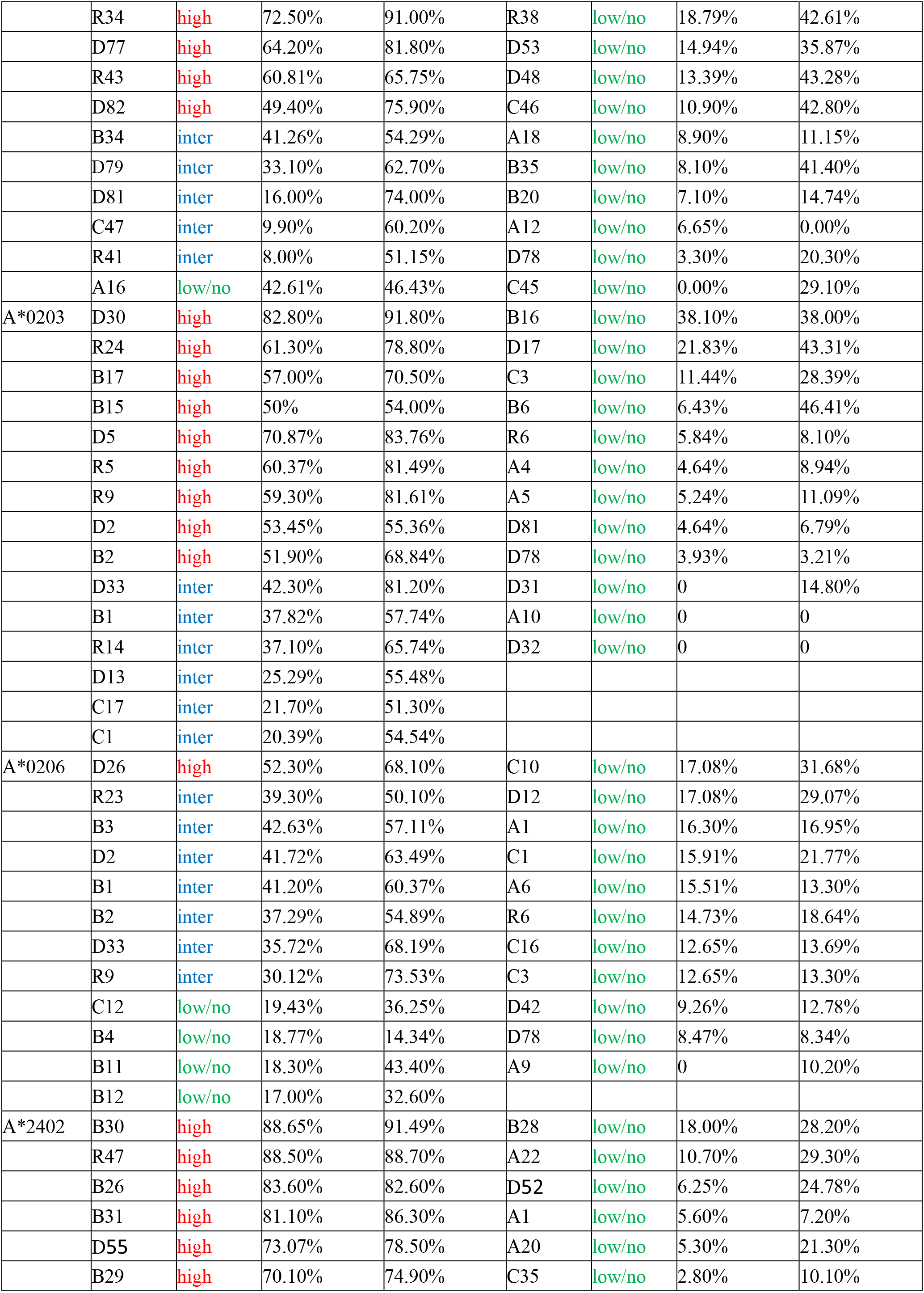
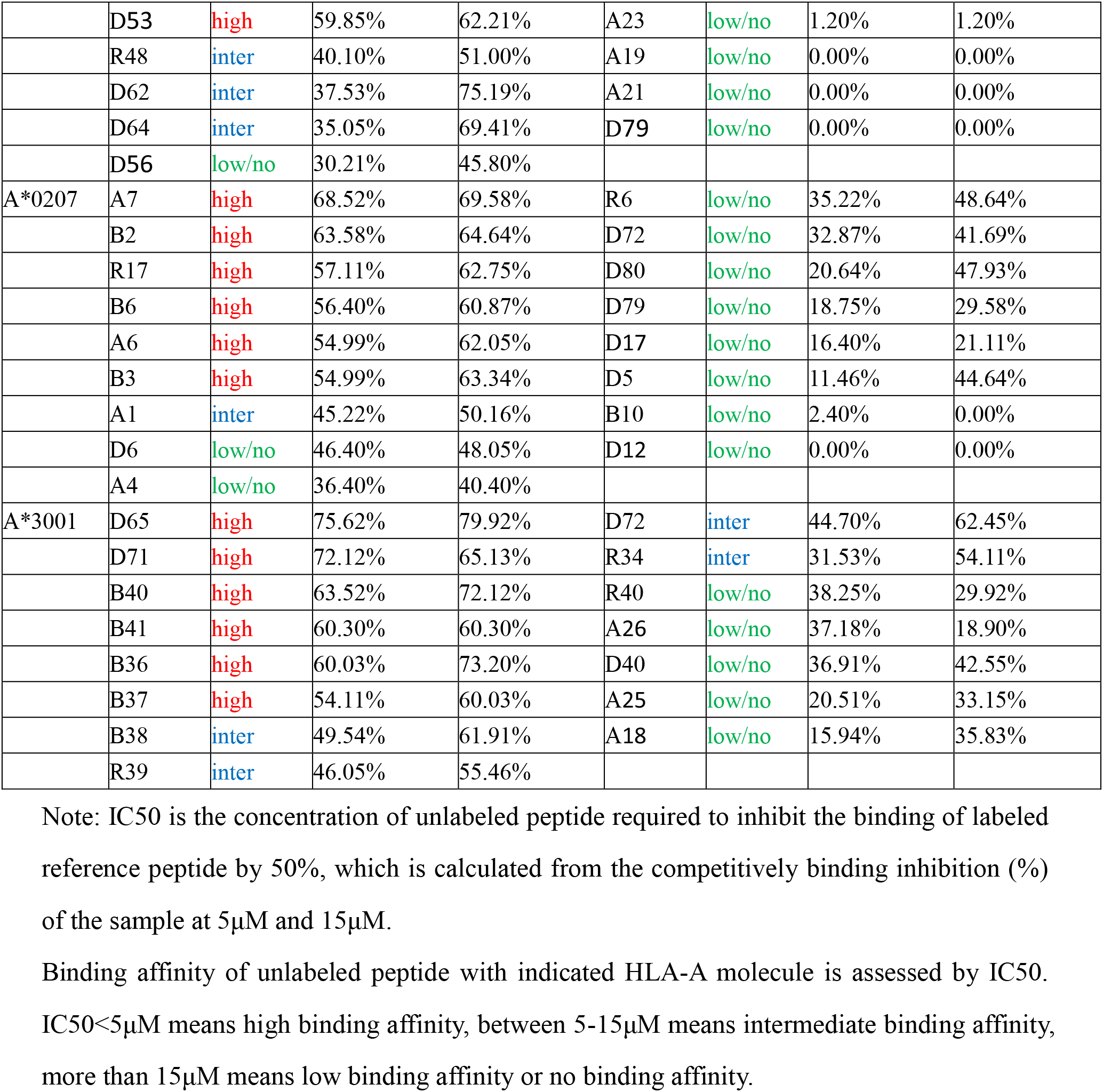
Affinity of SARS-CoV-2 CD8^+^ T cell epitopes with HLA-A allotypes as detected by HLA-A competitive binding experiments using HMy2.CIR cell lines.

**Table 3:** HLA-A restrictions of the 120 SARS-CoV-2 epitopes

| Epitope | On-silicon prediction | CIR cell lines HLA-A competitive binding assay |  |  | No test |
| --- | --- | --- | --- | --- | --- |
|  |  | High affinity | Inter affinity | low/no affinity |  |
| A1 | A0201,A0207,A0206,<br>A2402 |  | A0207 | A0206 > A2402 > A0201 |  |
| A3 | A0201 |  |  | A0201 |  |
| A4 | A0201, A0207, A0203 |  |  | A0207 > A0201 > A0203 |  |
| A5 | A0201, A0203 |  |  | A0201 > A0203 |  |
| A6 | A0207, A0206 | A0207 |  | A0206 |  |
| A7 | A0207 | A0207 |  |  |  |
| A9 | A0206 |  |  | A0206 |  |
| A10 | A0203 |  |  | A0203 |  |
| A12 | A1101, A1102, A3303 |  |  | A1101 > A3303 | A1102 |
| A16 | A1101, A3303 |  |  | A3303 > A1101 |  |
| A18 | A1102, A3303, A3001 |  |  | A3001 > A3303 > A1101 |  |
| A19 | A1102 |  |  | A1101 | A1102 |
| A20 | A2402 |  |  | A2402 |  |
| A21 | A2402 |  |  | A2402 |  |
| A22 | A2402 |  |  | A2402 |  |
| A23 | A2402 |  |  | A2402 |  |
| A25 | A3001 |  |  | A3001 |  |
| A26 | A3001 |  |  | A3001 |  |
| B1 | A0201, A0206, A0203 | A0201 | A0206 > A0203 |  |  |
| B2 | A0201, A0206, A0203,<br>A0207 | A0207 > A0203 | A0201 > A0206 |  |  |
| B3 | A0201, A0207, A0206 | A0201 > A0207 | A0206 |  |  |
| B4 | A0201, A0206 |  |  | A0206 > A0201 |  |
| B6 | A0201, A0207, A0203 | A0207 | A0201 | A0203 |  |
| B10 | A0207 |  |  | A0207 |  |
| B11 | A0206 |  |  | A0206 |  |
| B12 | A0206 |  |  | A0206 |  |
| B15 | A0203 | A0203 |  |  |  |
| B16 | A0203 |  |  | A0203 |  |
| B17 | A0203 | A0203 |  |  |  |
| B18 | A1101, A1102 |  |  | A1101 | A1102 |
| B20 | A1101, A1102, A3303 | A1101 |  | A3303 | A1102 |
| B21 | A1101, A1102 |  |  | A1101 | A1102 |
| B23 | A1101, A1102, A3303 |  |  | A1101 > A3303 | A1102 |
| B26 | A2402 | A2402 |  |  |  |
| B28 | A2402 |  |  | A2402 |  |
| B29 | A2402 | A2402 |  |  |  |
| B30 | A2402 | A2402 |  |  |  |
| B31 | A2402 | A2402 |  |  |  |
| B34 | A3303 |  | A3303 |  |  |
| B35 | A3303 |  |  | A3303 |  |
| B36 | A3001 | A3001 |  |  |  |
| B37 | A3001 | A3001 |  |  |  |
| B38 | A3001 |  | A3001 |  |  |
| B40 | A3001 | A3001 |  |  |  |
| B41 | A3001 | A3001 |  |  |  |
| C1 | A0201, A0206, A0203 |  | A0203 | A0206 > A0201 |  |
| C3 | A0201, A0206, A0203 |  |  | A0206 > A0203 > A0201 |  |
| C10 | A0206 |  |  | A0206 |  |
| C12 | A0206 |  |  | A0206 |  |
| C16 | A0206 |  |  | A0206 |  |
| C17 | A0203 |  | A0203 |  |  |
| C27 | A1101 |  | A1101 |  |  |
| C35 | A2402 |  |  | A2402 |  |
| C45 | A3303 |  |  | A3303 |  |
| C46 | A3303 |  |  | A3303 |  |
| C47 | A3303 |  | A3303 |  |  |
| C49 | A1102 |  | A1101 |  | A1102 |
| D2 | A0201, A0206, A0203 | A0203 | A0206 > A0201 |  |  |
| D5 | A0201, A0207, A0203 | A0203 |  | A0207 > A0201 |  |
| D6 | A0201, A0207 |  |  | A0207 > A0201 |  |
| D12 | A0201, A0207, A0206 |  | A0206 | A0201 > A0207 |  |
| D13 | A0201, A0203 |  | A0203 | A0201 |  |
| D17 | A0207, A0203 |  |  | A0203 > A0207 |  |
| D26 | A0206 | A0206 |  |  |  |
| D30 | A0203 | A0203 |  |  |  |
| D31 | A0203 |  |  | A0203 |  |
| D32 | A0203 |  |  | A0203 |  |
| D33 | A0203, A0206 |  | A0203 > A0206 |  |  |
| D34 | A1101, A1102 | A1101 |  |  | A1102 |
| D38 | A1101, A1102 | A1101 |  |  | A1102 |
| D40 | A1101, A1102, A3001 | A1101 |  | A3001 | A1102 |
| D41 | A1101 | A1101 |  |  |  |
| D42 | A1101, A0206, A0201 |  |  | A1101 > A0206 > A0201 |  |
| D46 | A1102 |  | A1101 |  | A1102 |
| D47 | A1102 | A1101 |  |  | A1102 |
| D48 | A1102, A3303 |  |  | A1101 > A3303 | A1102 |
| D50 | A1102 |  |  | A1101 | A1102 |
| D52 | A2402 |  |  | A2402 |  |
| D53 | A2402, A3303 | A2402 |  | A3303 |  |
| D55 | A2402 | A2402 |  |  |  |
| D56 | A2402 |  |  | A2402 |  |
| D62 | A2402 |  | A2402 |  |  |
| D64 | A2402 |  | A2402 |  |  |
| D65 | A3001 | A3001 |  |  |  |
| D71 | A3001 | A3001 |  |  |  |
| D72 | A3001,A1102,A0201,<br>A0207 | A1101 | A3001 | A0207>A0201 |  |
| D76 | A3303 | A3303 |  |  |  |
| D77 | A3303, A1102 | A3303 |  | A1101 | A1102 |
| D78 | A3303, A0203, A0206 |  |  | A0206 > A0203 > A3303 |  |
| D79 | A3303, A1101, A0207,<br>A2402 |  | A3303 | A0207>A1101 > A2402 |  |
| D80 | A3303,A1102,A0201,<br>A0207 | A3303 |  | A0207>A0201 > A1101 | A1102 |
| D81 | A3303, A1102, A0203 |  | A1101 > A3303 | A0203 | A1102 |
| D82 | A3303, A1102 | A3303 |  | A1101 | A1102 |
| R4 | A0201 |  | A0201 |  |  |
| R5 | A0201, A0203 | A0203 > A0201 |  |  |  |
| R6 | A0201,A0207,A0206,<br>A0203 |  |  | A0207>A0206>A0201><br>A0203 |  |
| R8 | A0201 | A0201 |  |  |  |
| R9 | A0201, A0206, A0203 | A0201 > A0203 | A0206 |  |  |
| R10 | A0201 | A0201 |  |  |  |
| R11 | A0201 |  | A0201 |  |  |
| R12 | A0201 | A0201 |  |  |  |
| R13 | A0201 |  |  | A0201 |  |
| R14 | A0201, A0203 |  | A0203 | A0201 |  |
| R15 | A0201 | A0201 |  |  |  |
| R17 | A0207 | A0207 |  |  |  |
| R23 | A0206 |  | A0206 |  |  |
| R24 | A0203 | A0203 |  |  |  |
| R30 | A1101 | A1101 |  |  |  |
| R32 | A1101, A1102 | A1101 |  |  | A1102 |
| R34 | A1101, A3001, A3303 | A3303 | A3001 | A1101 |  |
| R35 | A1101, A1102 |  | A1101 |  | A1102 |
| R38 | A1101, A3303 |  |  | A3303 > A1101 |  |
| R39 | A1101, A1102, A3001 |  | A1101 |  | A1102 |
| R40 | A1101, A3001 |  | A1101 | A3001 |  |
| R41 | A1101, A3303 |  | A3303 | A1101 |  |
| R42 | A1101 | A1101 |  |  |  |
| R43 | A1101, A3303 | A1101 > A3303 |  |  |  |
| R44 | A1101, A1102 |  | A1101 |  | A1102 |
| R47 | A2402 | A2402 |  |  |  |
| R48 | A2402 |  | A2402 |  |  |

Note: The HLA-A molecules highlighted in red displayed no affinity with indicated epitopes.
“>” means the affinity from high to low.

### 4. The peptide cocktail vaccines induced robust specific CD8^+^ T cell responses in HLA-A2/DR1 transgenic mice

To determine whether these peptides which was validated by DC-peptide-PBL co-culture experiments can stimulate T cell responses *in vivo*, 31 positive epitope peptides restricted by HLA-A2 molecule (HLA-A0201, A0203, A0206, A0207) were grouped into 4 peptide pools (Table S3) and were used to generate peptide cocktail vaccines in three formulations: the peptides-encapsulated and -surface coupled PLGA-NPs/peptides (Vaccine A), R848/peptides (Vaccine B), and poly I:C/peptides (Vaccine C) (Table S4). Additionally, T2 cell binding assay was used to define the affinity of the 31 epitopes with HLA-A0201 molecule. T2 cells were incubated with indicated peptide or no peptide and β2-microglobulin for 16 hours. Then, peptide-induced up-regulation of HLA-A0201 expression on T2 cells was measured by PE-labeled anti-HLA-A2.1 antibody staining and flow cytometry (Figure S6). According to the fluorescence index (FI), 18 epitopes showed high affinity (FI > 1.0), 7 epitopes displayed intermediate affinity (1.0 ≥ FI >0.5), and 6 epitopes exhibited low or no binding (FI ≤ 0.5) with HLA-A0201 molecules (Table S5). Finally, HLA-A0201^+/+^/DR1^+/+^ transgenic and H-2-β2m^-/-^/I-Aβ^-/-^ C57BL/6 mice were immunized with the three vaccines respectively. After three rounds *in vivo* stimulation, splenocytes of primed mice were tested for peptide-specific T cell responses by IFN-γ-ELSPOT, IFN-γ-ICS and IFN-γ-ELISA.

The 31 positive peptides were grouped into eight pools (Table S3) according to their derived proteins and the features of acid and alkalinity. Then splenocytes from each mouse were coincubated with each peptides pool or PBS for 20 hours in 96-well PVDF membrane plate and followed by IFN-γ-ELSPOT assay. The total spot forming unit (SFU) in 2×10^5^ splenocytes from each mouse in the three vaccine groups was 400-500 times more than that from control group (Figure 3A). Interestingly, splenocytes in all vaccine groups showed almost the strongest T cell responses to E protein and weakest T cell responses to N protein relative to other antigens (Figure 3B). Figure 4 presented the spots of ELISPOT assay from all mice. Two irrelevant CD8^+^ T cell epitope peptides (AFP_158-166_ and AFP_424-432_) were used as antigen-irrelevant control groups and obtained the negative results similar to that in no peptide group.

**Figure 3:**
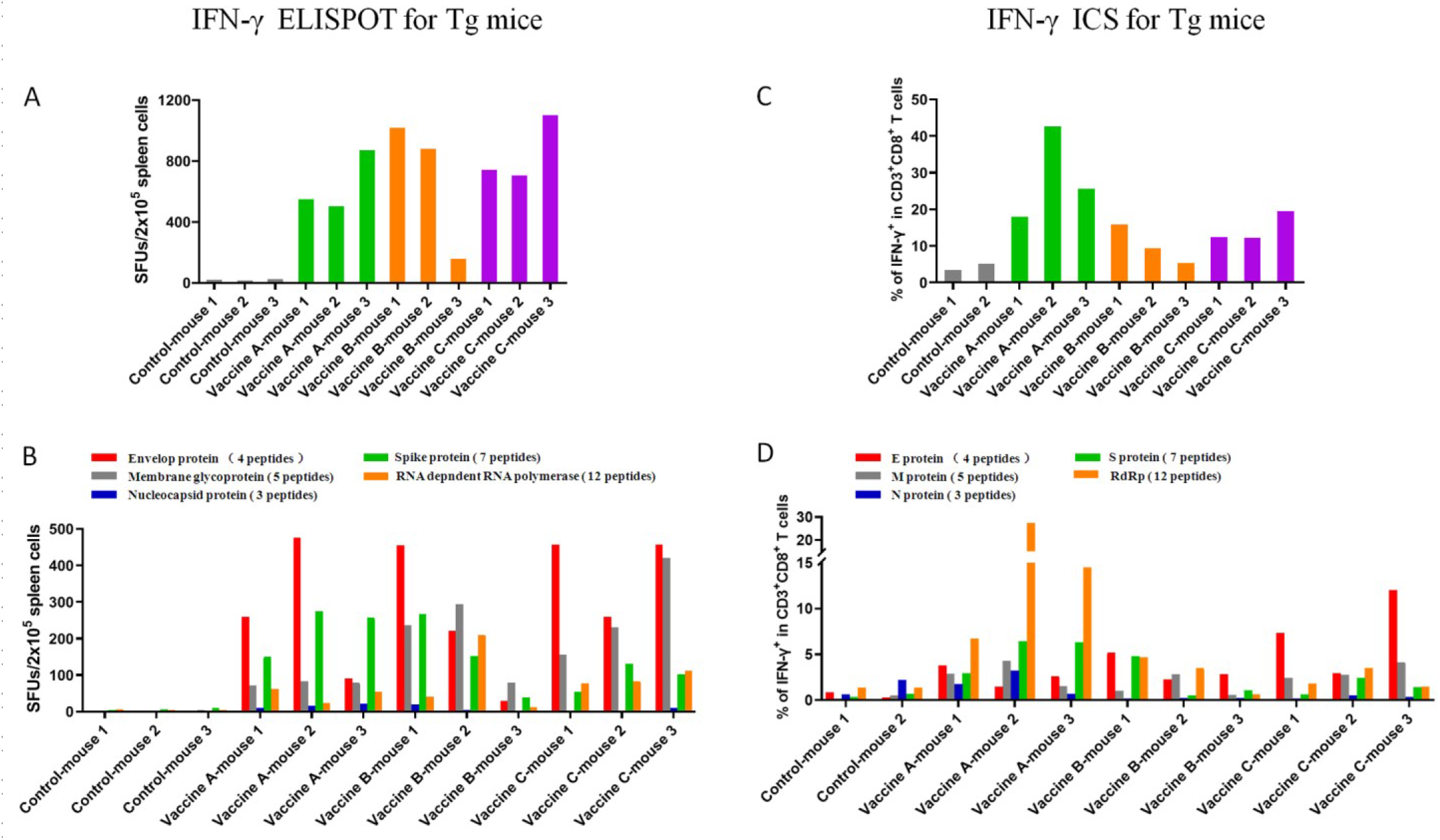
T cell epitope peptide cocktail vaccines elicited robust CD8^+^ T cell responses in transgenic mice. 31 positive epitope peptides restricted by HLA-A2 molecule were used to generate peptide cocktail vaccines in three formulations, and followed by three-round immunizations of HLA-A2/DR1 transgenic C57BL/6 mice. Then, splenocytes were collected 7 days after the last booster and *ex vivo* stimulated with distinct peptide pools according to single protein overnight, followed by IFN-γ ELISPOT and IFN-γ ICS. (A) Total IFN-γ SFUs responding to all peptide pools in each mouse. (B) Deconvolution of the total SFUs in each mouse from A into the single SARS-CoV-2 proteins. (C) Total frequency of IFN-γ^+^ T cells reacting to all peptide pools in CD3^+^CD8^+^ T cell population in each mouse. (D) Deconvolution of the total frequency in each mouse from C into the single SARS-CoV-2 proteins. Control group: N.S and PLGA-NPs; Vaccine A group: PLGA-NPs/peptides vaccines; Vaccine B group: R848/peptides vaccines; Vaccine C group: poly I: C/peptides vaccines.

**Figure 4:**
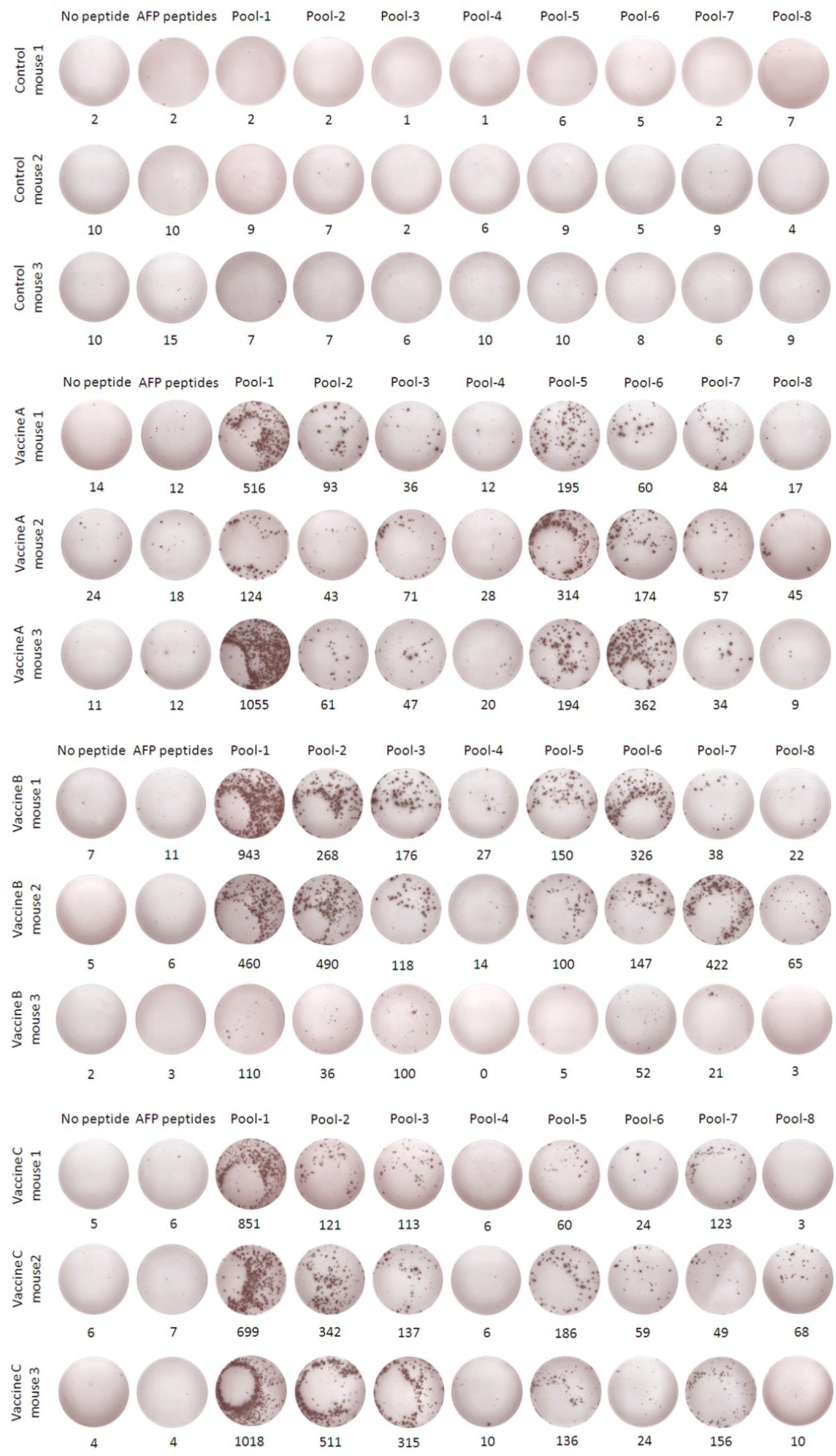
IFN-γ ELISPOT responses against the individual peptide pools. Splenocytes from each primed mouse were harvested 7 days after the last booster and *ex vivo* stimulated with 8 different peptide pools covering the 31 epitope peptides or with AFP peptides (AFP_158-166_, AFP_424-432_) as irrelevant control, or without peptide as negative control, and followed by IFN-γ ELISPOT.

To further confirm the results from ELISPOT assay and the specific CD8^+^ T cell responses, IFN-γ intracellular cytokine staining was performed. The 31 positive peptides were grouped into five pools (Table S3) according to their derived proteins. Then splenocytes from each mouse were coincubated with each peptides pool or PBS for 16 hours in 48-well plate and followed by another 6 hours coincubation with BFA/Monensin mixture. The resulting ICS showed that the frequencies of IFN-γ^+^ in CD3^+^CD8^+^ T cell populations from the three vaccine groups were about 20-30 times higher than that in control mice (Figure 3C). Splenocytes in vaccine A group showed the strongest CD8^+^ T cell responses to RdRP while splenocytes in vaccine C group showed the strongest CD8^+^ T cell responses to E protein (Figure 3D). Figure 5 presented the flow plots for ICS from all mice. Also, two irrelevant CD8^+^ T cell epitope peptides (AFP_158-166_ and AFP_424-432_) were used and obtained the results similar to the no peptide group.

**Figure 5:**
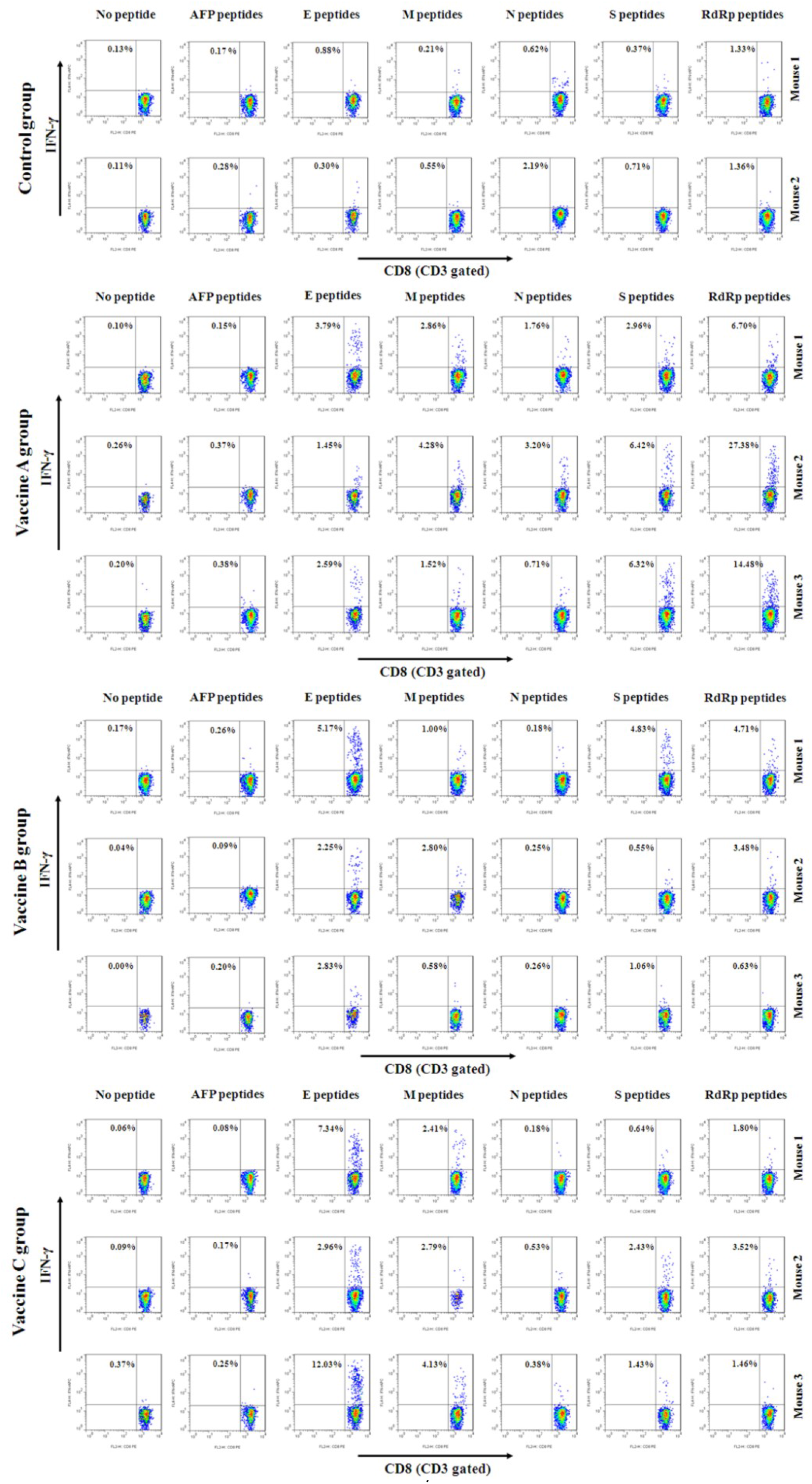
Flow plots of IFN-γ ICS responding to the individual peptide pools. Splenocytes from each primed mouse were harvested 7 days after the last booster and *ex vivo* stimulated with 5 different peptide pools according to single protein or with AFP peptides (AFP_158-166_, AFP_424-432_) as irrelevant control, or without peptide as negative control, and followed by IFN-γ ICS. The data in left upper quadrant mean the frequencies of IFN-γ^+^ T cells in CD3^+^/CD8^+^ cell populations.

Furthermore, ELISA was carried out to quantify IFN-γ in the supernatant after the splenocytes were incubated with each of the five peptide pools or PBS in 48-well plate for 72 hours. Accumulations of IFN-γ in the supernatant of the three vaccine groups were about 15-30 times higher than that in control group (Figure 6A), which is consistent with the results of ELISPOT and ICS. Splenocytes in all three vaccine groups showed relatively stronger responses to RdRP protein (Figure 6B).

**Figure 6:**
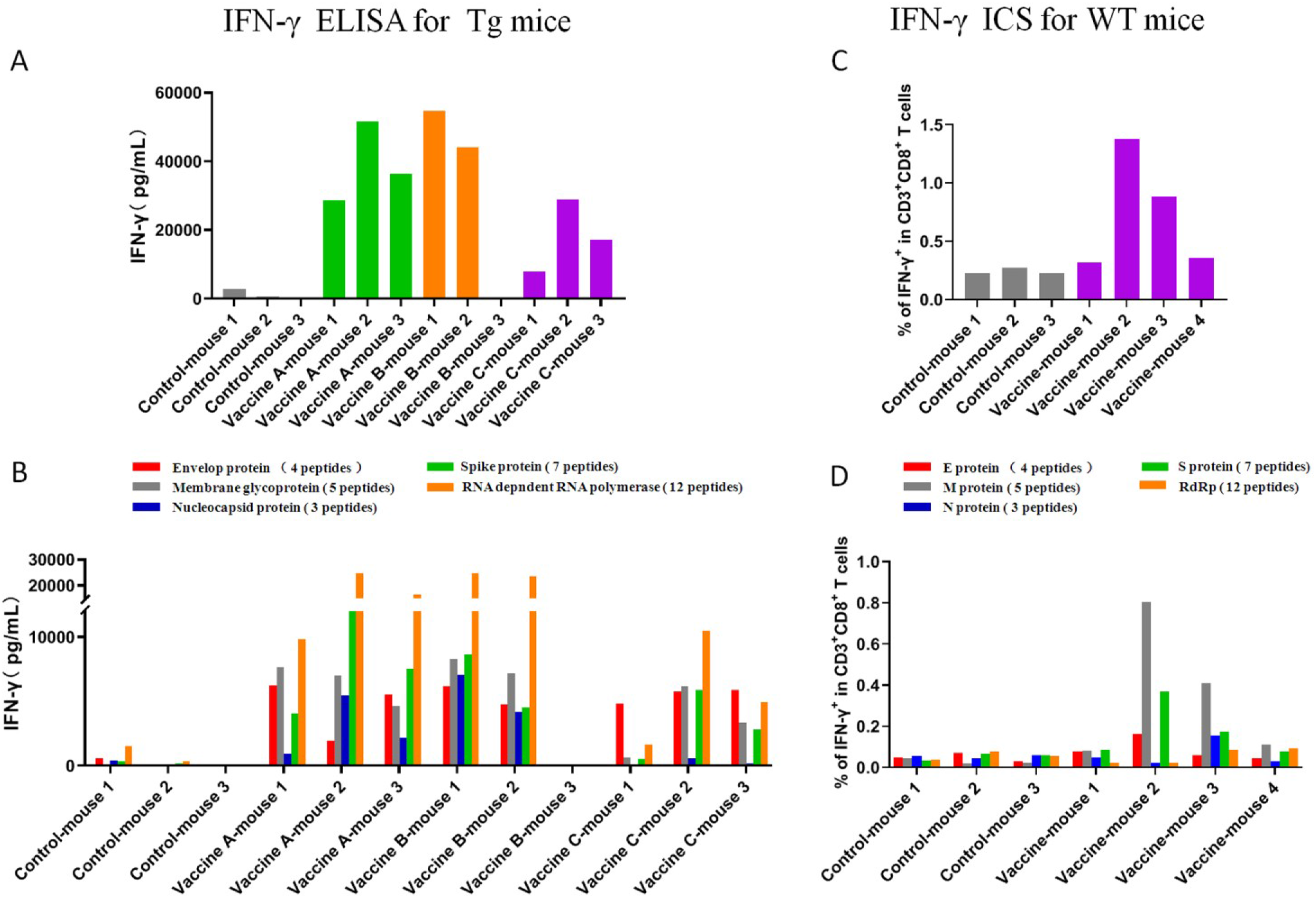
T cell epitope peptide cocktail vaccines elicited robust CD8^+^ T cell responses as detected by IFN-γ ELISA in transgenic mice and by IFN-γ ICS in WT mice. Splenocytes from each primed HLA-A2/DR1 tansgenic C57BL/6 mice were harvested 7 days after the last booster and *ex vivo* stimulated with 5 different peptide pools according to single protein or without peptide for 3 days. Then supernatants were collected and followed by IFN-γ ELISA. (A) Total IFN-γ level responding to all peptide pools in each mouse. (B) Deconvolution of the total IFN-γ level in each mouse from A into the single SARS-CoV-2 proteins. Similarly, wild-types C57BL/6 mice were divided into two groups and were immunized three times with the Poly I:C/peptides vaccines prepared using the 31 validated epitope peptides or normal saline, respectively. Splenocytes were then collected 7 days after the last booster and *ex vivo* stimulated with 5 different peptide pools overnight, followed by IFN-γ ICS. (C) Total frequency of IFN-γ^+^ T cells reacting to all peptide pools in CD3^+^CD8^+^ T cell population in each mouse. (D) Deconvolution of the total frequency in each mouse from C into the single SARS-CoV-2 proteins.

Taken together, these results indicated that all three forms of epitope peptide cocktail vaccines can stimulate robust specific CD8^+^ T cell responses in HLA-A2/DR1 transgenic mice, implying the potential of these validated SARS-CoV-2 T cell epitopes to be applied in vaccine development. Of note, the mouse-3 in Vaccine B group (R848/peptide vaccine) showed only weak or no T cell responses as detected by ELSPOT, ICS and ELISA. This failure may be due to the weak reactivity of entire T cell repertoire since the SFU of 2×10^5^ splenocytes was much less than other primed mice after stimulated by mitogen PHA as detected by ELISPOT (70 vs. 832.2 ± 328.9). In order to further confirm the *in vivo* results, Vaccine C (poly I:C/peptides) immunization experiment was repeated in the HLA-A0201^+/+^/DR1^+/+^ transgenic and H-2-β2m^-/-^/I-Aβ^-/-^ C57BL/6 mice and induced similar trend of robust CD8^+^ T cell responses compared to control group.

### 5. The peptide cocktail vaccine induced CD8^+^ T cell responses in wild-type C57BL/6 **mice**

To investigate whether the HLA-A2 molecules-restricted 9 or 10-mer peptides can also be cross-presented by mouse H-2K/D^b^ molecules, the wild-type C57BL/6 mice were immunized with vaccine C (peptide pool-v1, pool-v2, pool-v3 and pool-v4 mixed with poly I:C). After three rounds of *in vivo* stimulation according to the timeline in HLA-A2/DR1 transgenic mice, splenocytes from primed C57BL/6 mice were detected by ICS. The frequencies of IFN-γ^+^ T cells in CD3^+^/CD8^+^ populations were 4-7 times higher in the 2 of 4 vaccination mice than that in the control group (Figure 6C). The robust CD8^+^ T cell responses were mainly against the epitopes from M, N and S protein (Figure 6D). Figure 7A presented the flow plots of all mice. Also, two irrelevant CD8^+^ T cell epitope peptides (AFP_158-166_ and AFP_424-432_) were used and obtained the results similar to the no peptide group.

**Figure 7:**
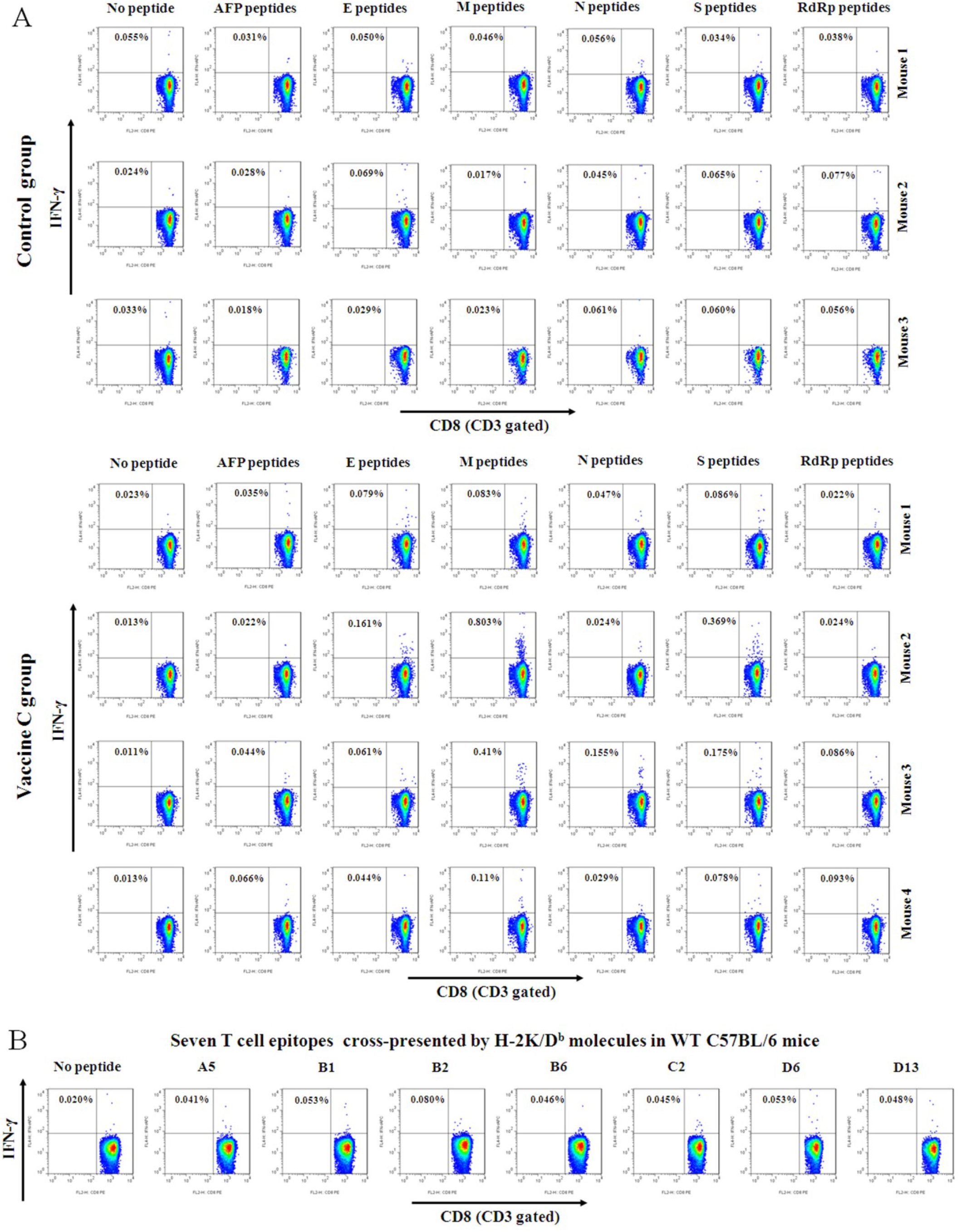
Flow plots of IFN-γ ICS responding to the individual peptide pools after WT mice immunizations. (A) Splenocytes from each primed WT mouse were harvested 7 days after the last booster and *ex vivo* stimulated with 5 different peptide pools according to single protein or with AFP peptides (AFP_158-166_, AFP_424-432_) as irrelevant control, or without peptide as negative control, and followed by IFN-γ ICS. The data in left upper quadrant mean the frequencies of IFN-γ^+^ T cells in CD3^+^/CD8^+^ cell populations. (B) 7 of 31 epitopes restricted by HLA-A2 were identified to be cross-presented by H-2K/D^b^ molecules. Splenocytes from each primed WT mouse were harvested as described and *ex vivo* stimulated with single peptide, or without peptide as negative control, and followed by IFN-γ ICS. The epitopes were identified as immunogenic peptides when the frequency of IFN-γ^+^ T cells in CD3^+^/CD8^+^ cell population increased by more than 100% compared with the negative control. Representative flow plots of the 7 positive epitopes were shown.

To further identify the epitopes cross-presented by mouse H-2K/D^b^ molecules, the splenocytes from primed Vaccine mouse-2 and Vaccine mouse-3 were coincubated with each of the 31 positive epitope peptides and followed by ICS. Flow cytometric data showed that 7 of 31 epitope peptides (A5, B1, B2, B6, C2, D6 and D13) can activate the primed splenocytes with a frequency of IFN-γ^+^ T cells in CD3^+^/CD8^+^ population two times higher than that of no *ex vivo* stimulation group (Figure 7B).

### 6. T cell epitope-based peptide cocktail vaccines do not lead to visible organ toxicity

To uncover whether the peptide-based vaccine immunizations cause organ toxicity, the heart, liver, lung and kidney from each mouse were checked at day 28 after the mice were inoculated three times with Vaccine A, Vaccine B or Vaccine C. The organs were immersed and stained with Hematoxylin-Eosin. As the scanning copy showed, no visible organ toxicity was found in all organs in each group (Figure S7).

## Discussion

SARS-CoV-2 has seriously hazarded public health and the economic development all over the word. Many people are suffering from the physical symptoms or the adverse impacts caused by this major epidemic disease. Despite of great efforts made by scientists around the world, the epidemic situation is still out of control. COVID-19 vaccine development is of major importance, but mainly biased towards neutralizing antibody protection with generally less effective at eliciting CD8^+^ T cell responses, which faces possible risk in clearing virus and in preventing from infection. Informed by protective immunity observed in natural infection, people have known that vaccine approaches that elicit antiviral SARS-CoV-2 specific CD4^+^ and CD8^+^ T cells in coordination with neutralizing antibodies will generate more robust and durable protective immunity^20, 33^. As known, memory T cell responses can persist for 6-17 years after SARS-CoV infection^10, 34^ and, in mice, protect against lethal virus challenge^35^. In contrast, memory B cells live short in host^34, 35^.

However, only several reports are thus far available about the vaccine candidate T cell epitopes that have been validated by functional experiments. 408 CD8^+^ T cell epitopes on-silicon predicted were applied in multiplexed peptide-MHC tetramer staining to detect the PBMCs from 30 convalescent COVID-19 patients, then 132 positive reactions were obtained with the validation of 42 CD8^+^ T cell epitopes presented by 6 HLA allotypes (HLA-A0101, A0201, A0301, A1101, A2402 and B0702)^32^. Prackar et al tested the binding stability of 777 CD4^+^ and CD8^+^ T cell epitopes that were predicted to be good binders across 11 MHC allotypes using an *in vitro* peptide MHC stability assay, and found that 174 peptides can stably bind to the HLA allotypes (HLA-A0101, A0201, A0301, A1101, A2402, B4001, C0102, C0401, C0701, C0702, DRB10401), of which 126 have not been reported previously^31^. After 14-day cocultures of predicted epitope pools with PBMCs from healthy individuals, 142 CD4^+^ T cell epitopes were defined by immunofluorescence spot assay (FluoroSpot)^30^. Furthermore, using genome-wide screening technology (T-Scan), 29 CD8^+^ T cell epitopes were identified, and presented by 6 different HLA allotypes (A0201, A0101, A0301, A1101, A2402, B0702)^29^. Thus far, the validated SARS-CoV-2 T cell epitopes, especially CD8^+^ T cell epitopes, are still limited and few epitopes have been used *in vivo* as vaccines. More importantly, these validated epitopes are only presented by a few HLA allotypes which can not cover a broad population in indicated geographical regions. This may hamper the development of T cell epitope vaccines and the precise evaluation on herd cellular immune protection.

Unlike the previous researches about T cell epitopes of SARS-CoV-2, this study has four points of worth noting. First, this study focus on a series of high-frequency HLA-A allotypes which gather a total HLA-A allele frequency of around 87% in Chinese population while 79%, 78%, 63%, 59.5%, 49.5% and 46.5% in Southeast Asia, Northeast Asia, Indonesia, South America, Europe and North America populations, respectively (http://www.allelefrequencies.net). Since T cell epitopes spread across the proteome of SARS-CoV-2 in a relatively equal distribution^9^, here four structural proteins and the RdRp consisted of nsp7, nsp8 and nsp12 were screened for the identification of CD8^+^ T cell epitopes cross-restricted by the high-frequency HLA-A allotypes. This study has provided a library of CD8^+^ T cell epitopes that not only covers broad antigenic targets recognized by SARS-CoV-2-specific CD8^+^ T cell clones, but also fits to the Asian herd genetic characteristics of HLA molecules, thus will facilitate the development of SARS-CoV-2 vaccines inducing antiviral CD8^+^ T cell responses for Asian populations.

Second, this study established a modified cell co-culture system of DC-peptide-PBL using healthy donor’s PBMCs to validate the immunogenicity of CD8^+^ T cell epitope on-silicon predicted. The most reliable and valuable method to validate the immunogenicity of candidate T cell epitopes is the detection of epitope-specific memory T cell clones in the PBMCs or other cell samples from COVID-19 patients or convalescent humans. However, it is less practicable in current China due to the difficulty obtaining clinical blood samples. So, an alternative approach using PBMCs from unexposed humans was used here. The DC-T or peptide-PBMC co-culture procedures using healthy donor’s PBMCs have been generally used to validate the CD4^+^ T cell epitopes by the exogenous antigen presenting mechanism^30^. In this study, DC-T and peptide-PBMC experiments were integrated as DC-peptide-PBL co-stimulation system, and initially applied for the validation of CD8^+^ T cell epitopes. DCs were induced from the healthy donors’ PBMCs, and coincubated for 14 days with candidate epitope peptides and autologous PBLs. In this co-culture system, the 9-mer or 10-mer peptides are maintained in culture media at a high concentration of 20 μg/mL during 14 days, and may be engulfed by DCs and cross-presented to CD8^+^ T cell by HLA-A molecules, or directly bind to the HLA-A molecules onto DCs and B cells followed by activation of naive CD8^+^ T cells or SARS-CoV-2 cross-reactive memory CD8^+^ T cells. This presumption was verified at least in part by the positive results of DC-peptide-PBL experiments in this study. Many candidate epitope peptides can increase the frequency of IFN-γ^+^ CD8^+^ T cells by 3-5 times in this co-culture system. Compared with the ICS, the detection of CD8^+^ T cell proliferation by using CSFE-prelabeled PBLs is less sensitive. Of note is that the healthy donor’s DC-T *in vitro* co-cultures usually can increase peptide-specific CD4^+^ T cells by around 10 times since the exogenous antigen procession and presentation mechanism^30^. More interestingly, 44 (36.66%) of the 120 validated CD8^+^ T cell epitope peptides can also simultaneously activate CD4^+^ T cells with the 2-6 times increase of IFN-γ^+^/CD4^+^ T cell frequency or CD4^+^ T cell proliferation (data not shown). The underlying mechanism remains to be further elucidated.

In order to further confirm the sensitivity and reliability of this co-culture system, some reference CD8^+^ T cell epitope peptides, which were derived from HCC-associated tumor antigens or from hepatitis B virus antigens and have been validated as real-world epitopes previously in-house by using HCC patients’ PBMCs or chronic hepatitis patient’s PBMCs, were tested in this system and achieved positive results. In addition, HLA-A molecule competitive binding experiments also were used to confirm the binding affinity of the 120 positive peptides with corresponding HLA-A molecules onto HMy2.CIR cell lines. The results are mostly consistent in that of DC-peptide-PBL experiments except the epitopes derived from E protein, and further identified the HLA-A restrictions and cross-restrictions of each epitope peptides. Furthermore, the robust CD8^+^ T cell responses elicited by the HLA-A2-binding peptide cocktails in HLA-A2/DR1 transgenic mice also indicated the *in vivo* immunogenicity of representative epitope peptides validated by this DC-peptide-PBL procedure.

For this study, the third point different from previous researches is that the HLA-A cross-restrictions of 120 positive CD8^+^ T cell epitopes were further identified by using HLA-A molecule competitive binding experiments with HMy2.CIR cell lines expressing indicated HLA-A molecules. As known, one T cell epitope can be presented by several HLA allotypes with distinct binding affinity. The HLA molecule restrictions and cross-restrictions of most SARS-CoV-2 T cell epitopes previously reported have not been elucidated, but only have been affirmed by on-silicon prediction and mere guesswork according to the donor’s HLA allele genotypes^29, 30, 32^.

More importantly, this study initially confirmed the *in vivo* feasibility of 9 or 10-mer peptide cocktail vaccines of SARS-CoV2. As known, HLA class II molecule restricted peptides (15 or 16-mer long) can induce CD4^+^ T cell responses *in vivo*. For SARS-CoV2, HLA-DR restricted peptides cocktail vaccine from Tubingen University of Germany has been enrolled in phase I clinical trial (NCT04546841). However, few 9-mer or 10-mer CD8^+^ T cell epitope peptides have been directly used *in vivo* as peptide vaccines. For tumor immunotherapy, tumor neoantigen peptide vaccines and mRNA vaccines have recently achieved robust T cell responses and encouraging clinical outcome^36, 37^. In clinical trials of melanomas^38, 39^ and non-small cell lung cancers^40^, the survival of patients was significantly prolonged and the recurrence rate was reduced. Patrick A. Ott^38^ identified up to 20 neoantigen peptides restricted by HLA-A or B allotypes from each melanoma patient using whole-exome sequencing of matched tumor and normal cell DNA and RNA sequencing of the tumor. Herein, the long peptides (with the length of 15–30 amino acids containing individual CD8^+^ T cell epitope) rather than 9-mer or 10-mer peptides were synthesized and mixed with TLR3 agonist poly-ICLC to generate personal tumor neoantigen peptide vaccines. Of 6 vaccinated melanoma patients, 4 had no recurrence at 25 months post-vaccination, while 2 with progressive disease were subsequently treated with anti-PD-1 therapy and experienced complete tumor regression. The long peptide vaccines elicited expansion of the repertoire of neoantigen-specific T cells in each patient, but most were CD4^+^ T cells rather than CD8^+^ T cells. In this study, 31 short peptides of SARS-CoV-2 CD8^+^ T cell epitope (with the length of 9-10 amino acids) were synthesized and mixed with R848, poly (I:C) or PLGA-NPs followed by immunizations in HLA-A2/DR1 transgenic mice and WT mice. The resulting data, for the first time to our knowledge, provide the experimental evidences that human MHC class I molecule-restricted short peptide cocktail vaccines can induce robust SARS-CoV2 specific CD8^+^ T cell responses *in vivo*.

Using patients’ PBMCs could only test whether the candidate peptide can be recognized by memory T cells *ex vivo*, while using healthy donor’ PBMCs only test whether the candidate peptides can elicit naïve T cells *in vitro*. Whether the candidate peptide is also able to activate naïve T cells *in vivo* is a better criterion to judge it as an ideal vaccine candidate peptide. As compared with the Ad5-ACE2-transduced and SARS-CoV-2-infected BALB/c and C57BL/6 mice^21^, HLA transgenic mice integrated with human HLA class I and II alleles are more suitable for preclinical study of the vaccine. HLA-A*02:01 is one of the most common HLA class I alleles in the world, thus HLA-A*02:01 transgenic mice (HHD mice) have been generally used in identifying HLA-A0201-restricted epitopes and in evaluating peptide vaccines ^41–43^. Herein, the HLA-A0201^+/+^/DR1^+/+^/H-2-β_2_m^-/-^/IAβ^-/-^ C57BL/6 mice were used in which the 9-mer or 10-mer peptides cocktail vaccines only can be presented by HLA-A2 or DR1 molecules, not by any H-2^b^ molecules, which indicated that this mouse model is more suitable to mimic the in-human antigen procession and presentation mechanism, without the interference caused by mouse H-2 molecules presentation. However, similar to the DC-peptide-PBL co-culture system, the detailed *in vivo* mechanism by which the exogenous 9 or 10-mer peptides elicit naive CD8^+^ T cells activation remains unclear.

The 120 CD8^+^ T cell epitopes spread across the E, M, N, S, and RdRp proteins of SARS-CoV-2 with a bias distribution. Notably, the shortest E protein presented the highest density of CD8^+^ T cell epitope (2.400/10aa), two times relative to M protein (1.216/10aa), while N, S and RdRp proteins exhibited a similar low density (0.286, 0.283, 0.290 per 10aa) in this study. These data are discriminated with other findings in which T cell epitopes relatively equally distributed across the proteome of SARS-CoV-2^9^.

Of the unexposed cohorts from the United States, Netherlands, Germany, Singapore, and United Kingdom, 20-50% could detect specific memory CD4^+^ T cells that showed cross-reactivity to SARS-CoV-2 antigens, but the frequency was about 10 times lower (0.1% vs.1%) than that of those infected with SARS-CoV-2, presumably due to the common T cell epitopes between SARS-CoV-2 and common-cold HCoVs^8, 9, 30, 44^. In addition, long-lasting memory T cells reactive to N protein of SARS-CoV can be detected in 2020 in the convalescent individuals with a history of SARS-CoV infection in 2003, and showed robust cross-responses against N protein of SARS-CoV-2^10^. In this study, depending on the sequence homologous alignment of SARS-CoV-2 with SARS-CoV, common-cold HCoVs, and current mutant variants of SARS-CoV-2, most epitopes (110) were common CD8^+^ T cell epitopes of SARS-CoV. Additionally, 15, 6, 14 and 12 epitopes were also highly homologous (0-2 amino acids deviation) with OC43, NL63, HKU1, and 229E, respectively, which are mainly derived from in RdRp protein (39/47). These theoretical data support the previous findings in unexposed humans, but are contradictory with the previous research in which CD8^+^ T cells generally do not cross-react with epitopes in the four common cold HCoVs^29^. Further detection using these homologous epitope peptides in unexposed cohort is needed. More importantly, of which 120 validated CD8^+^ T cell epitopes of SARS-CoV-2, only 4 epitopes from S protein displayed one amino acid distinct from the five wide-spread virus variants while the one amino acid deviation at position 1, 3, 5 or 6 may or may not change epitope immunogenicity, suggesting the advantage of T cell epitope cocktail vaccine over the B cell epitope vaccine producing neutralizing antibody on the antiviral protections against mutant SARS-CoV-2.

In summary, 120 kinds of CD8^+^ T cell epitopes derived from E, M, N, S, and RdRp proteins of SARS-CoV-2 and restricted by a series of high-frequency HLA-A allotypes were identified and validated. Among them, 110 and 47 epitopes are highly homologous with SARS-CoV and common-cold HCoVs respectively, 4 epitopes are distinct from current variants of SARS-CoV-2 with one amino acid substitution. HLA-A2-restricted 31 epitopes were generated as short peptide cocktail vaccines and triggered robust CD8^+^ T cell responses in HLA-A2/DR1 transgenic C57BL/6 mice and wild-type C57BL/6 mice. 7 epitopes were found to be cross-presented by HLA-A2 and H-2K/D^b^ molecules. Whether these CD8^+^ T cell responses elicited by the peptides could facilitate virus clearance will be studies further.

## Materials and methods

### 1. Preparation of PBMCs and HLA-A gene typing

The white blood cell filter trays after red blood cells preparation of healthy blood donors were gotten from Blood Component Preparation Section of Jiangsu Province Blood Center. Then white blood cells were collected from the white blood cell filter tray and PBMCs were isolated by density-gradient centrifugation using Ficoll-Paque. The fresh PBMCs were either used directly or cryopreserved in -80 °C until further test. HLA-A alleles were identified by PCR-sequencing-based tying. The human samples collection and use has been approved by Clinical Ethics Committee of Affiliated Zhongda hospital of Southeast University.

### 2. Mice

Female HLA-A*02:01/DR1 transgenic and H-2-β_2_m^-/-^/IAβ^-/-^ C57BL/6 mice at 10 weeks were generous gifts from Academy of Military Medical Sciences. Female C57BL/6 mice at 10 weeks of age were purchased from the Comparative Medicine Center of Yangzhou University (Yangzhou, China). Mice were maintained at the specific pathogen-free Animal Centre of Southeast University (Nanjing, China). Animal welfare and experimental procedures were performed in accordance with the Guide for the Care and Use of Laboratory Animals (Ministry of Science and Technology of China, 2006) and were approved by the Animal Ethics Committee of Southeast University.

### 3. On-silicon prediction of T cell epitopes and peptide synthesis

T cell epitopes spanning E, M, N, S, and RdRP proteins of SARS-CoV-2 (Wuhan strain) and presented by different HLA-A molecules were on-silicon predicted using five epitope predication tools and seven types of algorithms (IEDB-ANN, IEDB-SMM, SYFPEITHI, EPIJEN, NetMHC and ConvMHC). For each HLA-A molecule and for each protein, one to twenty 9-mer or 10-mer peptides with the highest score (highest affinity) as predicted by at least two tools were selected as candidate epitopes to be identified. The peptides were synthesized from China Peptides Co., Ltd with a purity above 95% defined by HPLC purification and mass spectrometry, and were used in cellular functional experiments. Lyophilized peptides were reconstituted at a stock concentration of 2mg/mL in DMSO-PBS solution.

### 4. DC-peptide-PBL co-culture experiment

Fresh PBMCs were suspended in serum-free RPMI 1640 and were allowed to adhere to culture flask for 2 hours in 5% CO_2_ at 37°C. Non-adherent cells (PBLs) were then removed and were cryopreserved at -80°C until further use. The resulting adherent cells were cultured in RPMI 1640 with 10% FCS, 1% penicillin/streptomycin, recombinant human GM-CSF (rhGM-CSF, 1000IU/mL, PrepoTech) and recombinant human IL-4 (rhIL-4, 500IU/mL, PrepoTech). At day 3 and day 5, half of the medium was replaced with fresh complete medium containing the cytokines with the same final concentration detailed above. At day 5, LPS (1μg/mL, sigma) was added to induce mature DCs (mDCs). At day 7, mature DCs (mDCs) were collected and were identified by flow cytometry (FACS Calibur, BD Bioscience) with FITC-labeled anti-CD83, anti-CD80, anti-CD86, anti-HLA-DR and PE-labeled anti-HLA-ABC and anti-CD1a, respectively. mDCs were incubated with single peptide (20μg/mL, which corresponding to the HLA-A allele of indicated healthy donor) in serum free RPMI 1640 in 48-well plate (5×10^4^ cells/well) for four hours in 5% CO_2_ at 37°C, then the PBLs from the same donor (recovered one day ago, and prelabeled with CFSE or not) were added into the well (1×10^6^ cells/well) for further 14-day co-culture. Recombinant human IL-2 (20 IU/mL) was added at day 11. At day 14, the corresponding peptide (20μg/mL) was added. At day 17, rhIL-2 was added again (10 IU/ mL). At day 21, cells were harvested and followed by ICS or T cell proliferation assay.

### 5. Intracellular IFN-γ staining of stimulated T cells

Cells from the DC-peptide-PBL co-cultures were harvested and coincubated with corresponding peptide (20μg/mL) or no peptide (negative control) for 16 hours in serum- free RPMI-1640 medium in 48-well plate at 37°C and 5% CO2. After that, BFA/Monensin mixture was added to the cells for another 6 hours culture. Cells were then harvested, washed, blocked with human FcR Blocking Reagent (MACS) for 20 min at 4°C and were stained with FITC-labeled anti-CD3 and APC-labeled anti-CD8 antibodies for 30 min at 4 °C. After washing, cells were fixed and permeabilized following the protocol and were further incubated with PE-anti-human IFN-γ (4S.B3) (BD) for another 30 min at 4°C followed by flow cytometry. The frequencies of IFN-γ^+^ cells in CD3^+^/CD8^+^ populations were calculated.

### 6. CD8^+^ T cell proliferation assay

In the DC-peptide-PBL co-culture, PBLs were pre-stained with CFSE. Briefly, PBLs were thawed, washed, and labeled with CFSE at a final concentration of 1.5μM for 20 min at 37°C. After washing, the CFSE-prelabeled PBLs were seeded into DC-peptide-PBL co-culture well and incubated for 14 days. At day 22, cells were harvested and blocked with human FcR Blocking Reagent (MACS) for 20 min, then stained with PE-labeled anti-CD3 and APC-labeled anti-CD8 antibodies for 30 min at 4°C for further analysis on the flow cytometry. The proliferation percentage of CD8^+^ T cells in CD3^+^/CD8^+^ population was analyzed according to the reduction of CFSE-staining brightness.

### 7. Generation of HMy2.CIR cell lines expressing indicated HLA-A molecule

Total mRNA was extracted from the PBMCs of the healthy donor with indicated HLA-A alleles, the cDNA of each HLA-A allele was amplified using PCR and followed by the routine construction of pcDNATM3.1/myc-His(-)A recombinant plasmid. After electrotransfection, the cell lines stably expressing indicated HLA-A molecule was screened by G418. Then the cell lines were stained with fluorescence-labeled monoclonal antibody W6/32 against HLA-ABC or anti-HLA-A24, the high-expression cells were then sorted using flow cytometry and followed by pure culture and sequencing analyses.

### 8. HLA-A molecule competitive binding assay

A set of plasmid-transfected HMy2.CIR cell lines expressing indicated HLA-A molecule were generated in house and sorted by flow cytometry. The cell lines were then used in the competitive peptide binding assay according to the references^45^. Briefly, the CIR cell lines expressing indicated HLA-A molecule were washed with acid buffer (0.131M citric acid and 0.061M sodium phosphate Na_2_HPO_4_, PH3.3, 0.22μm filtered) for 1 min, and then neutralized by IMDM medium containing 0.5%BSA. Cells were washed, seeded into 96-well U culture plate (1×10^5^ cells/100μL/well) with β_2_-m (1μg/mL). Then 25μL unlabeled peptide to be tested (5μM or 15μM) and 25μL fluorescent-labeled reference peptide (300nM) were added into the well and coincubated for 24h at 4°C. The reference peptides used in this research were FLPSDK(FITC)FPSV (for HLA-A0201, A0203 and A0206), YVNVNK(FITC)GLK (for HLA-A1101 and A3303), EYLVSK(FITC)GVW (for A2402), YLEPAK(FITC)AKY (for A0101) and ASRELK(FITC)VSY (for A3001). The plate was centrifuged at 600rpm for 5min at room temperature (RT). Cells were washed twice with 100μL cold 0.5% BSA-PBS. Finally, cells were resuspended with 150μL PBS and transferred to the flow tube and further analyzed with flow cytometry. Fluorescence polarization (FP) _sample is the FP value for the sample, while the minimum FP_ free is the FP value for free FITC-labeled reference peptide, and the maximum reaction FP_ no is the FP value for FITC-labeled reference peptide without unlabeled competitor peptide. Competitive binding (%) = [1-(FP_ sample-FP_ free) / (FP_ no -FP_ free)] × 100%. IC50 is the concentration of unlabeled peptide required to inhibit the binding of labeled reference peptide by 50%, which is calculated from the competitively binding inhibition (%) of the sample at 5μM and 15μM. Binding affinity of unlabeled peptide with indicated HLA-A molecule is assessed by IC50. IC50<5μM means high binding affinity, between 5-15μM means intermediate binding affinity, more than 15μM means low binding affinity or no binding affinity.

### 9. Preparation of peptide pools for vaccine immunization

Validated antigenic peptides restricted by HLA-A2 molecules (including A0201, A0203, A0206, A0207) were reconstituted in ideal solution before use at a final concentration of 5mg/mL for vaccine immunization and 2mg/mL for T cell response detection. Totally 31 antigenic peptides (9-mer or 10-mer) were grouped into four pools (pool-v1 to v4) for vaccine immunization (Table S3). For IFN-γ-ELISPOT assay, the 31 antigenic peptides were grouped into eight pools according to their derived protein and the feature of acid and alkalinity (Table S3). For IFN-γ ICS and ELISA, the 31 antigenic peptides were grouped into five pools according to their derived proteins (Table S3).

### 10. T2 cell binding assay

To assess the affinity of HLA-A2-restricted epitope peptides with HLA-A0201 molecule, peptide-induced up-regulation of HLA-A0201 expression on T2 cells was measured. Briefly, T2 cells were incubated with single peptide of the 31 epitopes (50μg/mL) or CMVpp65_495-503_ peptide (NLVPMVATV, 50μg/mL, as positive control) or OVA_257-264_ peptide (SIINFKEL, 50μg/mL, as negative control) or no peptide and 3μg/mL β2-m for 16 hours at 37 °C and 5% CO_2_. Then T2 cells were stained with PE-labeled anti-HLA-A2.1 antibody for 30 min at 4 °C followed by flow cytometry. The fluorescence index (FI) was calculated as follows: FI = (mean PE fluorescence with the given peptide - mean PE fluorescence without peptide)/ (mean PE without peptide). FI > 0.5 was the criteria of peptides with affinity while peptides with FI > 1 were regarded as high-affinity epitopes. FI ≤ 0.5 means low affinity or no binding.

### 11. Preparation of PLGA-NPs/peptides vaccine

Peptides-encapsulated PLGA-NPs were prepared freshly using the double-emulsion solvent evaporation method. To equal the amount of peptides in PLGA-NPs/peptides vaccine to the PolyI:C/peptides vaccine and R848/peptides vaccine, the loading efficiency of the PLGA-NPs was calculated before vaccination, then the PLGA-NPs carrying single peptide pool were prepared for future injections (one injection/mouse, 3 mice). Briefly, 60 mg of PLGA with or without single peptide pool (575μg/pool, 72-82 μg/peptide) was dissolved in 15mL of dichloromethane followed by ultrasonic dispersion for 30s at 40% amplitude to get the primary emulsion. Then, the primary emulsion was added into 150mL of 1% polyvinyl alcohol and sonicated for another 90 s to form the secondary emulsion. The resulting emulsion was added into 300mL of 0.5% PVA solution drop by drop with incessant magnetic stir to allow the evaporation of the dichloromethane. Four hours later, the solution was collected and centrifuged at 6,000 rpm for 5 min. The supernatant was harvested and was ultracentrifuged twice at 12,000 rpm for 10 min. The resulting PLGA-NPs were dispersed in deionized water and were further mixed with EDC and NHS solution for 1 h to allow surface activation of –COOH. After washing, the solution was added dropwise in 1% PEI with a magnetic stirrer for 4 hours at RT. Then, PEI-conjugated NPs were then collected and coincubated with single peptide pool (575μg/pool, 72-82 μg/peptide) in sterile PBS overnight at 4°C on a rotator. Finally, the peptides-encapsulated and - surface coupled PLGA-NPs/peptides vaccine was collected and preserved at 4 °C for further use.

### 12. Preparation of Poly I:C/peptides and R848/peptides vaccines and mice immunization

At day 0, mice were injected subcutaneously as the primary immunization. After that, booster immunizations were applied at day 7 and 21. At day 28, mice were executed for further study. The amount of each peptide for each inoculation was 10μg/mouse per time point, so, the amount of each peptide pool was 70 or 80μg/mouse/time point. Each mouse was inoculated with four peptide pools per time point. Each peptide pool was inoculated at one injection site (subcutaneously at tail root, back of the neck and around the groin). Twelve female HLA-A2/DR1 transgenic mice were randomly divided into four groups. The immunization groups, vaccines formula and vaccination scheme were described in Table S4.

### 13. ELISPOT and ICS

96-well PVDF-membrane microplates (Merck&Millipore) were coated with anti-IFN-γ capture monoclonal antibody (BD) at 4°C overnight, and were washed and blocked. Spleen cells (2×10^5^/100μL) of primed mice were added into each well, together with single peptide pool (2μg/well for each peptide), PHA (10μg/mL as positive controls), irrelevant epitope peptides (HLA-A2-restricted AFP_158-166_ and A24-restricted AFP_424-432_, 2μg/well for each peptide as no-specific control) or no peptide (negative control). After incubation for 20 h at 37 °C and 5% CO_2_, the plates were washed and then incubated with biotinylated anti-IFN-γ detecting antibody (BD) for 2h at RT. The plates were washed and then incubated with the streptavidin-conjugated HRP (BD) for 1h at RT. After washing the plates, AEC solution (BD) was used as the color developing agent and the developed spots were imaged and enumerated with professional plate reader.

Additionally, Spleen cells of primed mice were incubated with single peptide pool (20μg/mL for each peptide), PHA (10μg/mL), irrelevant epitope peptides (AFP_158-166_ and AFP_424-432_, 20μg/mL for each peptide) or no peptide for 16 hours in serum-free RPMI-1640 medium in 48-well plate at 37°C and 5% CO2. After that, BFA/Monensin mixture was added to the cells for another 6-hour culture. Cells were then harvested, washed, blocked with anti-mouse CD16/CD32 for 20 minutes at 4°C and were stained with FITC-labeled anti-CD3 and PE-labeled anti-CD8 antibodies for 30 min at 4 °C. After washing, cells were fixed and permeabilized following the protocol and were further incubated with APC-anti-mouse IFN-γ (XMG1.2) (BD) for another 30 min at 4°C followed by flow cytometry. The frequencies of IFN-γ^+^ cells in CD3^+^/CD8^+^ populations were calculated.

### 14. ELISA

Spleen cells were incubated with single peptide pool (16μg/mL for each peptide) or no peptide (negative control) in 48-well plate for 3 days at 37°C and 5% CO_2_. Then, the supernatants were collected for ELISA. The mouse IFN-γ detection ELISA kit (Dakewe, China) was used to quantify the IFN-γ in supernatants according to manufactor’s protocol.

### 15. Hematoxylin-eosin staining

28 days after primary immunization, the heart, liver, lung and kidney from executed mice were immersed in 4% paraformaldehyde overnight. After that, individual lobes of organs’ biopsy material were placed in processing cassettes, dehydrated through a serial alcohol gradient, and embedded in paraffin wax blocks. 5-μm-thick tissue sections were dewaxed in xylene, rehydrated through decreasing concentrations of ethanol, and washed in PBS. Then hematoxylin and eosin staining was carried out routinely.

## Acknowledgement

This work was supported by National Nature Science Foundation of China (82041006) and COVID-19 Emergency Research Fund of Zhejiang University of China (2020XGZX021). The sponsors had no role in study design, data collection and analysis, preparation of the manuscript, or decision to submit the article for publication.

## Author Contributions

C.S., Y.H. and G.Z. designed and supervised the research. X.J. and Y.D. performed the main experiments of this study. S.S. performed the transgenic mice experiments. X.W. and Z.Z. assisted in the PBMC preparation, cell cultures and flow cytometry and consequent data analysis. X.L. performed the HLA genotyping and assisted in the generation of HMy2.CIR cell lines expressing indicated HLA-A allotypes. A.S and Y.W assisted in the preparation of peptide cocktail vaccines and mice immunizations. M.L. and X.C collected healthy donors’ blood samples and separated the PBMCs. J.L. and B.L. assisted in the on-silicon prediction of epitopes and affinity analysis with HLA-A molecules. C.S. and X.J. wrote the manuscript with discussions from all authors. Y.H., J. Z and H.Q. analyzed and organized the whole data and assisted in the revision of manuscript.

Supplementary information is available at Cellular and Molecular Immunology’s website.

**Figure S1: Generation of mature mDC from adherent monocytes.** PBMCs from healthy donors were seeded into culture flask and the monocytes adhered for 2 h as described in the Methods section. After washing out the non-adherent cells in both systems, the cells were cultured for 5 days with 1,000 IU/mL GM-CSF and 500 IU/mL IL-4. Then the immature DCs were matured with 1μg/mL LPS for another 48 hours. Immature DCs and mature DCs were stained for CD1a, CD80, CD83, CD86, HLA-ABC and HLA-DR. The unstained and stained populations in the histograms are shown in grey and black, respectively. (A) Phenotype of immature DCs on day 5. (B) Phenotype of mature DCs on day 7.

**Figure S2: 120 epitopes were validated by DC-peptide-PBL co-culture experiments (IFN-γ ICS flow plots).** DCs were induced for 7 days from healthy donor’s PBMCs, then coincubated with candidate epitope peptides and autologous PBLs for 14 days. Cells were harvested and stimulated by corresponding candidate peptides for another 16 hours followed by IFN-γ ICS. The presented are flow plots for each positive epitope peptide. The data in horizontal coordinates mean the frequency of IFN-γ^+^ T cells in CD3^+^/CD8^+^ T cell population.

**Figure S3: 120 candidate epitopes were validated by DC-peptide-PBL co-culture experiments (CFSE staining flow plots).** DCs were coincubated with candidate epitope peptides and CFSE-prelabeled PBLs for 14 days. Cells were then analyzed by flow cytometry. The presented are flow plots for each positive epitope peptide. The data in horizontal coordinates mean the proliferation percentage of CD8^+^ T cells in CD3^+^/CD8^+^ T cell population.

**Figure S4: Eight kinds of HMy2.CIR cell lines expressing one indicated HLA-A allotype.** The transfected HMy2.CIR cell lines expressing HLA-A2402, A0203, A0201, A0206, A1101, A3303, A0101, or A3001 were generated, respectively, and then sorted by flow cytometry followed by pure culture and sequencing analyses. The purity of CIR-A2402 cells was 94.1% after being sorted with FITC-anti-A24 staining. The purity of CIR-A0203 cell was 84.3% after being sorted with PE-anti-HLA-ABC (W6/32) or PE-anti-HLA-A2.1 staining. The purities of CIR-A0201/0206/1101/3303/0101/3001 were all more than 80% after being sorted with PE-anti-HLA-ABC staining.

**Figure S5: Binding affinity of 120 validated epitopes with HLA-A allotypes as defined by HLA-A molecule competitive binding experiments.** A series of unlabeled epitope peptides of SRAS-CoV-2 were coincubated, at 5μM and 15μM respectively, with fluorescent-labeled reference peptides and CIR cell lines expressing the corresponding HLA-A molecules for 24 hours. Then the competitively binding inhibition (%) of the epitope peptide at 5μM and 15μM was calculated by measuring the CIR cells fluorescence strength. Shown are the histograms of two concentrations (5μM and 15μM). Black solid line was the histogram of 5μM test peptide; dotted line was the histogram of 15μM, test peptide; black filled line was the maximal fluorescence (FITC-labeled reference peptide without competitive peptides) while the lightest gray line was the negative control (background fluorescence with 1640 alone).

**Figure S6: Affinity of HLA-A2-restricted epitope peptides with HLA-A0201 molecule as detected by T2 cell binding assay.** T2 cells were incubated with single peptide of the 31 epitopes, or with CMVpp65_495-503_ peptide as positive control, OVA_257-264_ peptide as negative control, or no peptide and β2-m for 16 hours, then followed by PE-labeled anti-HLA-A2.1 antibody staining to test the up-regulation of HLA-A0201 molecules onto T2 cells. The fluorescence index (FI) was calculated with flow cytometry.

**Figure S7: The peptide cocktail vaccines have no visible toxicity on the organs.** Seven days after the last booster, all mice were executed. Heart, liver, lung and kidney were taken out, immersed and were finally stained with Hematoxylin-Eosin. No obvious pathological damage was found in all organs in all groups. The representative HE staining of heart, liver, lung and kidney in each mouse from four groups were exhibited in (A), (B), (C) and (D), respectively.

## References

1. Reche, P. A. Potential Cross-Reactive Immunity to SARS-CoV-2 From Common Human Pathogens and Vaccines. Front. Immunol. 11, 586984 (2020).

2. Chen, Z. et al. T and B cell Epitope analysis of SARS-CoV-2 S protein based on immunoinformatics and experimental research. J. Cell. Mol. Med. 25, 1274–1289 (2021).

3. Jeyanathan, M. et al. Immunological considerations for COVID-19 vaccine strategies. Nat. Rev. Immunol. 20, 615–632(2020).

4. Krammer, F. SARS-CoV-2 vaccines in development. Nature 586, 516–527 (2020).

5. Addetia, A. et al. Neutralizing Antibodies Correlate with Protection from SARS-CoV-2 in Humans during a Fishery Vessel Outbreak with a High Attack Rate. J. Clin. Microbiol. 58, e02107–02120 (2020).

6. Long, Q. X. et al. Clinical and immunological assessment of asymptomatic SARS-CoV-2 infections. Nat. Med. 26, 1200–1204 (2020).

7. Schmidt, M. E. et al. The CD8 T Cell Response to Respiratory Virus Infections. Front. Immunol. 9, 678 (2018).

8. Braun, J. et al. SARS-CoV-2-reactive T cells in healthy donors and patients with COVID-19. Nature 587, 270–274 (2020).

9. Grifoni, A. et al. Targets of T Cell Responses to SARS-CoV-2 Coronavirus in Humans with COVID-19 Disease and Unexposed Individuals. Cell 181, 1489–1501 (2020).

10. Le, B. N. et al. SARS-CoV-2-specific T cell immunity in cases of COVID-19 and SARS, and uninfected controls. Nature 584, 457–462 (2020).

11. Peng, Y. et al. Broad and strong memory CD4(+) and CD8(+) T cells induced by SARS-CoV-2 in UK convalescent individuals following COVID-19. Nat. Immunol. 21, 1336–1345 (2020).

12. Sekine, T. et al. Robust T Cell Immunity in Convalescent Individuals with Asymptomatic or Mild COVID-19. Cell 183, 158–168 (2020).

13. Xu, Z. et al. Pathological findings of COVID-19 associated with acute respiratory distress syndrome. The. Lancet. Respiratory. Medicine. 8, 420–422 (2020).

14. Wang, F. et al. Characteristics of Peripheral Lymphocyte Subset Alteration in COVID-19 Pneumonia. J. Infect. Dis. 221, 1762–1769 (2020).

15. Jiang, Y. et al. COVID-19 pneumonia: CD8(+) T and NK cells are decreased in number but compensatory increased in cytotoxic potential. Clin. Immunol. 218, 108516 (2020).

16. Rodriguez, L. et al. Systems-Level Immunomonitoring from Acute to Recovery Phase of Severe COVID-19. Cell. Rep. Med. 1, 100078 (2020).

17. Mathew, D. et al. Deep immune profiling of COVID-19 patients reveals distinct immunotypes with therapeutic implications. Science 369, 6508 (2020).

18. Meckiff, B. J. et al. Imbalance of Regulatory and Cytotoxic SARS-CoV-2-Reactive CD4(+) T Cells in COVID-19. Cell 183, 1340–1353 (2020).

19. Zhang, J. Y. et al. Single-cell landscape of immunological responses in patients with COVID-19. Nat. Immunol. 21, 1107–1118 (2020).

20. Sauer, K. et al. An Effective COVID-19 Vaccine Needs to Engage T Cells. Front. Immunol. 11, 581807 (2020).

21. Zhuang, Z. et al. Mapping and role of T cell response in SARS-CoV-2-infected mice. J. Exp. Med. 218, e20202187 (2021).

22. Anand, R. et al. Computational perspectives revealed prospective vaccine candidates from five structural proteins of novel SARS corona virus 2019 (SARS-CoV-2). PeerJ 8, e9855 (2020).

23. Saha, R. et al. Designing a next generation multi-epitope based peptide vaccine candidate against SARS-CoV-2 using computational approaches. 3. Biotech. 11, 47 (2021).

24. Singh, H. et al. Designing spike protein (S-Protein) based multi-epitope peptide vaccine against SARS COVID-19 by immunoinformatics. Heliyon 6, e05528 (2020).

25. Behmard, E. et al. Immunoinformatic design of a COVID-19 subunit vaccine using entire structural immunogenic epitopes of SARS-CoV-2. Sci. Rep. 10, 20864 (2020).

26. Banerjee, S., et al. Immuno-informatics approach for multi-epitope vaccine designing against SARS-CoV-2. bioRxiv https://www.biorxiv.org/content/10.1101/2020.07.23.218529v3 (2020).

27. Noorimotlagh, Z. et al. Immune and bioinformatics identification of T cell and B cell epitopes in the protein structure of SARS-CoV-2: A systematic review. Int. Immunopharmacol. 86, 106738 (2020).

28. Cun, Y. et al. COVID-19 coronavirus vaccine T cell epitope prediction analysis based on distributions of HLA class I loci (HLA-A, -B, -C) across global populations. Hum. Vaccin. Immunother. 17, 1097–1108 (2021).

29. Ferretti, A. K. et al. Unbiased Screens Show CD8+ T Cells of COVID-19 Patients Recognize Shared Epitopes in SARS-CoV-2 that Largely Reside outside the Spike Protein. Immunity 53, 1095–1107 (2020).

30. Mateus, J. G. et al. Selective and cross-reactive SARS-CoV-2 T cell epitopes in unexposed humans Science 370, 89–94 (2020).

31. Prachar, M. et al. Identification and validation of 174 COVID-19 vaccine candidate epitopes reveals low performance of common epitope prediction tools. Sci. Rep. 10, 20465 (2020).

32. Kared, H. et al. SARS-CoV-2-specific CD8+ T cell responses in convalescent COVID-19 individuals. J. Clin. Invest. 131, e145476 (2021).

33. Rydyznski, M. C. Antigen-Specific Adaptive Immunity to SARS-CoV-2 in Acute COVID-19 and Associations with Age and Disease Severity. et al. Cell 183, 996–1012 (2020).

34. Tang, F. et al. Lack of peripheral memory B cell responses in recovered patients with severe acute respiratory syndrome: a six-year follow-up study. J. Immunol. 186, 7264–7268 (2011)

35. Channappanavar, R. et al. Virus-specific memory CD8 T cells provide substantial protection from lethal severe acute respiratory syndrome coronavirus infection. J. Virol. 88, 11034–11044 (2014).

36. Alcazer, V., et al. Neoepitopes-based vaccines: challenges and perspectives. Eur. J. Cancer. 108, 55–60 (2019).

37. Sahin, U. T. Personalized vaccines for cancer immunotherapy. Science 359, 1355–1360 (2018).

38. Ott, P. A. et al. An immunogenic personal neoantigen vaccine for patients with melanoma. Nature 547, 217–221 (2017).

39. Sahin, U. et al. Personalized RNA mutanome vaccines mobilize poly-specific therapeutic immunity against cancer. Nature 547, 222–226 (2017).

40. Li, F. et al. Rapid tumor regression in an Asian lung cancer patient following personalized neo-epitope peptide vaccination. Oncoimmunology 5, e1238539 (2016).

41. Wang, B. et al. Identification of an HLA-A*0201-restricted CD8+ T-cell epitope SSp-1 of SARS-CoV spike protein. Blood 104, 200–206 (2004).

42. Lacey, S. F. et al. Characterization of immunologic properties of a second HLA-A2 epitope from a granule protease in CML patients and HLA-A2 transgenic mice. Blood 118, 2159–2169 (2011).

43. Takagi, A. et al. Identification of HLA-A*02:01-restricted candidate epitopes derived from the non-structural polyprotein 1a of SARS-CoV-2 that may be natural targets of CD8(+) T cell recognition in vivo. J. Virol. Dec 2, 01837–20 (2020).

44. Weiskopf, D., et al. Phenotype and kinetics of SARS-CoV-2-specific T cells in COVID-19 patients with acute respiratory distress syndrome. Sci. Immunol. 5, eabd2071 (2020).

45. Tsuboi, A. et al. Enhanced induction of human WT1-specific cytotoxic T lymphocytes with a 9-mer WT1 peptide modified at HLA-A*2402-binding residues. Cancer. Immunol. Immunother. 51, 614–620 (2002).

