## Supplemental tables and figures for "Screening of HLA-A restricted T cell epitopes of SARS-CoV-2 and induction of CD8^+^ T cell responses in HLA-A transgenic mice"

Table S1 270 SARS-CoV-2 T cell epitopes on-silicon predicted were synthesized for further validation

| Protein | Epitope No | HLA-A allele | Epitope sequence | Start Position | End position |
|---|---|---|---|---|---|
| E protein | A1 | HLA-A*02:01 | FLAFVVFLL | 20 | 28 |
| | A2 | HLA-A*02:01 | SLVKPSFYV | 50 | 58 |
| | A3 | HLA-A*02:01 | VLLFLAFVV | 17 | 25 |
| | A4 | HLA-A*02:01 | FLLVTLAIL | 26 | 34 |
| | A5 | HLA-A*02:01 | RLCAYCCNIV | 38 | 47 |
| | A6 | HLA-A*02:07 | SVLLFLAFV | 16 | 24 |
| | A7 | HLA-A*02:07 | LFLAFVVFLL | 19 | 28 |
| | A8 | HLA-A*02:06 | FVVFLLVTL | 23 | 31 |
| | A9 | HLA-A*02:06 | LIVNSVLLFL | 12 | 21 |
| | A10 | HLA-A*02:03 | FLAFVVFLLV | 20 | 29 |
| | A11 | HLA-A*02:03 | FVSEETGTL | 4 | 12 |
| | A12 | HLA-A*11:01 | NIVNVSLVK | 45 | 53 |
| | A13 | HLA-A*11:01 | VTLAILTALR | 29 | 38 |
| | A14 | HLA-A*11:01 | SLVKPSFYVY | 50 | 59 |
| | A15 | HLA-A*11:01 | SFYVYSRVK | 55 | 63 |
| | A16 | HLA-A*11:01 | TLAILTALR | 30 | 38 |
| | A17 | HLA-A*11:02 | LVKPSFYVY | 51 | 59 |
| | A18 | HLA-A*11:02 | RVKNLNSSR | 61 | 69 |
| | A19 | HLA-A*11:02 | VSLVKPSFY | 49 | 57 |
| | A20 | HLA-A*24:02 | VFLLVTLAI | 25 | 33 |
| | A21 | HLA-A*24:02 | VLLFLAFVVF | 17 | 26 |
| | A22 | HLA-A*24:02 | TLIVNSVLLF | 11 | 20 |
| | A23 | HLA-A*24:02 | VNSVLLFLAF | 14 | 23 |
| | A24 | HLA-A*33:03 | KPSFYVYSR | 53 | 61 |
| | A25 | HLA-A*30:01 | SSRVPDLLV | 67 | 75 |
| | A26 | HLA-A*30:01 | IVNSVLLFL | 13 | 21 |
| M protein | B1 | HLA-A*02:01 | GLMWLSYFI | 89 | 97 |
| | B2 | HLA-A*02:01 | KLLEQWNLV | 15 | 23 |
| | B3 | HLA-A*02:01 | FVLAAVYRI | 65 | 73 |
| | B4 | HLA-A*02:01 | FLFLTWICLL | 26 | 35 |
| | B5 | HLA-A*02:01 | LIFLWLLWPV | 51 | 60 |
| | B6 | HLA-A*02:01 | TLACFVLAAV | 61 | 70 |
| | B7 | HLA-A*02:01 | FLWLLWPVTL | 53 | 62 |
| | B8 | HLA-A*02:07 | SMWSFNPET | 108 | 116 |
| | B9 | HLA-A*02:07 | LLWPVTLAC | 56 | 64 |
| | B10 | HLA-A*02:07 | FLYIIKLIFL | 45 | 54 |
| | B11 | HLA-A*02:06 | IAMACLVGL | 82 | 90 |
| | B12 | HLA-A*02:06 | VTLACFVLA | 60 | 68 |

| | | | | | |
|---|---|---|---|---|---|
| | B13 | HLA-A*02:06 | LVIGFLFLT | 22 | 30 |
| | B14 | HLA-A*02:06 | YIIKLIFLWL | 47 | 56 |
| | B15 | HLA-A*02:03 | FIASFRLFA | 96 | 104 |
| | B16 | HLA-A*02:03 | ILRGHLRIA | 144 | 152 |
| | B17 | HLA-A*02:03 | AMACLVGLM | 83 | 91 |
| | B18 | HLA-A*11:01 | ATSRTLSYYK | 171 | 180 |
| | B19 | HLA-A*11:01 | GTITVEELK | 6 | 14 |
| | B20 | HLA-A*11:01 | LSYFIASFR | 93 | 101 |
| | B21 | HLA-A*11:01 | YSRYRIGNYK | 196 | 205 |
| | B22 | HLA-A*11:01 | AVILRGHLR | 142 | 150 |
| | B23 | HLA-A*11:01 | LVIGAVILR | 138 | 146 |
| | B24 | HLA-A*11:01 | FIASFRLFAR | 96 | 105 |
| | B25 | HLA-A*11:02 | RIAGHHLGR | 150 | 158 |
| | B26 | HLA-A*24:02 | SYFIASFRLF | 94 | 103 |
| | B27 | HLA-A*24:02 | MWLSYFIASF | 91 | 100 |
| | B28 | HLA-A*24:02 | LYIIKLIFLW | 46 | 55 |
| | B29 | HLA-A*24:02 | QWNLVIGFLF | 19 | 28 |
| | B30 | HLA-A*24:02 | LWPVTLACF | 57 | 65 |
| | B31 | HLA-A*24:02 | RFLYIIKLIF | 44 | 53 |
| | B32 | HLA-A*33:03 | IASFRLFAR | 97 | 105 |
| | B33 | HLA-A*33:03 | QFAYANRNR | 36 | 44 |
| | B34 | HLA-A*33:03 | SFRLFARTR | 99 | 107 |
| | B35 | HLA-A*33:03 | YYKLGASQR | 178 | 186 |
| | B36 | HLA-A*30:01 | RTRSMWSFN | 105 | 113 |
| | B37 | HLA-A*30:01 | RNRFLYIIK | 42 | 50 |
| | B38 | HLA-A*30:01 | TSRTLSYYK | 172 | 180 |
| | B39 | HLA-A*30:01 | ANRNRFLYI | 40 | 48 |
| | B40 | HLA-A*30:01 | RYRIGNYKL | 198 | 206 |
| | B41 | HLA-A*30:01 | HLRIAGHHL | 148 | 156 |
| | B42 | HLA-A*30:01 | RVAGDSGFA | 186 | 194 |
| N protein | C1 | HLA-A*02:01 | LLLDRLNQL | 222 | 230 |
| | C2 | HLA-A*02:01 | GMSRIGMEV | 316 | 324 |
| | C3 | HLA-A*02:01 | WLTYTGAIKL | 330 | 339 |
| | C4 | HLA-A*02:01 | YLGTGPEAGL | 112 | 121 |
| | C5 | HLA-A*02:01 | ALALLLLDRL | 218 | 227 |
| | C6 | HLA-A*02:01 | IIWVATEGA | 130 | 138 |
| | C7 | HLA-A*02:01 | ILLNKHIDA | 351 | 359 |
| | C8 | HLA-A*02:07 | DLDDFSKQL | 399 | 407 |
| | C9 | HLA-A*02:07 | KLDDKDPNF | 338 | 346 |
| | C10 | HLA-A*02:06 | QTVTLLPAA | 390 | 398 |
| | C11 | HLA-A*02:06 | LQLPQGTTL | 159 | 167 |
| | C12 | HLA-A*02:06 | NTASWFTAL | 48 | 56 |
| | C13 | HLA-A*02:06 | TTLPKGFYA | 165 | 173 |

| | | | | | |
|---|---|---|---|---|---|
| | C14 | HLA-A*02:06 | RTATKAYNV | 262 | 270 |
| | C15 | HLA-A*02:06 | QIAQFAPSA | 303 | 311 |
| | C16 | HLA-A*02:06 | LALLLLDRL | 219 | 227 |
| | C17 | HLA-A*02:03 | QLQQSMSSA | 406 | 414 |
| | C18 | HLA-A*02:03 | RMAGNGGDA | 209 | 217 |
| | C19 | HLA-A*02:03 | SAFFGMSRI | 312 | 320 |
| | C20 | HLA-A*11:01 | ASAFFGMSR | 311 | 319 |
| | C21 | HLA-A*11:01 | ATEGALNTPK | 134 | 143 |
| | C22 | HLA-A*11:01 | KSAAEASKK | 249 | 257 |
| | C23 | HLA-A*11:01 | FTALTQHGK | 53 | 61 |
| | C24 | HLA-A*11:01 | QLPQGTTLPK | 160 | 169 |
| | C25 | HLA-A*11:01 | LLNKHIDAYK | 352 | 361 |
| | C26 | HLA-A*11:01 | AGLPYGANK | 119 | 127 |
| | C27 | HLA-A*11:01 | QQQGQTVTK | 240 | 248 |
| | C28 | HLA-A*11:01 | VTPSGTWLTY | 324 | 333 |
| | C29 | HLA-A*11:02 | KTFPPTEPK | 361 | 369 |
| | C30 | HLA-A*24:02 | YYRRATRRI | 86 | 94 |
| | C31 | HLA-A*24:02 | QFAPSASAFF | 306 | 315 |
| | C32 | HLA-A*24:02 | KHIDAYKTF | 355 | 363 |
| | C33 | HLA-A*24:02 | TWLTYTGAI | 329 | 337 |
| | C34 | HLA-A*24:02 | GYYRRATRRI | 85 | 94 |
| | C35 | HLA-A*24:02 | LSPRWYFYYL | 104 | 113 |
| | C36 | HLA-A*30:01 | RSRNSSRNS | 189 | 197 |
| | C37 | HLA-A*30:01 | GTRNPANNA | 147 | 155 |
| | C38 | HLA-A*30:01 | SSRGTSPAR | 201 | 209 |
| | C39 | HLA-A*30:01 | RSKQRRPQG | 36 | 44 |
| | C40 | HLA-A*30:01 | LIRQGTDYK | 291 | 299 |
| | C41 | HLA-A*30:01 | SSRNSTPGS | 193 | 201 |
| | C42 | HLA-A*30:01 | SSRSSSRSR | 183 | 191 |
| | C43 | HLA-A*30:01 | RQKRTATKA | 259 | 267 |
| | C44 | HLA-A*30:01 | SSRSRNSSR | 187 | 195 |
| | C45 | HLA-A*33:03 | NVTQAFGRR | 269 | 277 |
| | C46 | HLA-A*33:03 | IGYYRRATR | 84 | 92 |
| | C47 | HLA-A*33:03 | NTPKDHIGTR | 140 | 149 |
| | C48 | HLA-A*33:03 | QASSRSSSR | 181 | 189 |
| | C49 | HLA-A*11:02 | YKTFPPTEPK | 360 | 369 |
| S protein | D1 | HLA-A*02:01 | YLQPRTFLL | 269 | 277 |
| | D2 | HLA-A*02:01 | FIAGLIAIV | 1220 | 1228 |
| | D3 | HLA-A*02:01 | ELLHAPATV | 516 | 524 |
| | D4 | HLA-A*02:01 | SIIAYTMSL | 691 | 699 |
| | D5 | HLA-A*02:01 | KLNDLCFTNV | 386 | 395 |
| | D6 | HLA-A*02:01 | RLDKVEAEV | 983 | 991 |
| | D7 | HLA-A*02:01 | VLNDILSRL | 976 | 984 |

| | | | | | |
|---|---|---|---|---|---|
| | D8 | HLA-A*02:01 | FTISVTTEI | 718 | 726 |
| | D9 | HLA-A*02:01 | LLFNKVTLA | 821 | 829 |
| | D10 | HLA-A*02:01 | KIADYNYKL | 417 | 425 |
| | D11 | HLA-A*02:01 | VVFLHVTYV | 1060 | 1068 |
| | D12 | HLA-A*02:01 | FVFLVLLPLV | 2 | 11 |
| | D13 | HLA-A*02:01 | MIAQYTSAL | 869 | 877 |
| | D14 | HLA-A*02:01 | GLIAIVMVTI | 1223 | 1232 |
| | D15 | HLA-A*02:01 | SVTTEILPV | 721 | 729 |
| | D16 | HLA-A*02:07 | KLPDDFTGCV | 424 | 433 |
| | D17 | HLA-A*02:07 | RLQSLQTYV | 1000 | 1008 |
| | D18 | HLA-A*02:07 | LLPLVSSQCV | 7 | 16 |
| | D19 | HLA-A*02:06 | AVDCALDPL | 288 | 296 |
| | D20 | HLA-A*02:06 | KQLSSNFGA | 964 | 972 |
| | D21 | HLA-A*02:06 | FQFCNDPFL | 133 | 141 |
| | D22 | HLA-A*02:06 | MQMAYRFNGI | 900 | 909 |
| | D23 | HLA-A*02:06 | YQDVNCTEV | 612 | 620 |
| | D24 | HLA-A*02:06 | KQIYKTPPI | 786 | 794 |
| | D25 | HLA-A*02:06 | LQIPFAMQM | 894 | 902 |
| | D26 | HLA-A*02:06 | LQSYGFQPT | 492 | 500 |
| | D27 | HLA-A*02:06 | NTQEVFAQV | 777 | 785 |
| | D28 | HLA-A*02:06 | TQLNRALTGI | 761 | 770 |
| | D29 | HLA-A*02:03 | HLMSFPQSA | 1048 | 1056 |
| | D30 | HLA-A*02:03 | FLHVTYVPA | 1062 | 1070 |
| | D31 | HLA-A*02:03 | QLNRALTGI | 762 | 770 |
| | D32 | HLA-A*02:03 | FKIYSKHTPI | 201 | 210 |
| | D33 | HLA-A*02:03 | FVSNGTHWFV | 1095 | 1104 |
| | D34 | HLA-A*11:01 | GVYFASTEK | 89 | 97 |
| | D35 | HLA-A*11:01 | MTSCCSCLK | 1237 | 1245 |
| | D36 | HLA-A*11:01 | RLFRKSNLK | 454 | 462 |
| | D37 | HLA-A*11:01 | SSTASALGK | 939 | 947 |
| | D38 | HLA-A*11:01 | NSASFSTFK | 370 | 378 |
| | D39 | HLA-A*11:01 | VTLADAGFIK | 826 | 835 |
| | D40 | HLA-A*11:01 | CTLKSFTVEK | 301 | 310 |
| | D41 | HLA-A*11:01 | SLIDLQELGK | 1196 | 1205 |
| | D42 | HLA-A*11:01 | FIEDLLFNK | 817 | 825 |
| | D43 | HLA-A*11:01 | EILPVSMTK | 725 | 733 |
| | D44 | HLA-A*11:01 | AQALNTLVK | 956 | 964 |
| | D45 | HLA-A*11:01 | QIYKTPPIK | 787 | 795 |
| | D46 | HLA-A*11:02 | VTYVPAQEK | 1065 | 1073 |
| | D47 | HLA-A*11:02 | ASANLAATK | 1020 | 1028 |
| | D48 | HLA-A*11:02 | GTHWFVTQR | 1099 | 1107 |
| | D49 | HLA-A*11:02 | GVLTESNKK | 550 | 558 |
| | D50 | HLA-A*11:02 | GVYYHKNNK | 142 | 150 |

| | | | | | |
|---|---|---|---|---|---|
| | D51 | HLA-A*24:02 | QYIKWPWYI | 1208 | 1216 |
| | D52 | HLA-A*24:02 | VYAWNRKRI | 350 | 358 |
| | D53 | HLA-A*24:02 | NYNYLYRLF | 448 | 456 |
| | D54 | HLA-A*24:02 | VYSTGSNVF | 635 | 643 |
| | D55 | HLA-A*24:02 | PYRVVVLSF | 507 | 515 |
| | D56 | HLA-A*24:02 | LYNSASFSTF | 368 | 377 |
| | D57 | HLA-A*24:02 | VYSSANNCTF | 159 | 168 |
| | D58 | HLA-A*24:02 | CYFPLQSYGF | 488 | 497 |
| | D59 | HLA-A*24:02 | PFAMQMAYRF | 897 | 906 |
| | D60 | HLA-A*24:02 | TYVPAQEKNF | 1066 | 1075 |
| | D61 | HLA-A*24:02 | IYSKHTPINL | 203 | 212 |
| | D62 | HLA-A*24:02 | IYKTPPIKDF | 788 | 797 |
| | D63 | HLA-A*24:02 | TFEYVSQPF | 167 | 175 |
| | D64 | HLA-A*24:02 | CFTNVYADSF | 391 | 400 |
| | D65 | HLA-A*30:01 | ATRFASVYA | 344 | 352 |
| | D66 | HLA-A*30:01 | ITRFQTLLA | 235 | 243 |
| | D67 | HLA-A*30:01 | TTRTQLPPA | 19 | 27 |
| | D68 | HLA-A*30:01 | KCYGVSPTK | 378 | 386 |
| | D69 | HLA-A*30:01 | AYRFNGIGV | 903 | 911 |
| | D70 | HLA-A*30:01 | RKRISNCVA | 355 | 363 |
| | D71 | HLA-A*30:01 | KNLREFVFK | 187 | 195 |
| | D72 | HLA-A*30:01 | ASVYAWNRK | 348 | 356 |
| | D73 | HLA-A*30:01 | RARSVASQSI | 683 | 692 |
| | D74 | HLA-A*30:01 | GTKRFDNPV | 75 | 83 |
| | D75 | HLA-A*30:01 | HVSGTNGTK | 69 | 77 |
| | D76 | HLA-A*33:03 | SVYAWNRKR | 349 | 357 |
| | D77 | HLA-A*33:03 | VYYPDKVFR | 36 | 44 |
| | D78 | HLA-A*33:03 | QTNSPRRAR | 677 | 685 |
| | D79 | HLA-A*33:03 | NVYADSFVIR | 394 | 403 |
| | D80 | HLA-A*33:03 | YYVGYLQPR | 265 | 273 |
| | D81 | HLA-A*33:03 | GIYQTSNFR | 311 | 319 |
| | D82 | HLA-A*33:03 | NGVGYQPYR | 501 | 509 |
| | D83 | HLA-A*33:03 | STGSNVFQTR | 637 | 646 |
| RdRp protein | R1 | HLA-A*02:01 | LLMPILTLT | 240 | 248 |
| | R2 | HLA-A*02:01 | TMADLVYAL | 123 | 131 |
| | R3 | HLA-A*02:01 | LMIERFVSL | 854 | 862 |
| | R4 | HLA-A*02:01 | AMRNAGIVGV | 195 | 204 |
| | R5 | HLA-A*02:01 | SLAIDAYPL | 861 | 869 |
| | R6 | HLA-A*02:01 | NLLKDCPAV | 88 | 96 |
| | R7 | HLA-A*02:01 | NLIDSYFVV | 64 | 72 |
| | R8 | HLA-A*02:01 | FVNEFYAYL | 741 | 749 |
| | R9 | HLA-A*02:01 | ILHCANFNV | 307 | 315 |
| | R10 | HLA-A*02:01 | KIFVDGVPFV | 332 | 341 |

| | | | | | |
|---|---|---|---|---|---|
| | R11 | HLA-A*02:01 | VMCGGSLYV | 667 | 675 |
| | R12 | HLA-A*02:01 | MLDMYSVML | 899 | 907 |
| | R13 | HLA-A*02:01 | NMLRIMASL | 628 | 636 |
| | R14 | HLA-A*02:01 | RLANECAQV | 654 | 662 |
| | R15 | HLA-A*02:01 | QLLFVVEVV | 468 | 476 |
| | R16 | HLA-A*02:07 | FVDGVPFVV | 334 | 342 |
| | R17 | HLA-A*02:07 | YLPYPDPSRI | 828 | 837 |
| | R18 | HLA-A*02:07 | FPPTSFGPLV | 321 | 330 |
| | R19 | HLA-A*02:06 | RQLLFVVEV | 467 | 475 |
| | R20 | HLA-A*02:06 | RILGAGCFV | 836 | 844 |
| | R21 | HLA-A*02:06 | SVAALTNNV | 397 | 405 |
| | R22 | HLA-A*02:06 | MILSDDAVV | 756 | 764 |
| | R23 | HLA-A*02:06 | LSFKELLVYA | 366 | 375 |
| | R24 | HLA-A*02:03 | MLKTVYSDV | 601 | 609 |
| | R25 | HLA-A*02:03 | MLRIMASLV | 629 | 637 |
| | R26 | HLA-A*02:03 | SLSHRFYRL | 647 | 655 |
| | R27 | HLA-A*02:03 | AMYTPHTVL | 923 | 931 |
| | R28 | HLA-A*02:03 | SIAATRGATV | 578 | 587 |
| | R29 | HLA-A*02:03 | LLSTDGNKI | 707 | 715 |
| | R30 | HLA-A*11:01 | ASGNLLLDK | 383 | 391 |
| | R31 | HLA-A*11:01 | TSFGPLVRK | 324 | 332 |
| | R32 | HLA-A*11:01 | KSAGFPFNK | 500 | 508 |
| | R33 | HLA-A*11:01 | KVAGFAKFLK | 41 | 50 |
| | R34 | HLA-A*11:01 | MTNRQFHQK | 566 | 574 |
| | R35 | HLA-A*11:01 | AVAKHDFFK | 95 | 103 |
| | R36 | HLA-A*11:01 | AIDAYPLTK | 863 | 871 |
| | R37 | HLA-A*11:01 | LVASIKNFK | 775 | 783 |
| | R38 | HLA-A*11:01 | VVSTGYHFR | 341 | 349 |
| | R39 | HLA-A*11:01 | TVKPGNFNK | 409 | 417 |
| | R40 | HLA-A*11:01 | KTNCCRFQEK | 50 | 59 |
| | R41 | HLA-A*11:01 | AISDYDYYR | 449 | 457 |
| | R42 | HLA-A*11:01 | CSQHTMLVK | 813 | 821 |
| | R43 | HLA-A*11:01 | CSLSHRFYR | 646 | 654 |
| | R44 | HLA-A*11:01 | GTSTDVVYR | 25 | 33 |
| | R45 | HLA-A*11:02 | ATVVIGTSK | 585 | 593 |
| | R46 | HLA-A*11:02 | KLFDRYFKY | 281 | 289 |
| | R47 | HLA-A*24:02 | SYYSLLMPI | 236 | 244 |
| | R48 | HLA-A*24:02 | SYFVVKRHTF | 68 | 77 |
| | R49 | HLA-A*24:02 | AYANSVFNI | 688 | 696 |
| | R50 | HLA-A*24:02 | YFNKKDWYDF | 156 | 165 |
| | R51 | HLA-A*24:02 | FYGGWHNML | 594 | 602 |
| | R52 | HLA-A*24:02 | IYNDKVAGF | 37 | 45 |
| | R53 | HLA-A*24:02 | RYNLPTMCDI | 457 | 466 |

| | | | | | |
|---|---|---|---|---|---|
| | R54 | HLA-A*24:02 | KYVRNLQHRL | 718 | 727 |
| | R55 | HLA-A*24:02 | NFNKDFYDF | 414 | 422 |
| | R56 | HLA-A*24:02 | EYADVFHLYL | 876 | 885 |
| | R57 | HLA-A*24:02 | FYAYLRKHF | 745 | 753 |
| | R58 | HLA-A*24:02 | TYHPNCVNCL | 293 | 302 |
| | R59 | HLA-A*24:02 | LYLQYIRKL | 883 | 891 |
| | R60 | HLA-A*30:01 | ATRGATVVI | 581 | 589 |
| | R61 | HLA-A*30:01 | RLKLFDRYFK | 279 | 288 |
| | R62 | HLA-A*30:01 | RVRQALLKT | 181 | 189 |
| | R63 | HLA-A*30:01 | RQFHQKLLK | 569 | 577 |
| | R64 | HLA-A*30:01 | HISRQRLTK | 113 | 121 |
| | R65 | HLA-A*30:01 | KARLYYDSM | 511 | 519 |
| | R66 | HLA-A*33:03 | DFYDFAVSK | 418 | 426 |
| | R67 | HLA-A*33:03 | MVPHISRQR | 110 | 118 |
| | R68 | HLA-A*33:03 | LLKSIAATR | 575 | 583 |
| | R69 | HLA-A*33:03 | DALFAYTKR | 525 | 533 |
| | R70 | HLA-A*33:03 | RVCGVSAAR | 10 | 18 |

Table S2: Homologous analyses of 120 SARS-CoV-2 CD8+ T cell epitopes with SARS-CoV, common-cold HCoVs and variants of SARS-CoV-2.

| Protein | Epitope | SARS-CoV-2 | SARS-CoV | OC43 | NL63 | HKU1 | 229E |
|---|---|---|---|---|---|---|---|
| E | A1 | FLAFVVFLL | FLAFVVFLL | | | | |
| | A3 | VLLFLAFVV | VLLFLAFVV | | | | |
| | A4 | FLLVTLAIL | FLLVTLAIL | | | | |
| | A5 | RLCAYCCNIV | RLCAYCCNIV | | | | |
| | A6 | SVLLFLAFV | SVLLFLAFV | | | | |
| | A7 | LFLAFVVFLL | LFLAFVVFLL | | | | |
| | A9 | LIVNSVLLFL | LIVNSVLLFL | | | | |
| | A10 | FLAFVVFLLV | FLAFVVFLLV | | | | |
| | A12 | NIVNVSLVK | NIVNVSLVK | | | | |
| | A16 | TLAILTALR | TLAILTALR | | | | |
| | A18 | RVKNLNSSR | RVKNLNSSE | | | | |
| | A19 | VSLVKPSFY | VSLVKPTVY | | | | |
| | A20 | VFLLVTLAI | VFLLVTLAI | | | | |
| | A21 | VLLFLAFVVF | VLLFLAFVVF | | | | |
| | A22 | TLIVNSVLLF | TLIVNSVLLF | | | | |
| | A23 | VNSVLLFLAF | VNSVLLFLAF | | | | |
| | A25 | SSRVPDLLV | SSEGVPDLLV | | | | |
| | A26 | IVNSVLLFL | IVNSVLLFL | | | | |
| M | B1 | GLMWLSYFI | GLMWLSYFV | | | | |
| | B2 | KLLEQWNLV | QLLEQWNLV | | | | |
| | B3 | FVLAAVYRI | FVLAAVYRI | | | | |
| | B4 | FLFLTWICLL | FLFLAWIMLL | | | | |
| | B6 | TLACFVLAAV | TLACFVLAAV | | | | |
| | B10 | FLYIIKLIFL | FLYIIKLVFL | | | | |
| | B11 | IAMACLVGL | IAMACIVGL | | | | |
| | B12 | VTLACFVLA | VTLACFVLA | | | | |
| | B15 | FIASFRLFA | FVASFRLFA | | | | |
| | B16 | ILRGHLRIA | IIRGHLRMA | | | | |
| | B17 | AMACLVGLM | AMACIVGLM | | | | |
| | B18 | ATSRTLSYYK | ATSRTLSYYK | | | | |
| | B20 | LSYFIASFR | LSYFVASFR | | | | |
| | B21 | YSRYRIGNYK | YNRYRIGNYK | | | | |
| | B23 | LVIGAVILR | LVIGAVIIR | | | | |
| | B26 | SYFIASFRLF | SYFVASFRLF | | | | |
| | B28 | LYIIKLIFLW | LYIIKLVFLW | | | | |
| | B29 | QWNLVIGFLF | QWNLVIGFLF | | | | |
| | B30 | LWPVTLACF | LWPVTLACF | | | | |
| | B31 | RFLYIIKLIF | RFLYIIKLVF | | | | |
| | B34 | SFRLFARTR | SFRLFARTR | | | | SFRLFRRAR |

| | | | | | | | |
|---|---|---|---|---|---|---|---|
| | B35 | YYKLGASQR | YYKLGASQR | | | | |
| | B36 | RTRSMWSFN | RTRSMWSFN | RTGSFWSFN | | RTGSWWSFN | |
| | B37 | RNRFLYIIK | RNRFLYIIK | | | | |
| | B38 | ANRNRFLYI | SNRNRFLYI | | | | |
| | B40 | RYRIGNYKL | RYRIGNYKL | | | | |
| | B41 | HLRIAGHHL | HLRMAGHSL | | | | |
| N | C1 | LLLDRLNQL | LLLDRLNQL | | | | |
| | C3 | WLTYTGAIKL | WLTYHGAIKL | | | | |
| | C10 | QTVTLLPAA | PTVTLLPAA | | | | |
| | C12 | NTASWFTAL | NTASWFTAL | | | | |
| | C16 | LALLLLDRL | LALLLLDRL | | | | |
| | C17 | QLQQSMSSA | QLQNSMSGA | | | | |
| | C27 | QQQGQTVTK | QQQGQTVTK | | | | |
| | C35 | LSPRWYFYYL | LSPRWYFYYL | LLPRWYFYYL | | LLPRWYFYYL | |
| | C45 | NVTQAFGRR | NVTQAFGRR | | | | NVTQCFGPR |
| | C46 | IGYYRRATR | IGYYRRATR | | | | |
| | C47 | NTPKDHIGTR | NTPKDHIGTR | | | | |
| | C49 | YKTFPPTEPK | YKTFPPTEPK | | | | |
| S | D2 | FIAGLIAIV | FIAGLIAIV | | | | |
| | D5 | KLNDLCFTNV | KLNDLCFSNV | | | | |
| | D6 | RLDKVEAEV | RLDKVEAEV | | | | |
| | D12 | FVFLVLLPLV | | | | | |
| | D13 | MIAQYTSAL | MIAAYTAAL | | | | |
| | D17 | RLQSLQTYV | RLQSLQTYV | | | | |
| | D26 | LQSYGFQPT | | | | | |
| | D30 | FLHVTYVPA | FLHVTYVPS | | | | |
| | D31 | QLNRALTGI | QLNRALSGI | | | | |
| | D32 | FKIYSKHTPI | | | | | |
| | D33 | FVSNGTHWFV | | | | | |
| | D34 | GVYFASTEK | GIYFAATEK | | | | |
| | D38 | NSASFSTFK | NSTFFSTFK | | | | |
| | D40 | CTLKSFTVEK | | | | | |
| | D41 | SLIDLQELGK | SLIDLQELGK | | | | |
| | D42 | FIEDLLFNK | FIEDLLFNK | AIEDLLFDK | | FFEDLLFDK | |
| | D46 | VTYVPAQEK | VTYVPSQER | | | | |
| | D47 | ASANLAATK | ASANLAATK | | | | |
| | D48 | GTHWFVTQR | GTSWFITQR | | | | |
| | D50 | GVYYHKNNK | | | | | |
| | D52 | VYAWNRKRI | VYAWERKKI | | | | |
| | D53 | NYNYLYRLF | | | | | |
| | D55 | PYRVVVLSF | PYRVVVLSF | | | | |
| | D56 | LYNSASFSTF | LYNSTFFSTF | | | | |
| | D62 | IYKTPPIKDF | | | | | |

| | | | | | | | |
|---|---|---|---|---|---|---|---|
| | D64 | CFTNVYADSF | CFSNVYADSF | | | | |
| | D65 | ATRFASVYA | ATKFPSVYA | | | | |
| | D71 | KNLREFVFK | KHLREFVFK | | | | |
| | D72 | ASVYAWNRK | PSVYAWERK | | | | |
| | D76 | SVYAWNRKR | SVYAWERKK | | | | |
| | D77 | VYYPDKVFR | VYYPDEIFR | | | | |
| | D78 | QTNSPRRAR | | | | | |
| | D79 | NVYADSFVIR | NVYADSFVVK | | | | |
| | D80 | YYVGYLQPR | | | | | |
| | D81 | GIYQTSNFR | GIYQTSNFR | | | | |
| | D82 | NGVGYQPYR | TGIGYQPYR | | | | |
| RdRp | R4 | AMRNAGIVGV | AMRDAGIVGV | | | | |
| | R5 | SLAIDAYPL | SLAIDAYPL | SLAIDAYPL | SLAIDAYPL | SLAIDAYPL | SLAIDAYPL |
| | R6 | NLLKDCPAV | NLVKDCPAV | | | | NLLKGCNAV |
| | R8 | FVNEFYAYL | FVDEFYAYL | | | | |
| | R9 | ILHCANFNV | ILHCANFNV | IIHCANFNI | | IIHCANFNI | ILHCSNFNT |
| | R10 | KIFVDGVPFV | KIFVDGVPFV | QIFVDGVPFV | | QIFVDGVPFV | |
| | R11 | VMCGGSLYV | VMCGGSLYV | VMCGGCYYV | | VMCGGCYYV | |
| | R12 | MLDMYSVML | MLDMYSVML | | | | |
| | R13 | NMLRIMASL | NMLRIMASL | | | | |
| | R14 | RLANECAQV | RLANECAQV | RLANECAQV | RLGNELAQV | RLANECAQV | RLSNELAQV |
| | R15 | QLLFVVEVV | QLLFVVEVV | QLLFVLEVV | | QLLFVLEVV | |
| | R17 | YLPYPDPSRI | YLPYPDPSRI | YLPYPNPSRI | YLPYPDPSRI | YLPYPDPSRI | YLPYPDPSRI |
| | R23 | LSFKELLVYA | LSFKELLVYA | | | | |
| | R24 | MLKTVYSDV | MLKTVYSDV | | | | |
| | R30 | ASGNLLLDK | ASGNLLLDK | | | | |
| | R32 | KSAGFPFNK | KSAGFPFNK | KSAGYPFNK | KSAGWPLNK | KSAGYPFNK | KSAGWPLNK |
| | R34 | MTNRQFHQK | MTNRQFHQK | MTGRMFHQK | MTTRQYHQK | MTGRMFHQK | MTTRQFHQK |
| | R35 | AVAKHDFFK | AVAVHDFFK | | | | AVAKHDFFT |
| | R38 | VVSTGYHFR | VVSTGYHFR | | | | |
| | R39 | TVKPGNFNK | TVKPGNFNK | TVKPGNFNQ | | TVKPGNFNQ | TVKPGHFNK |
| | R40 | KTNCCRFQEK | KTNCCRFQEK | | | | |
| | R41 | AISDYDYYR | AISDYDYYR | | | | AIKDFDYYR |
| | R42 | CSQHTMLVK | CSQHTMLVK | CSQHTMLVK | | CSQHTMLVK | |
| | R43 | CSLSHRFYR | CNLSHRFYR | CSQSDRFYR | | | |
| | R44 | GTSTDVVYR | GTSTDVVYR | | | | |
| | R47 | SYYSLLMPI | SYYSLLMPI | | SYYSYMMPI | | |
| | R48 | SYFVVKRHTF | SYFVVKRHTM | | | | |
| Total | | 120 | 110 | 15 | 6 | 14 | 12 |
| Protein | Epitope | SARS-CoV-2 | B.1.1.7 | B.1.351 | P.1 | B.1.617 | Denmark variant |
| S | D50 | GVYYHKNNK | GVYHKNNK | GVYYHKNNK | GVYYHKNNK | GVYYHKNNK | GVYYHKNNK |
| | D53 | NYNYLYRLF | NYNYLYRLF | NYNYLYRLF | NYNYLYRLF | NYNYLYRLF | NYNYLFRLF |

| | | | | | | | |
|---|---|---|---|---|---|---|---|
| | D78 | QTNSPRRAR | QTNSPRRAR | QTNSPRRAR | QTNSPRRAR | QTNSRRRAR | QTNSPRRAR |
| | D82 | NGVGYQPYR | YGVGYQPYR | YGVGYQPYR | YGVGYQPYR | NGVGYQPYR | NGVGYQPYR |

Table S3: 31 epitope peptides were grouped into several pools for vaccine generation and T cell immunity detection.

| | | |
|---|---|---|
| For vaccine generation | pool-v1 | A1+A3+A4+A5+B1+B2+B3+B4 |
| | pool-v2 | B6+C1+C2+C3+D2+D5+D6+D7 |
| | pool-v3 | D11+D12+D13+R3+R4+R5+R6+R8 |
| | pool-v4 | R9+R10+R11+R12+R13+R14+R15 |
| For ELISPOT assay | pool-s1 | A1+A3+A4+A5 |
| | pool-s2 | B1+B2+ B4+B6 |
| | pool-s3 | B3 |
| | pool-s4 | C1+C2+C3 |
| | pool-s5 | D2+D5+D6+D7+ D12+D13 |
| | pool-s6 | D11 |
| | pool-s7 | R5+R6+R8+R11+R12+R14+R15 |
| | pool-s8 | R3+R4+R9+R10+R13 |
| For ICS and ELISA | pool-c1 | A1+A3+A4+A5 |
| | Pool-c2 | B1+B2+B3+B4+B6 |
| | pool-c3 | C1+C2+C3 |
| | pool-c4 | D2+D5+D6+D7+D11+D12+D13 |
| | pool-c5 | R3+ R4+R5+R6+R8+R9+R10+R11+R12 + R13+R14+R15 |

Table S4: Immunization groups, vaccines formula and vaccination scheme

| | |
|---|---|
| PLGA-NPs /peptides Vaccine A | pool-v1: PLGA-NPs/pool-v1(60mg/240μg) mixed with 75μL Poly I:C and 225μLNS, injected into 3 mice, 100μL/mouse, one injection site/mouse<br>pool-v2: PLGA-NPs/pool-v2(60mg/240μg) mixed with 75μL Poly I:C and 225μLNS, injected into 3 mice, 100μL/mouse, one injection site/mouse<br>pool-v3: PLGA-NPs/pool-v3(60mg/240μg) mixed with 75μL Poly I:C and 225μLNS, injected into 3 mice, 100μL/mouse, one injection site/mouse<br>pool-v4: PLGA-NPs/pool-v4(60mg/210μg) mixed with 75μL Poly I:C and 225μLNS, injected into 3 mice, 100μL/mouse, one injection site/mouse |
| R848 /peptides Vaccine B | pool-v1:50μL pool-v1 mixed with 75μL R848 and 175μLNS, injected into 3 mice, 100μL/mouse, one injection site/mouse<br>pool-v2:50μL pool-v2 mixed with 75μL R848 and 175μLNS, injected into 3 mice, 100μL/mouse, one injection site/mouse<br>pool-v3:50μL pool-v3 mixed with 75μL R848 and 175μLNS, injected into 3 mice, 100μL/mouse, one injection site/mouse<br>pool-v4:45μL pool-v4 mixed with 75μL R848 and 180μLNS, injected into 3 mice, 100μL/mouse, one injection site/mouse |
| PolyI:C /peptides Vaccine C | pool-v1: 50μL pool-v1mixed with 75μL Poly I: C and 175μLNS, injected into 3 mice, 100μL/mouse, one injection site/mouse<br>pool-v2: 50μL pool-v2mixed with 75μL Poly I: C and 175μLNS, injected into 3 mice, 100μL/mouse, one injection site/mouse<br>pool-v3:50μL pool-v3mixed with 75μL Poly I: C and 175μLNS, injected into 3 mice, 100μL/mouse, one injection site/mouse<br>pool-v4:45μL pool-v4 mixed with 75μL Poly I: C and 180μLNS, injected into 3 mice, 100μL/mouse, one injection site/mouse |
| Control group | Control mouse 1: 100μL NS/injection site, 4 injection sites<br>Control mouse 2: 80mg empty PLGA-NPs mixed with 400μL NS, 100μL/injection site, 4 injection sites<br>Control mouse3: 100μL NS/injection site, 4 injection sites |

Table S5: Affinity of 31 epitopes with HLA-A0201 molecules on T2 cells

| Protein | Peptide | High affinity (FI>1) | Intermediate affinity (0.5＜FI ≤1) | Low or no binding (FI ≤ 0.5) | FI |
|---|---|---|---|---|---|
| E | A1 | | + | | 0.65 |
| | A3 | | | + | 0.39 |
| | A4 | | + | | 0.82 |
| | A5 | + | | | 1.65 |
| | B1 | + | | | 2.28 |
| M | B2 | | + | | 0.94 |
| | B3 | + | | | 3.34 |
| | B4 | | + | | 0.51 |
| | B6 | + | | | 1.73 |
| N | C1 | + | | | 1.26 |
| | C2 | | | + | 0.21 |
| | C3 | | | + | 0.43 |
| S | D2 | + | | | 1.12 |
| | D5 | + | | | 2.59 |
| | D6 | + | | | 3.65 |
| | D7 | | + | | 0.89 |
| | D11 | + | | | 2.11 |
| | D12 | | | + | 0.35 |
| | D13 | | | + | 0.24 |
| RdRp | R3 | + | | | 2.91 |
| | R4 | + | | | 1.98 |
| | R5 | + | | | 4.00 |
| | R6 | + | | | 2.48 |
| | R8 | + | | | 2.11 |
| | R9 | + | | | 3.65 |
| | R10 | | + | | 0.58 |
| | R11 | + | | | 1.95 |
| | R12 | | + | | 0.79 |
| | R13 | | | + | 0.41 |
| | R14 | + | | | 2.21 |
| | R15 | + | | | 3.67 |
| Total | 31 | 18 | 7 | 6 | |

Figure S1:

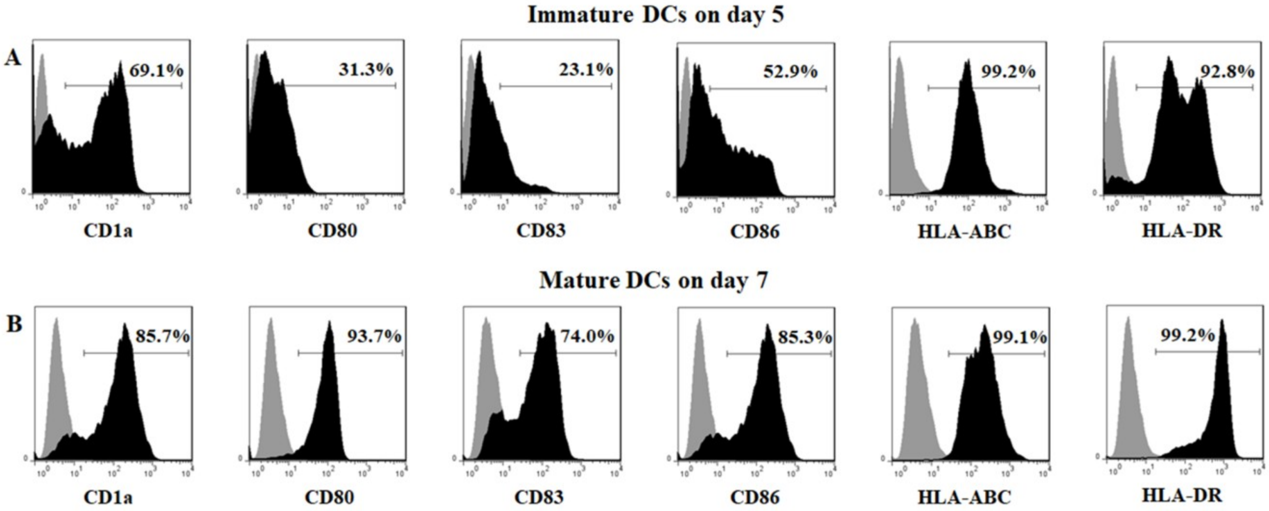

Figure S1: Generation of mature mDC from adherent monocytes. PBMCs from healthy donors were seeded into culture flask and the monocytes adhered for 2 h as described in the Methods section. After washing out the non-adherent cells in both systems, the cells were cultured for 5 days with 1,000 IU/mL GM-CSF and 500 IU/mL IL-4. Then the immature DCs were matured with 1μg/mL LPS for another 48 hours. Immature DCs and mature DCs were stained for CD1a, CD80, CD83, CD86, HLA-ABC and HLA-DR. The unstained and stained populations in the histograms are shown in grey and black, respectively. (A) Phenotype of immature DCs on day 5. (B) Phenotype of mature DCs on day 7.

Figure S2:

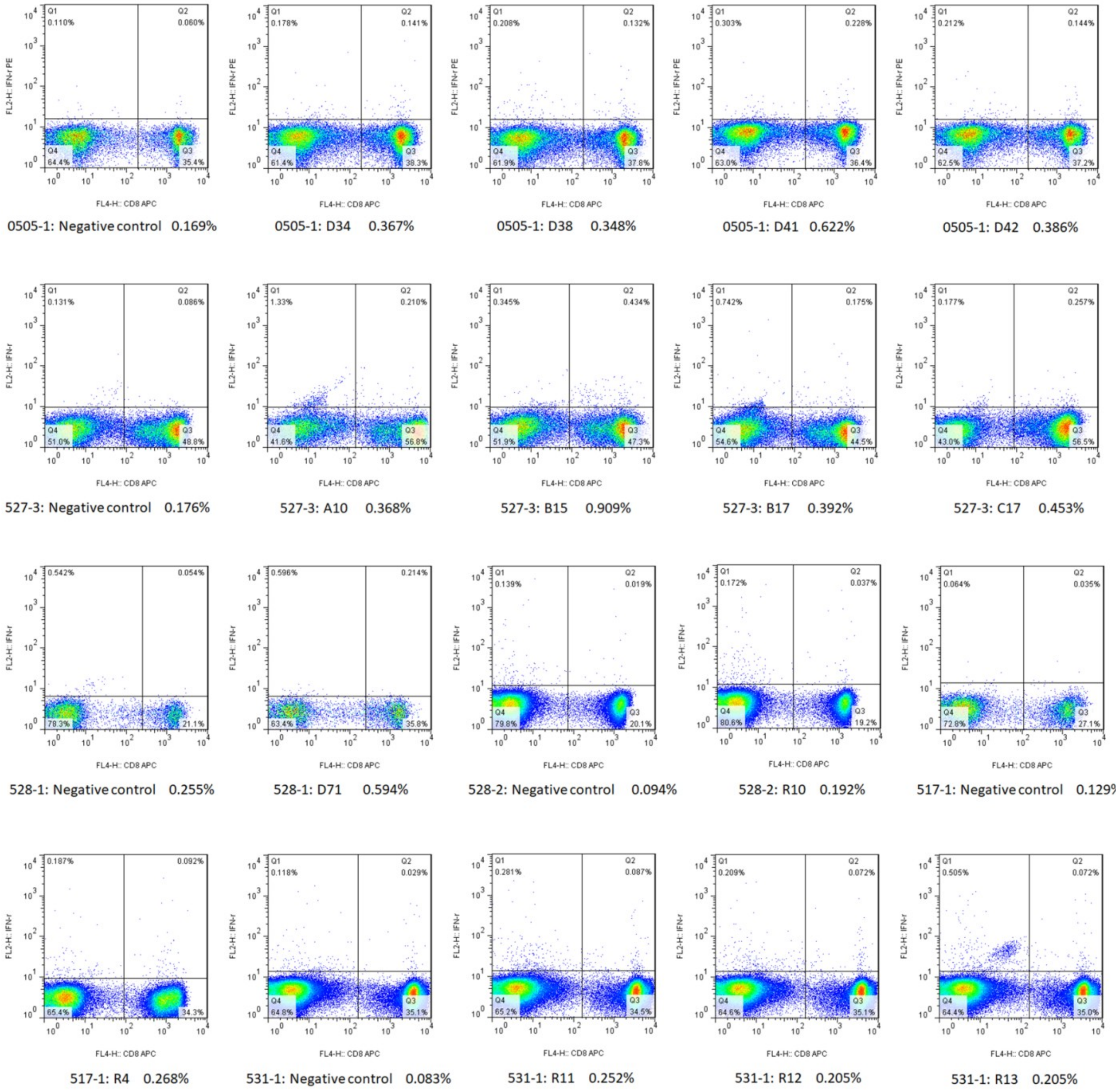
0505-1: Negative control 0.169%
0505-1: D34 0.367%
0505-1: D38 0.348%
0505-1: D41 0.622%
0505-1: D42 0.386%
527-3: Negative control 0.176%
527-3: A10 0.368%
527-3: B15 0.909%
527-3: B17 0.392%
527-3: C17 0.453%
528-1: Negative control 0.255%
528-1: D71 0.594%
528-2: Negative control 0.094%
528-2: R10 0.192%
517-1: Negative control 0.129
517-1: R4 0.268%
531-1: Negative control 0.083%
531-1: R11 0.252%
531-1: R12 0.205%
531-1: R13 0.205%

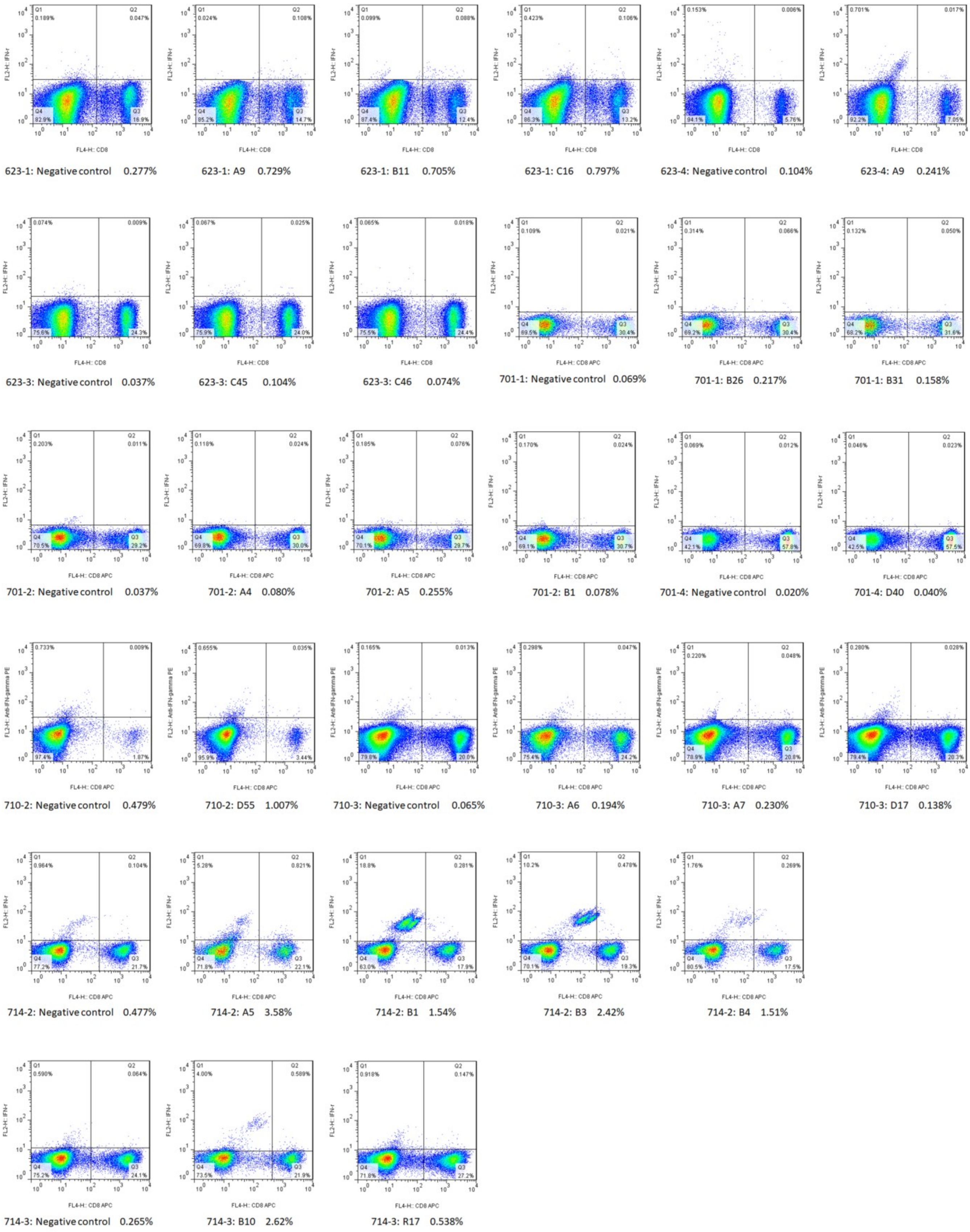
623-1: Negative control 0.277%
623-1: A9 0.729%
623-1: B11 0.705%
623-1: C16 0.797%
623-4: Negative control 0.104%
623-4: A9 0.241%
623-3: Negative control 0.037%
623-3: C45 0.104%
623-3: C46 0.074%
701-1: Negative control 0.069%
701-1: B26 0.217%
701-1: B31 0.158%
701-2: Negative control 0.037%
701-2: A4 0.080%
701-2: A5 0.255%
701-2: B1 0.078%
701-4: Negative control 0.020%
701-4: D40 0.040%
710-2: Negative control 0.479%
710-2: D55 1.007%
710-3: Negative control 0.065%
710-3: A6 0.194%
710-3: A7 0.230%
710-3: D17 0.138%
714-2: Negative control 0.477%
714-2: A5 3.58%
714-2: B1 1.54%
714-2: B3 2.42%
714-2: B4 1.51%
714-3: Negative control 0.265%
714-3: B10 2.62%
714-3: R17 0.538%

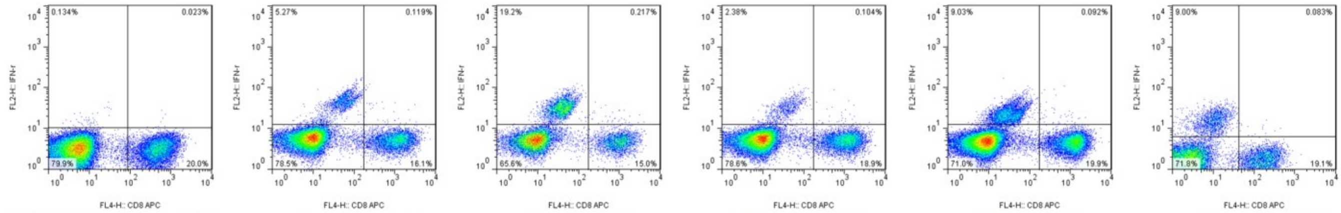

714-4: Negative control 0.114% | 714-4: A20 0.734% | 714-4: A21 1.43% | 714-4: A22 0.546% | 714-4: A23 0.46% | 714-4: B26 0.43%

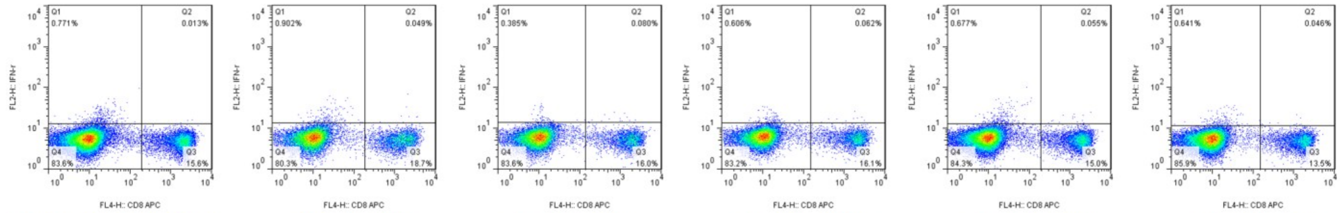

715-2: Negative control 0.083% | 715-2: B28 0.261 | 715-2: B29 0.498% | 715-2: B30 0.384% | 715-2: R47 0.365% | 715-2: R48 0.34%

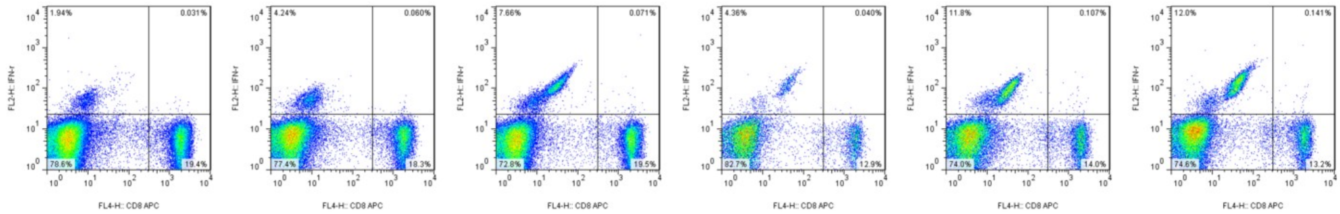

720-2: Negative control 0.160% | 720-2: R11 0.327% | 720-2: R13 0.363% | 720-3: Negative control 0.309% | 720-3: D30 0.758% | 720-3: B6 1.057%

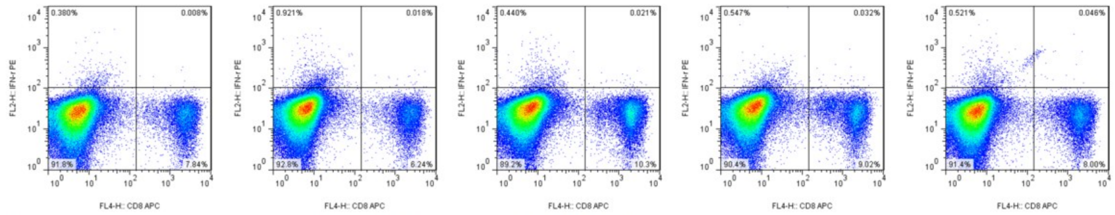

721-2: Negative control 0.102% | 721-2: D62 0.288% | 721-2: D64 0.203% | 721-2: D52 0.354% | 721-2: D53 0.572%

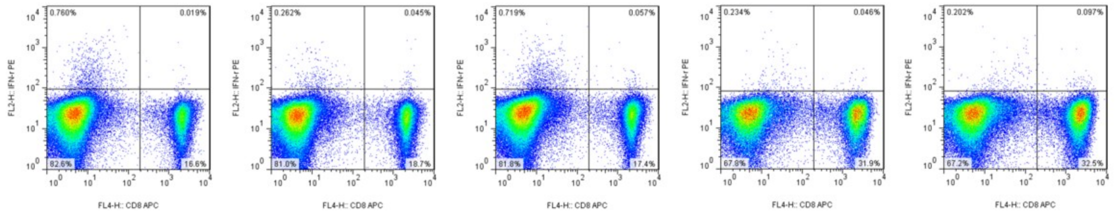

721-4: Negative control 0.114% | 721-4: B37 0.240% | 721-4: B38 0.327% | 722-3: Negative control 0.144% | 722-3: D6 0.298%

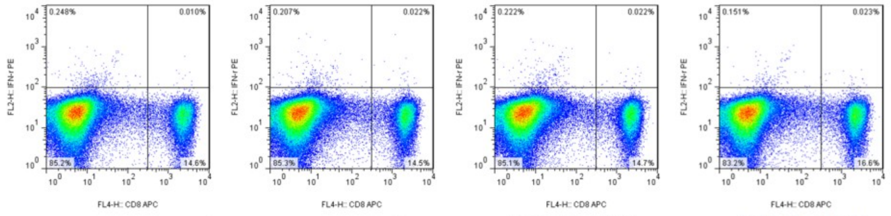

722-2: Negative control 0.068% | 722-2: R34 0.151% | 722-2: A18 0.149% | 722-2: A19 0.138%

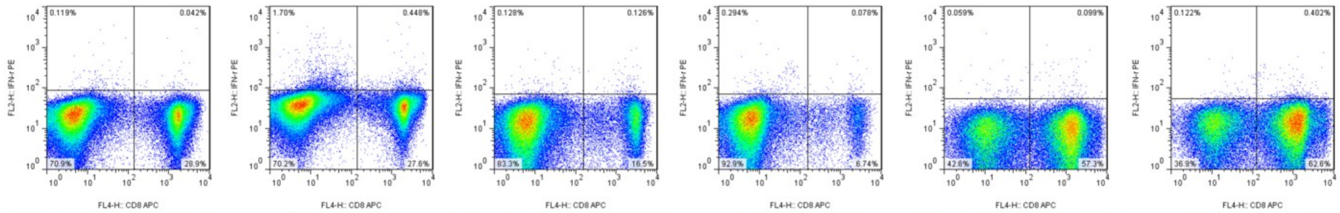

722-4: Negative control 0.145% | 722-4: D13 1.597% | 724-2: Negative control 0.566% | 724-2: B34 1.144% | 724-4: Negative control 0.172% | 724-4: C3 0.638%

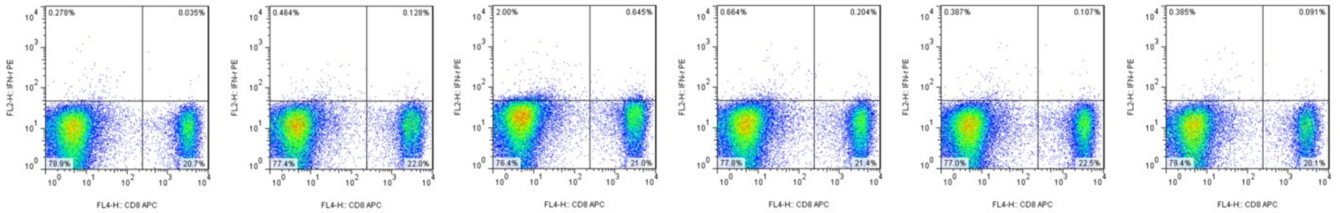

724-3: Negative control 0.169% | 724-3: D50 0.578% | 724-3: C49 2.980% | 724-3: D46 0.944% | 724-3: D47 0.473% | 724-3: D48 0.451%

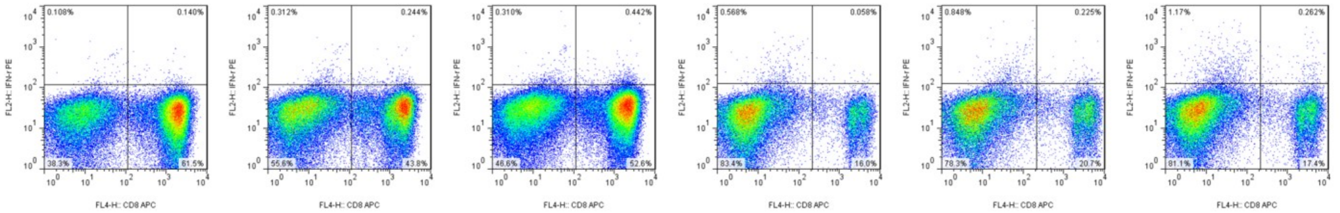

727-1: Negative control 0.227% | 727-1: C35 0.554% | 727-1: D56 0.833% | 727-3: Negative control 0.361% | 727-3: C47 1.075% | 727-3: D76 1.483%

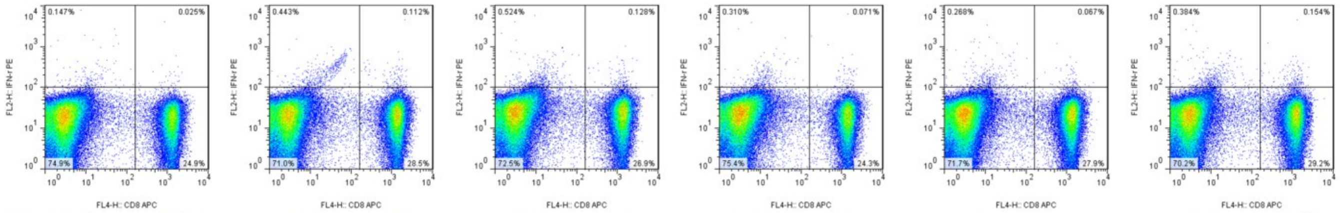

727-2: Negative control 0.100% | 727-2: A26 0.391% | 727-2: B36 0.474% | 727-2: B37 0.291% | 727-2: B40 0.240% | 727-2: B41 0.525%

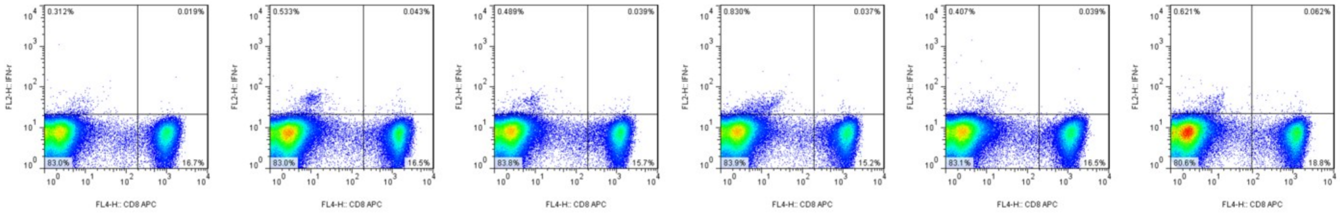

731-4: Negative control 0.114% | 731-4: B2 0.26% | 731-4: R5 0.248% | 731-4: R8 0.243% | 731-4: R9 0.236% | 731-4: R12 0.329%

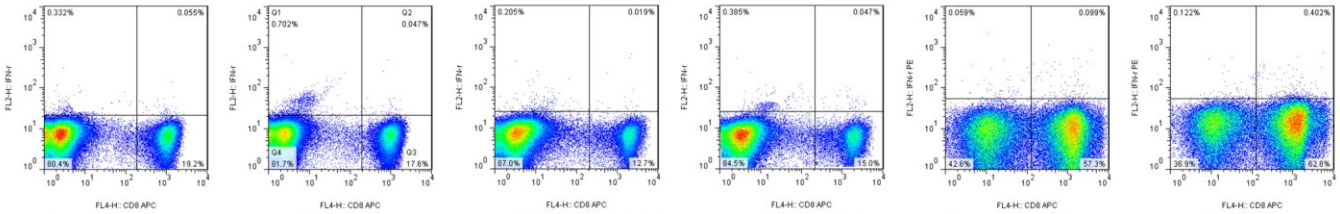

731-4: R13 0.286% | 731-4: R14 0.266% | 731-1: Negative control 0.149% | 731-1: R23 0.312% | 724-4: Negative control 0.172% | 724-4: C3 0.638%

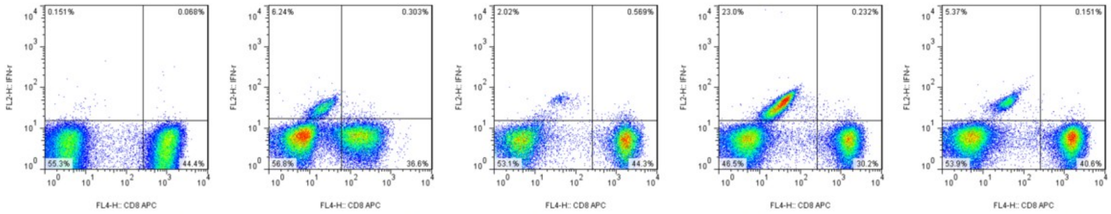

820-3: Negative control 0.153%　820-3: A4 0.821%　820-3: A5 1.268%　820-3: B1 0.762%　820-3: B29 0.371%

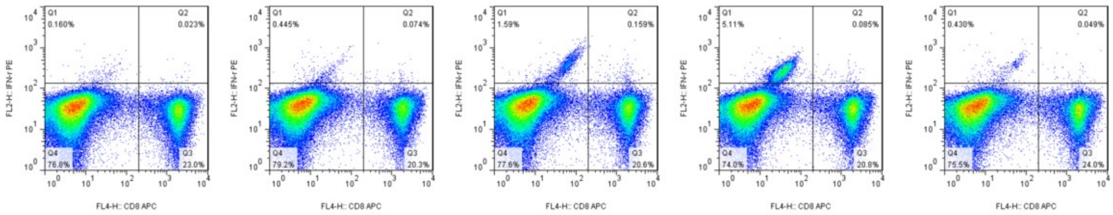

821-1: Negative control 0.100%　821-1: A6 0.363%　821-1: A9 0.766%　821-1: B11 0.407%　821-1: B12 0.204%

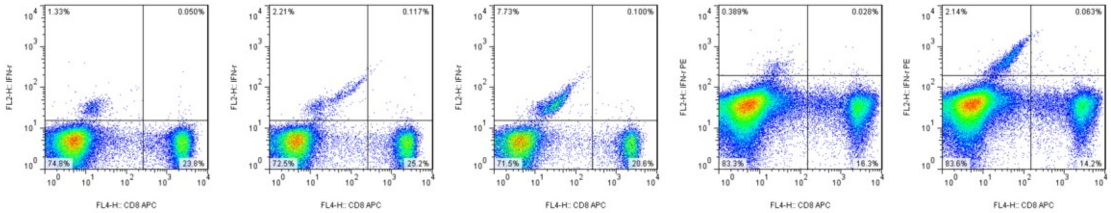

820-2: Negative control 0.21%　820-2: B10 0.462%　820-2: A1 0.483%　821-2: Negative control 0.171%　821-2: A12 0.442%

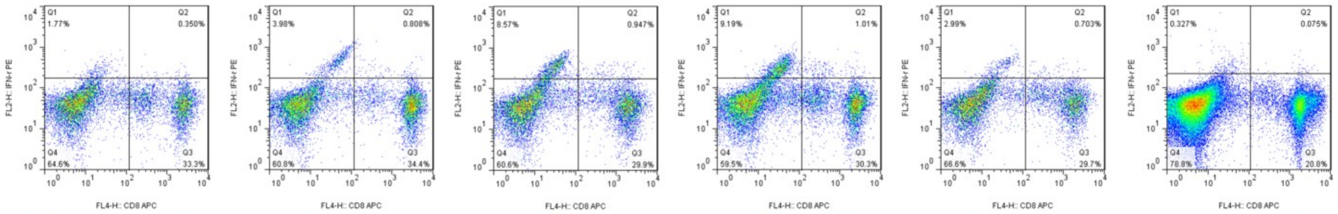

822-1: Negative control 1.040%　822-1: A1 2.295%　822-1: A5 3.070%　822-1: B1 3.226%　822-1: D2 2.312%　822-2: Negative control 0.359%

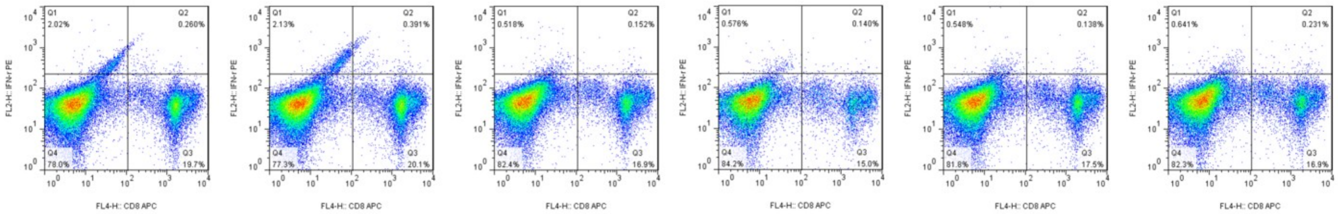

822-2: A12 1.303%　822-2: B23 1.908%　822-2: B35 0.891%　822-2: D72 0.925%　822-2: D48 0.782%　822-2: D53 1.348%

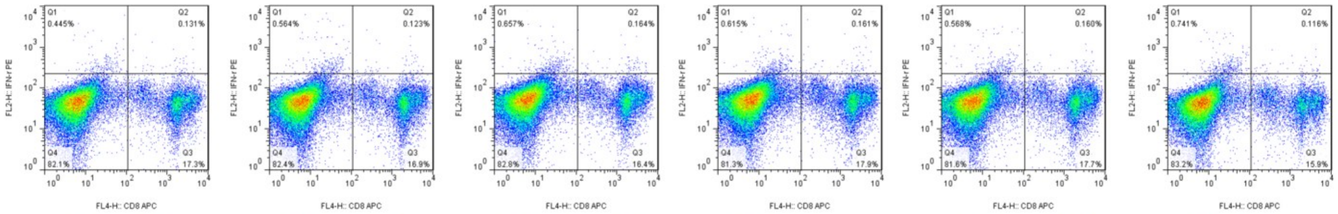

822-2: D77 0.752%　822-2: D78 0.723%　822-2: D79 0.990%　822-2: D80 0.891%　822-2: D81 0.896%　822-2: D82 0.724%

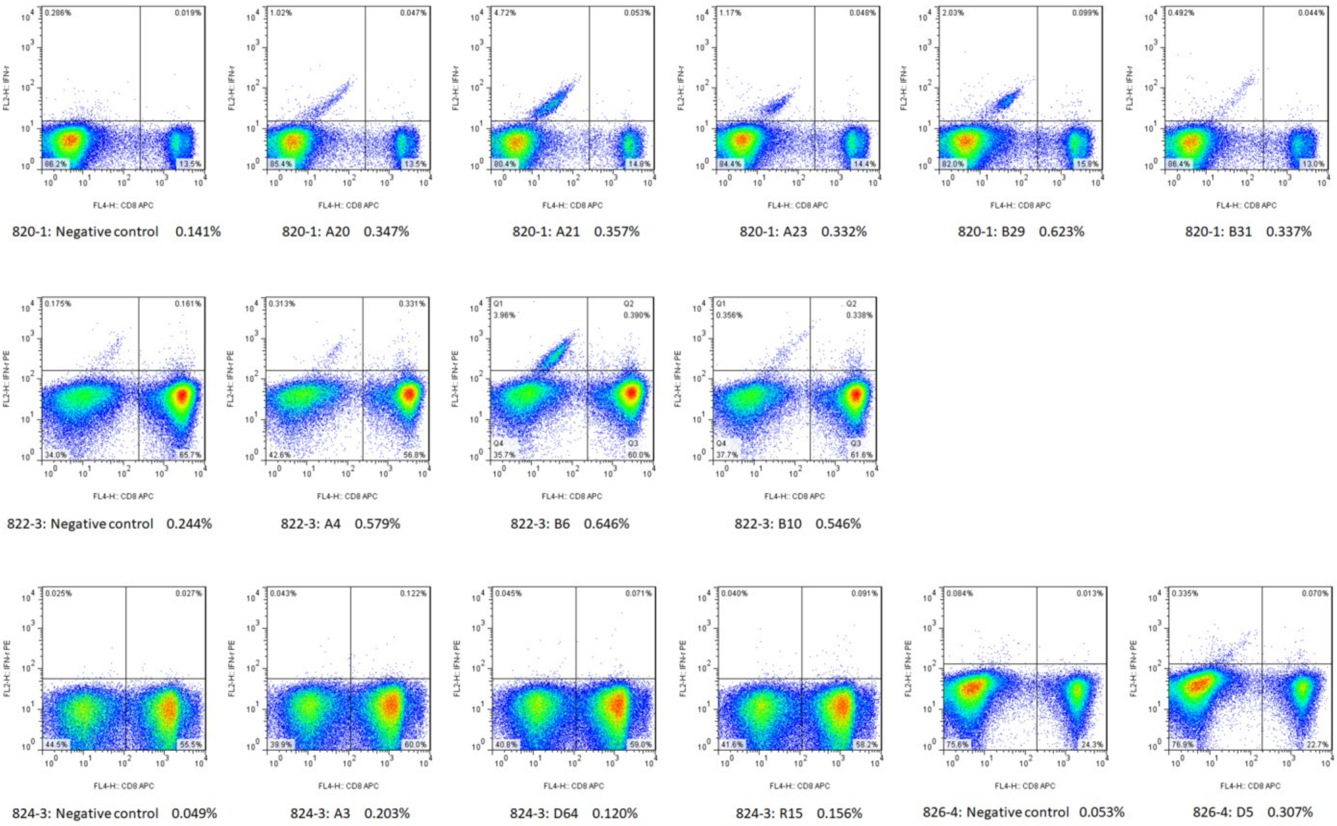

Figure S2: Immunogenicity of 120 candidate epitopes was validated by DC-peptide-PBL co-culture experiments (IFN-γ ICS flow plots). DCs were induced for 7 days from healthy donor's PBMCs, and then coincubated with candidate epitope peptides and autologous PBLs for 14 days. Cells were harvested and stimulated by corresponding candidate peptides for another 16 hours followed by IFN-γ ICS. The presented are flow plot spots for each positive epitope peptide. The data in horizontal coordinates mean the frequency of IFN-$\gamma^{+}$ T cells in $CD3^{+}/CD8^{+}$ T cell population.

Figure S3:

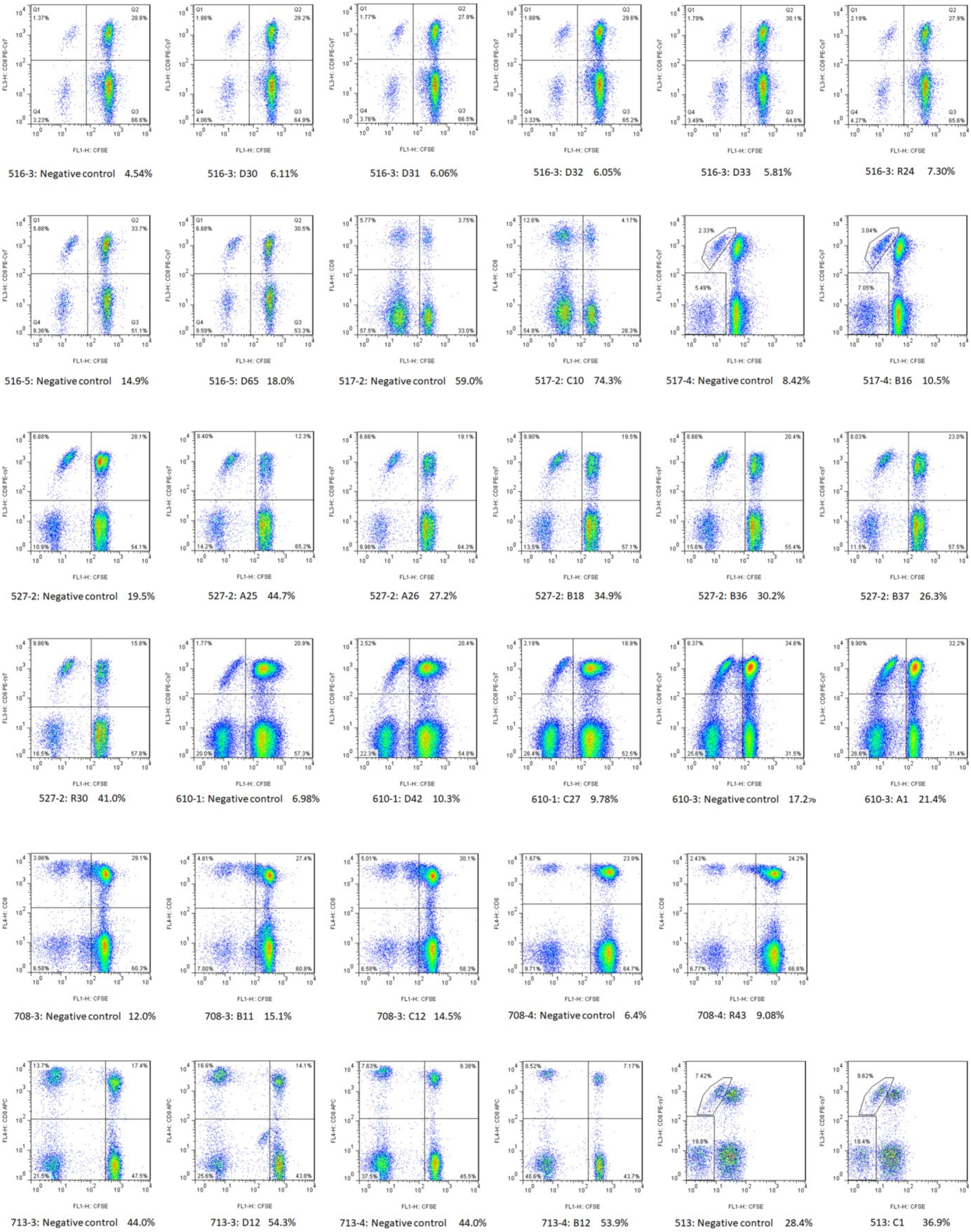

516-3: Negative control 4.54%
516-3: D30 6.11%
516-3: D31 6.06%
516-3: D32 6.05%
516-3: D33 5.81%
516-3: R24 7.30%
516-5: Negative control 14.9%
516-5: D65 18.0%
517-2: Negative control 59.0%
517-2: C10 74.3%
517-4: Negative control 8.42%
517-4: B16 10.5%
527-2: Negative control 19.5%
527-2: A25 44.7%
527-2: A26 27.2%
527-2: B18 34.9%
527-2: B36 30.2%
527-2: B37 26.3%
527-2: R30 41.0%
610-1: Negative control 6.98%
610-1: D42 10.3%
610-1: C27 9.78%
610-3: Negative control 17.2%
610-3: A1 21.4%
708-3: Negative control 12.0%
708-3: B11 15.1%
708-3: C12 14.5%
708-4: Negative control 6.4%
708-4: R43 9.08%
713-3: Negative control 44.0%
713-3: D12 54.3%
713-4: Negative control 44.0%
713-4: B12 53.9%
513: Negative control 28.4%
513: C1 36.9%

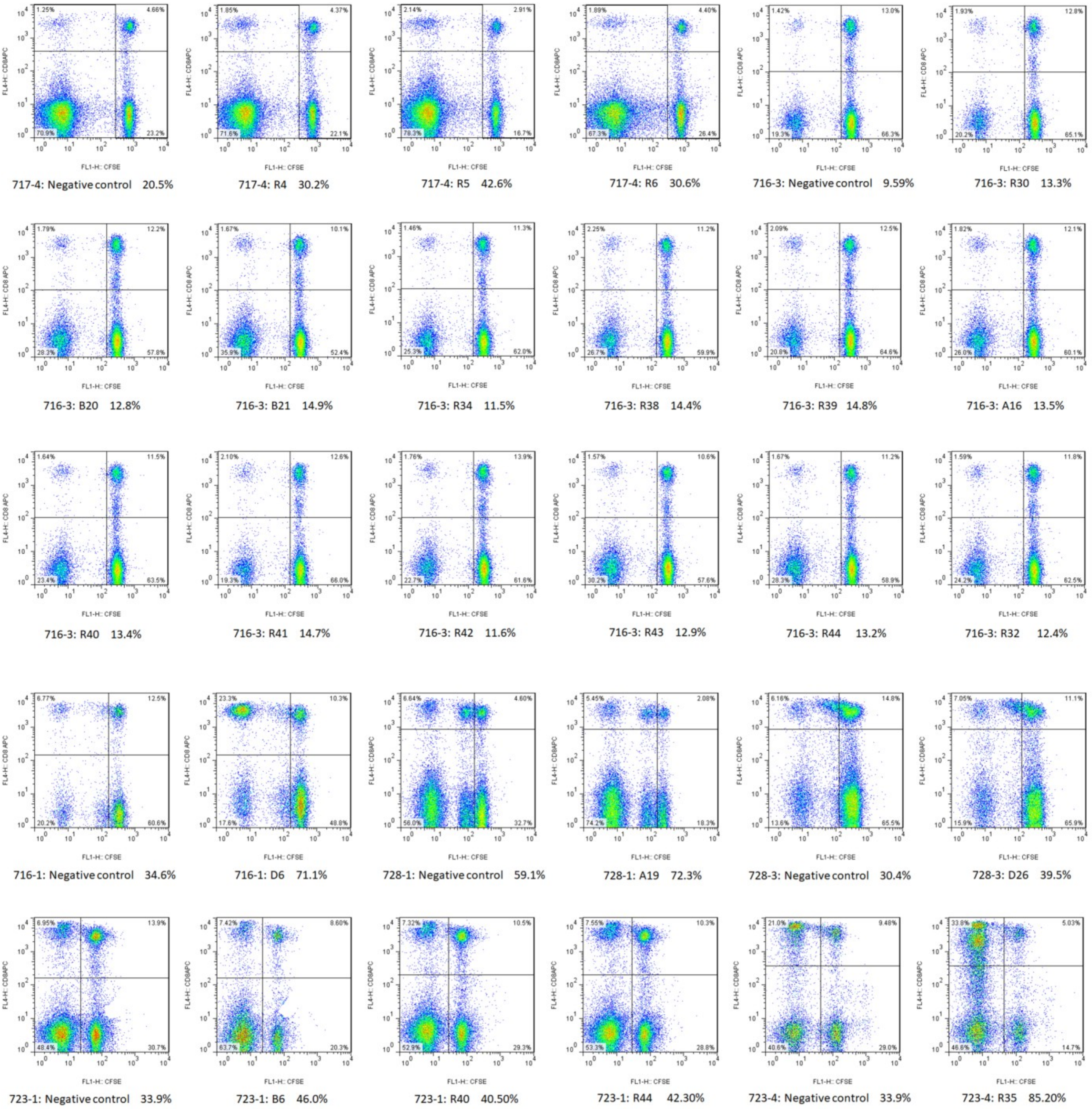

Figure S3: Immunogenicity of 120 candidate epitopes was validated by DC-peptide-PBL co-culture experiments (CFSE staining flow plots). DCs were coincubated with candidate epitope peptides and CFSE-prelabeled PBLs for 14 days. Cells were then analyzed by flow cytometry. The presented are flow plot spots for each positive epitope peptide. The data in horizontal coordinates mean the proliferation percentage of $CD8^+$ T cells in $CD3^+/CD8^+$ T cell population.

Figure S4:

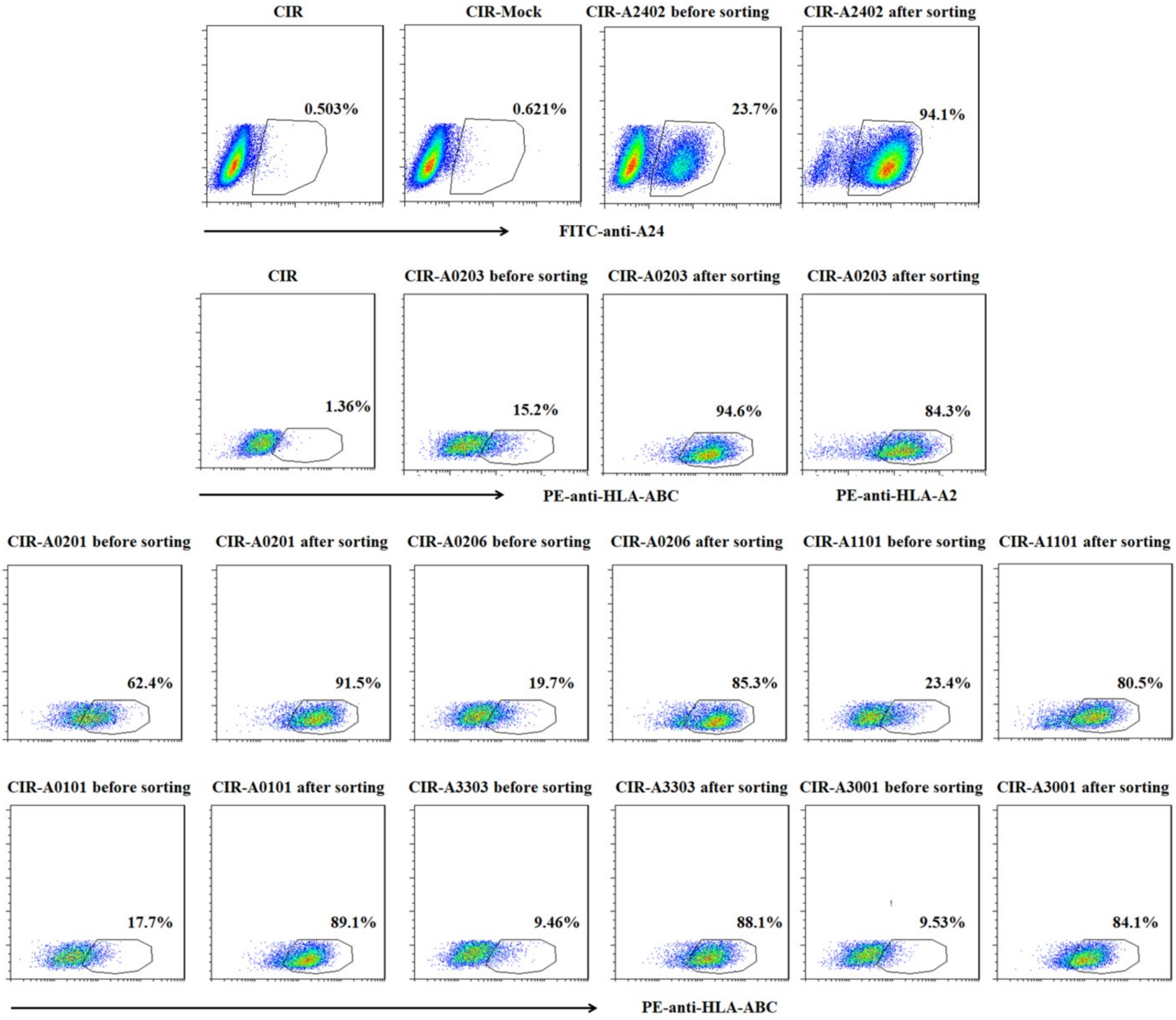

**Figure S4: Eight kinds of HMy2.CIR cell lines expressing one indicated HLA-A allotype.** The transfected HMy2.CIR cell lines expressing HLA-A2402, A0203, A0201, A0206, A1101, A3303, A0101, or A3001 were generated, respectively, then sorted by flow cytometry followed by pure culture and sequencing analyses. The purity of CIR-A2402 cells was 94.1% after being sorted with FITC-anti-A24 staining. The purity of CIR-A0203 cell was 84.3% after being sorted with PE-anti-HLA-ABC (W6/32) or PE-anti-HLA-A2.1 staining. The purities of CIR-A0201/0206/1101/3303/0101/3001 were all more than 80% after being sorted with PE-anti-HLA-ABC staining.

Figure S5:

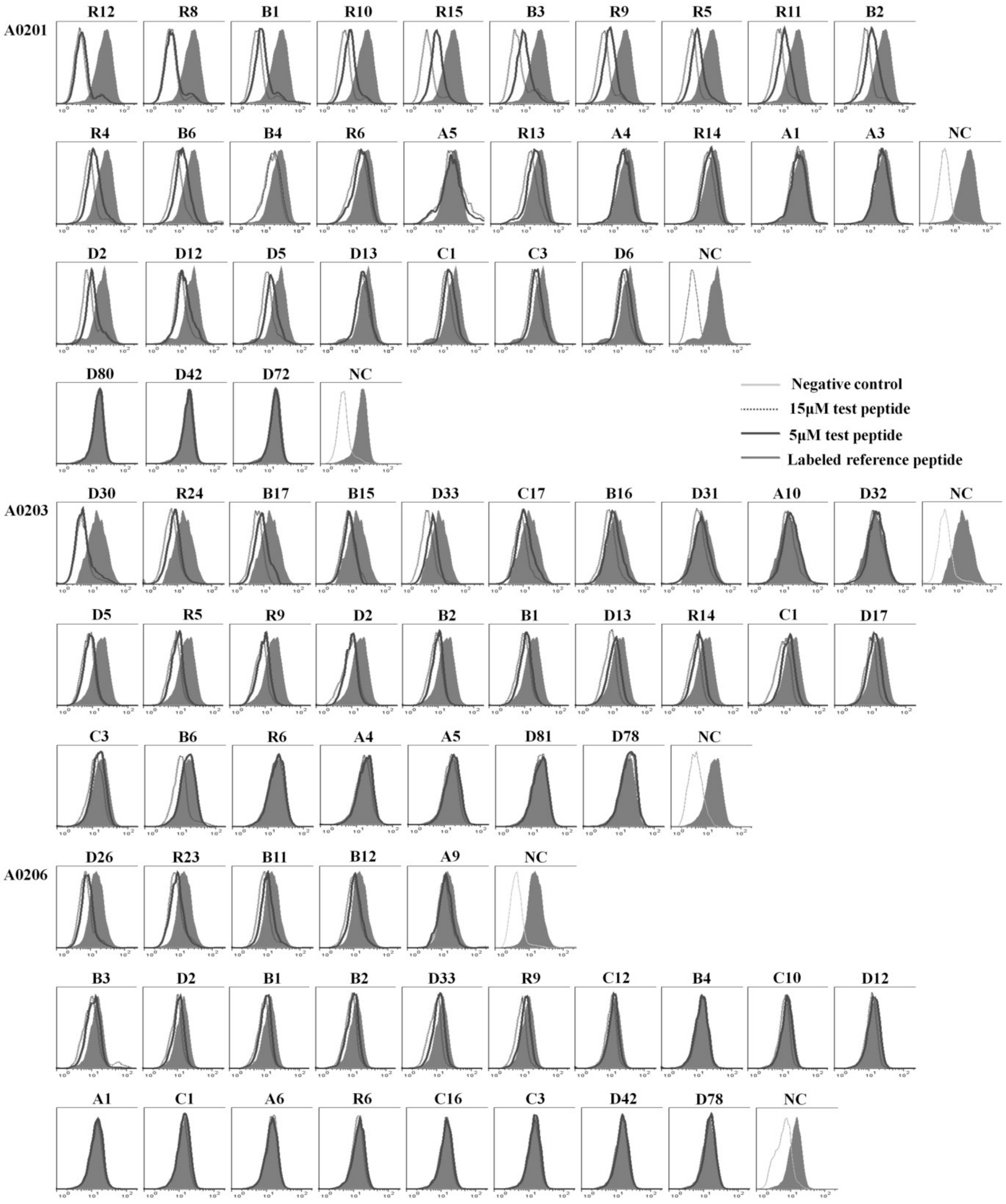
A0201
R12 R8 B1 R10 R15 B3 R9 R5 R11 B2
R4 B6 B4 R6 A5 R13 A4 R14 A1 A3 NC
D2 D12 D5 D13 C1 C3 D6 NC
D80 D42 D72 NC
Negative control
15μM test peptide
5μM test peptide
Labeled reference peptide
A0203
D30 R24 B17 B15 D33 C17 B16 D31 A10 D32 NC
D5 R5 R9 D2 B2 B1 D13 R14 C1 D17
C3 B6 R6 A4 A5 D81 D78 NC
A0206
D26 R23 B11 B12 A9 NC
B3 D2 B1 B2 D33 R9 C12 B4 C10 D12
A1 C1 A6 R6 C16 C3 D42 D78 NC

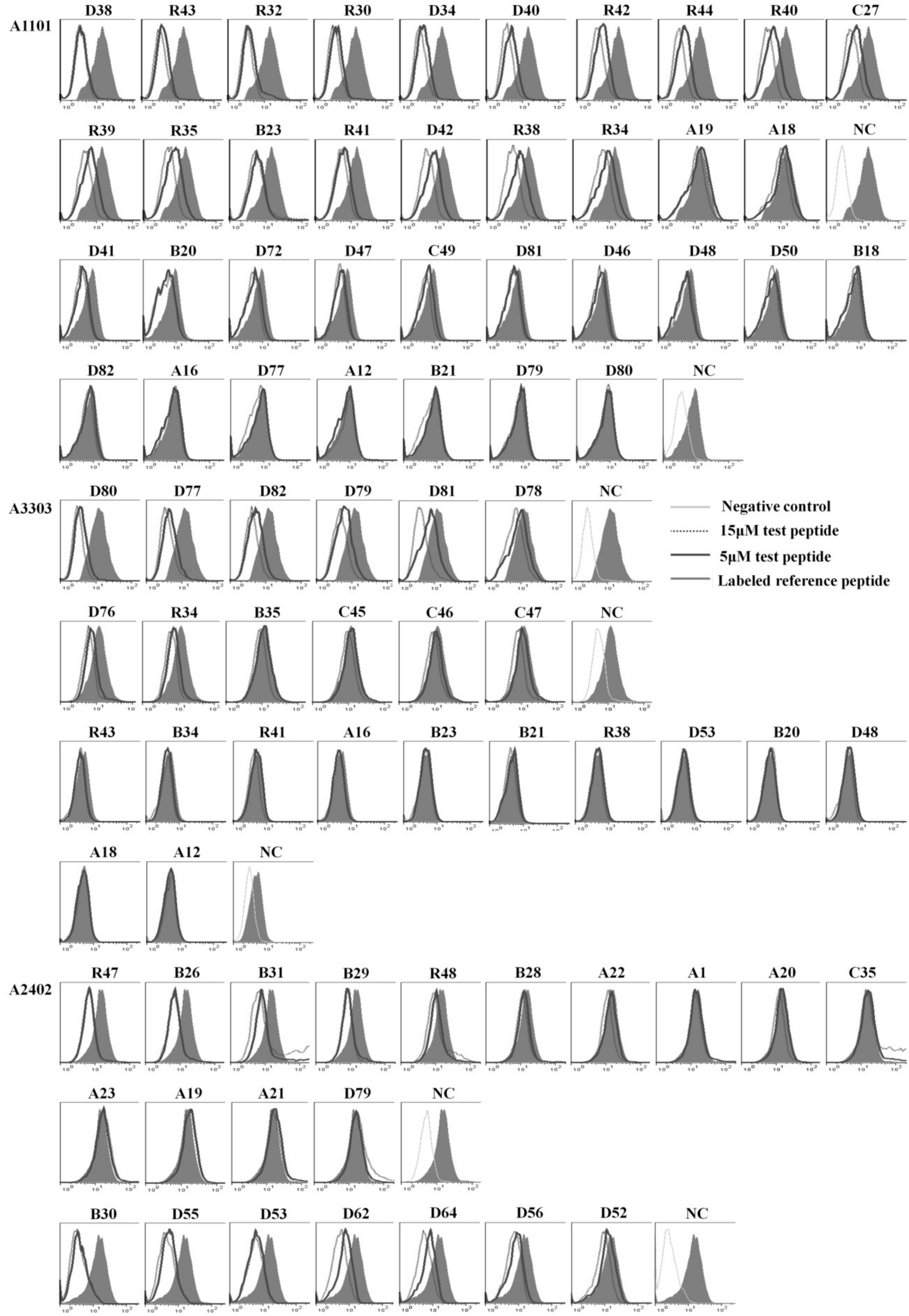
A1101
D38
R43
R32
R30
D34
D40
R42
R44
R40
C27
R39
R35
B23
R41
D42
R38
R34
A19
A18
NC
D41
B20
D72
D47
C49
D81
D46
D48
D50
B18
D82
A16
D77
A12
B21
D79
D80
NC
A3303
D80
D77
D82
D79
D81
D78
NC
Negative control
15μM test peptide
5μM test peptide
Labeled reference peptide
D76
R34
B35
C45
C46
C47
NC
R43
B34
R41
A16
B23
B21
R38
D53
B20
D48
A18
A12
NC
A2402
R47
B26
B31
B29
R48
B28
A22
A1
A20
C35
A23
A19
A21
D79
NC
B30
D55
D53
D62
D64
D56
D52
NC

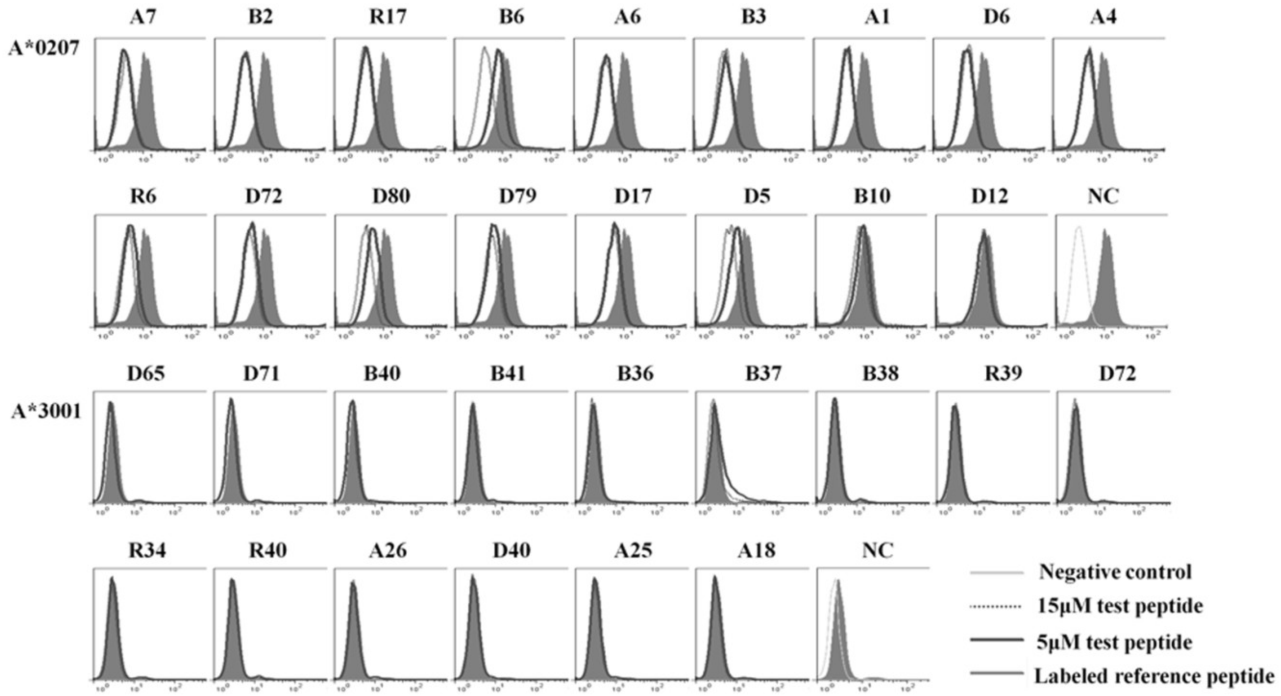

**Figure S5: Binding affinity of 120 validated epitopes with HLA-A allotypes as defined by HLA-A molecule competitive binding experiments.** A series of unlabeled epitope peptides of SRAS-CoV-2 were coincubated, at 5μM and 15μM respectively, with fluorescent-labeled reference peptides and CIR cell lines expressing the corresponding HLA-A molecules for 24 hours. Then the competitively binding inhibition (%) of the epitope peptide at 5μM and 15μM was calculated by measuring the CIR cells fluorescence strength. Shown are the histograms of two concentrations (5μM and 15μM). Black solid line was the histogram of 5μM test peptide; dotted line was the histogram of 15μM, test peptide; black filled line was the maximal fluorescence (FITC-labeled reference peptide without competitive peptides) while the lightest gray line was the negative control (background fluorescence with 1640 alone).

Figure S6:

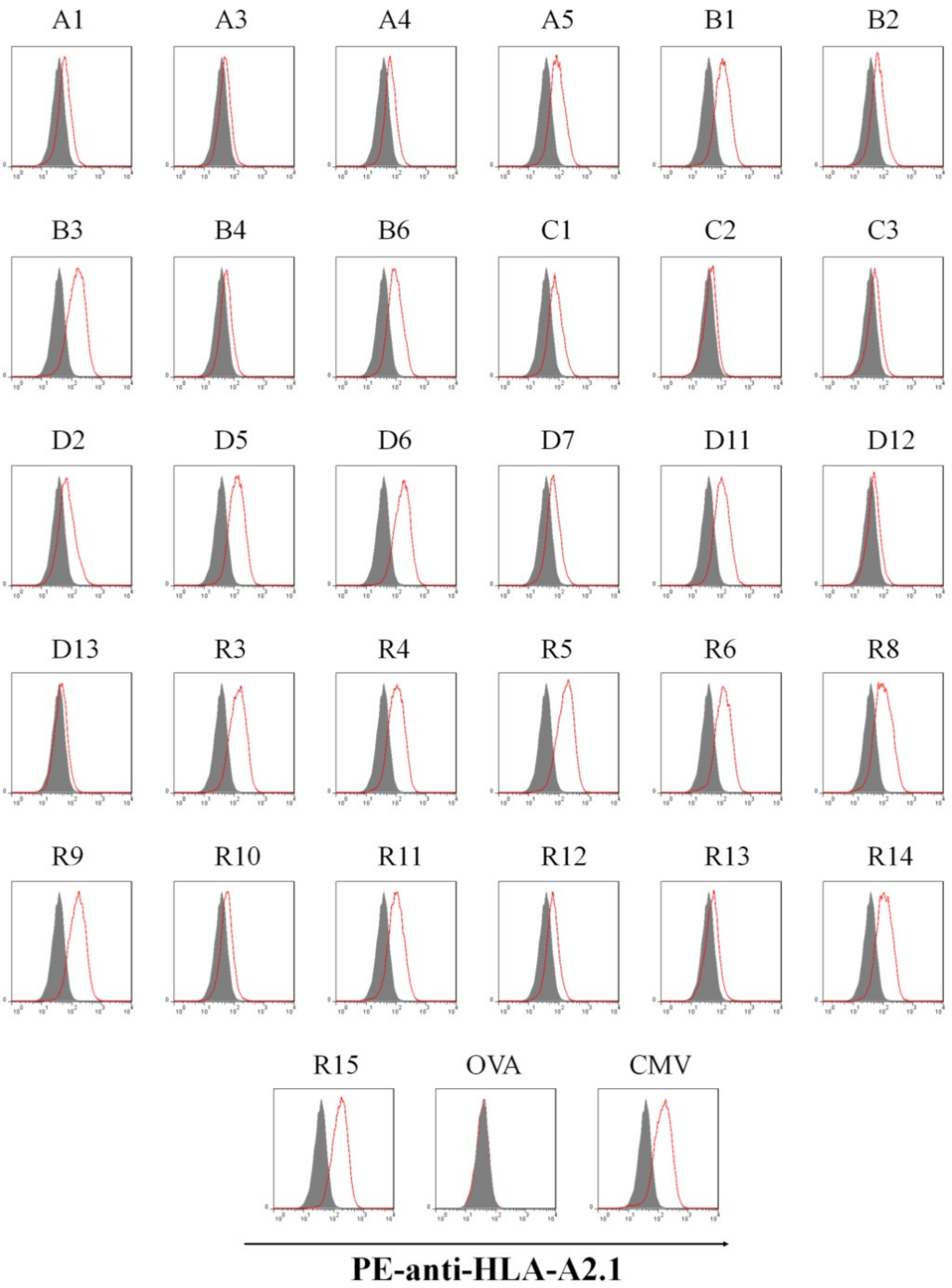

Figure S6: Affinity of 31 epitope peptides with HLA-A0201 molecule as detected by T2 cell binding assay. T2 cells were incubated with single peptide of the 31 epitopes restricted by HLA-A2 molecules, or with CMVpp65$_{495\text{-}503}$ peptide as positive control, OVA$_{257\text{-}264}$ peptide as negative control, or no peptide and β2-microglobulin for 16h, and then followed by PE-anti-HLA-A2.1 antibody staining to test the up-regulation of HLA-A0201 molecules onto T2 cells. Red solid line was the histogram of indicated peptides while background fluorescence was the histogram of T2 cells alone without peptide.

Figure S7:

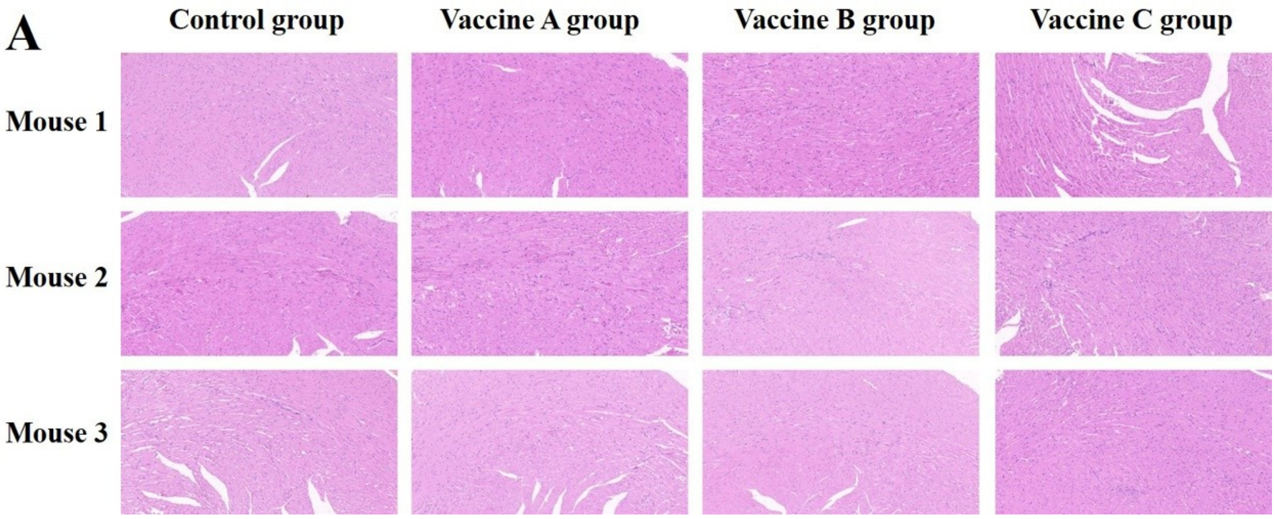
A
Control group
Vaccine A group
Vaccine B group
Vaccine C group
Mouse 1
Mouse 2
Mouse 3

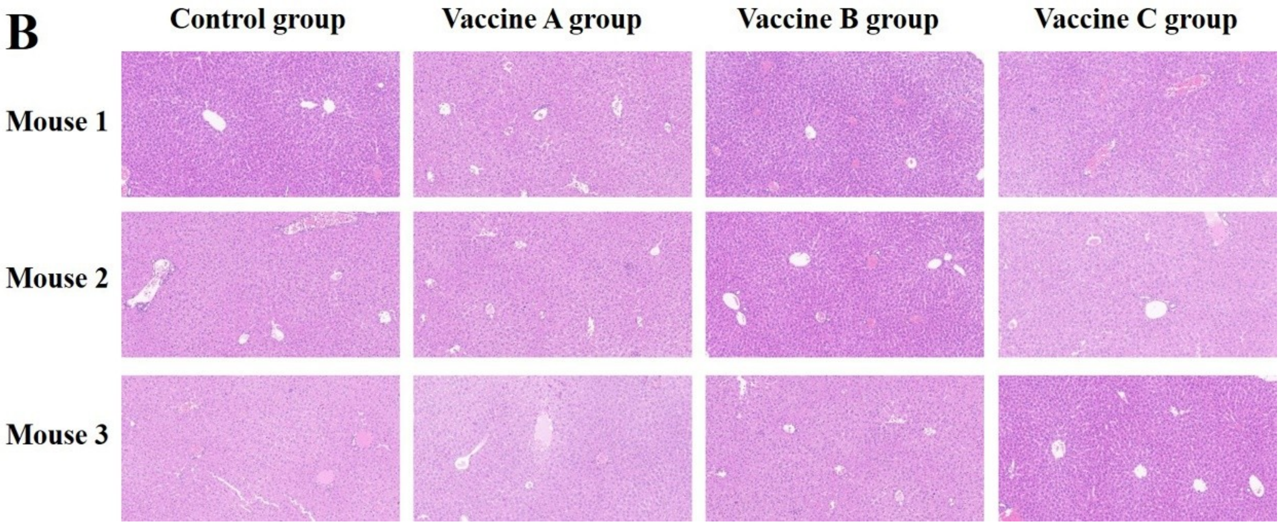
B
Control group
Vaccine A group
Vaccine B group
Vaccine C group
Mouse 1
Mouse 2
Mouse 3

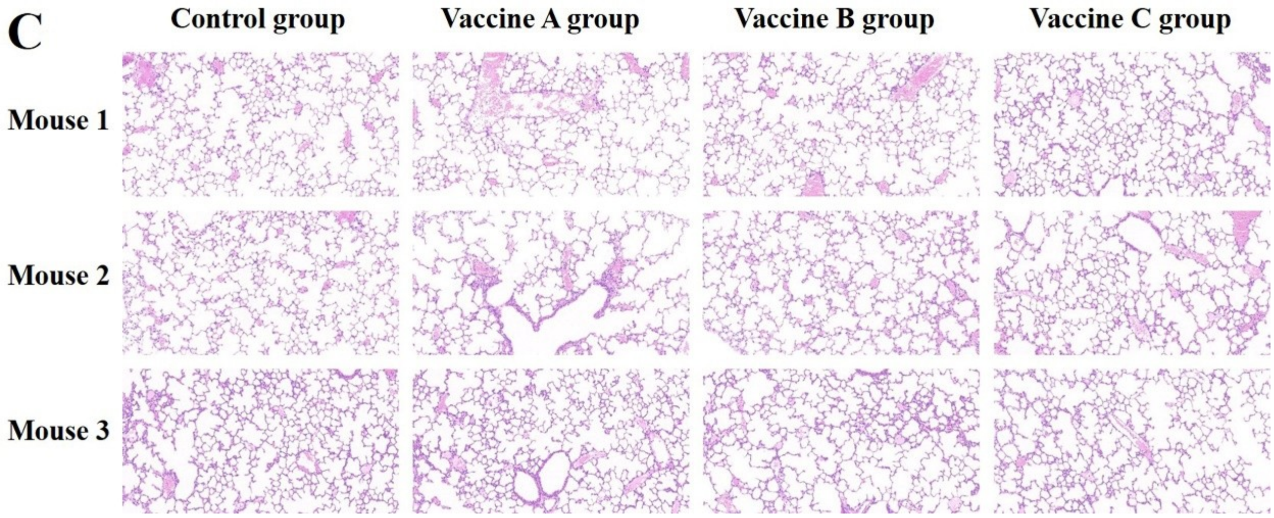

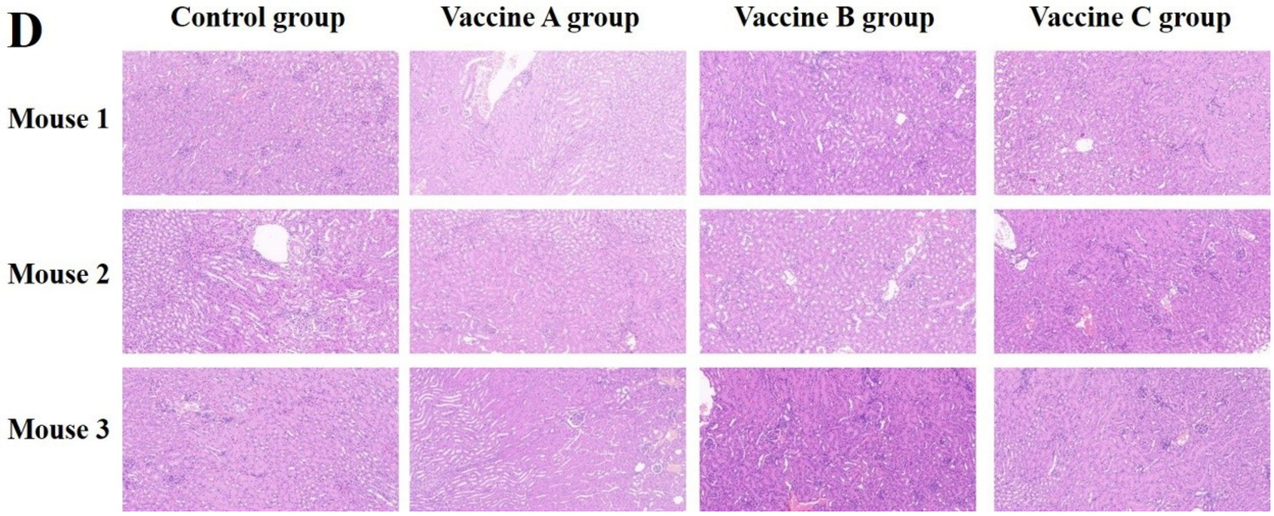

Figure S7: The peptides based vaccines have no visible toxicity on the organs. Seven days after the last booster, all mice were executed. Heart, liver, lung and kidney were taken out, immersed and were finally stained with Hematoxylin-Eosin. The tissue slides were then scanned and read with software called case viewer. No obvious pathological damage was found in all organs in all groups. The representative HE staining of heart, liver, lung and kidney in each mouse from four groups were exhibited in (A), (B), (C), and (D), respectively.
